# Chemoproteomics identifies proteoform-selective caspase-2 inhibitors

**DOI:** 10.1101/2023.10.25.563785

**Authors:** José O. Castellón, Samuel Ofori, Ernest Armenta, Nikolas Burton, Lisa M. Boatner, Evan E. Takayoshi, Marina Faragalla, Annie Zhou, Ky Tran, Jeremy Shek, Tianyang Yan, Heta S. Desai, Keriann M. Backus

## Abstract

Caspases are a highly conserved family of cysteine-aspartyl proteases known for their essential roles in regulating apoptosis, inflammation, cell differentiation, and proliferation. Complementary to genetic approaches, small-molecule probes have emerged as useful tools for modulating caspase activity. However, due to the high sequence and structure homology of all twelve human caspases, achieving selectivity remains a central challenge for caspase-directed small-molecule inhibitor development efforts. Here, using mass spectrometry-based chemoproteomics, we first identify a highly reactive non-catalytic cysteine that is unique to caspase-2. By combining both gel-based activity-based protein profiling (ABPP) and a *tobacco etch virus* (TEV) protease activation assay, we then identify covalent lead compounds that react preferentially with this cysteine and afford a complete blockade of caspase-2 activity. Inhibitory activity is restricted to the zymogen or precursor form of monomeric caspase-2. Focused analogue synthesis combined with chemoproteomic target engagement analysis in cellular lysates and in cells yielded both pan-caspase reactive molecules and caspase-2 selective lead compounds together with a structurally matched inactive control. Application of this focused set of tool compounds to stratify caspase contributions to initiation of intrinsic apoptosis, supports compensatory caspase-9 activity in the context of caspase-2 inactivation. More broadly, our study highlights future opportunities for the development of proteoform-selective caspase inhibitors that target non-conserved and non-catalytic cysteine residues.

## INTRODUCTION

Programmed cell death or apoptosis is a tightly regulated biological process required for the removal of irreversibly damaged or unwanted cells. Caspases are cysteine aspartate proteases responsible for the initiation and execution of programmed cell death^1–3^. Caspase-mediated proteolysis has also been implicated in a number of non-apoptotic processes, including cellular activation^4–5^, differentiation^6–8^, cell proliferation^9^, immune response^4, 10–12^, cell cycle regulation^9, 13^, and inflammation^10, 14^. Human cancers^9, 15–17^, neurodegenerative diseases^18^ and monogenic disorders^4, 12^ have all been linked to aberrant caspase activity. Motivated by these numerous important and diverse functions, there is ongoing interest in the functional stratification of individual caspases.

Delineating the unique and overlapping functions of caspase family members has been hindered by their high sequence and structural homology^19–21^ as well as compensatory functional activities—12 caspases are encoded by the human genome^22^. While genetic tools, including CRISPR-Cas9, have been successfully applied to the study of individual family members^23^, such approaches are complicated by the compensatory expression of closely related caspase homologues^24^.

Chemical probes are useful tools for studying protein function; complementary to genetic approaches, chemical probes offer the added advantages of producing graded effects (both agonism and antagonism) and acute application^25–26^, features well suited to the study of essential genes and post translational processes. Peptide-based and active-site directed cysteine-reactive inhibitors are widely utilized to inactivate and study caspase function. Exemplary electrophilic peptides with established caspase inhibitor activity include peptide-aldehydes (e.g. DEVD-CHO)^27–28^, peptide-chloromethyl ketones and -fluoromethyl ketones (e.g. YVAD-cfk and z-VAD-fmk)^29–30^, and peptide-acyloxymethyl ketones (AOMK)^31–32^. Recent studies have revealed the utility of covalent fragment electrophiles in targeting individual caspases via covalent modification of catalytic cysteine residues^33–34^. However, the overlapping substrate profiles and the generally limited inhibitor selectivity^35–36^ together demonstrate the need for alternative approaches to achieve selective caspase inhibition.

Recent studies, including our own^33, 37^, have revealed three alternative approaches to peptide-based caspase inhibitors. First, allosteric inhibitors have been reported that target caspase-1, -6, and -7^38^, have revealed three alternative approaches to peptide-based caspase inhibitors. First, allosteric inhibitors have been reported that target caspase-1, -6, and -7^38–41^. Second, our recent work revealed that improved selectivity could be achieved using non-peptidic compounds that function by targeting the catalytic cysteine in the precursor or zymogen form of the enzyme^33, 37^. Third, recent studies have revealed that caspases harbor functional and ligandable non-catalytic cysteine residues that are unique to individual caspases. For example, our recent work revealed that mutation of a near active site cysteine residue (C409) in caspase-8 afforded an almost completely inactive enzyme^42^. For pro-caspase-3, nitrosylation of C163 in pro-caspase-3 blocks apoptosis in human T cells^43^. Covalent modification of C264 in Caspase-6 by enantioenriched vinyl-sulfonamide probes afforded selective blockade of enzyme activity^44^. Whether additional non-catalytic caspase cysteines can be selectively targeted by chemical probes remains an open question.

Among all caspases, caspase-2 remains particularly enigmatic. While typically annotated as an apoptosis initiating (initiator) caspase^45^, caspase-2 shares substrates with executioner caspases and can be cleaved by caspase-3^46–47^. Caspase-2 is most well characterized as playing an important role in regulating cell death in response to DNA-damage^13, 16, 48–51^. Caspase-2 acts as a tumor suppressor affording MDM2 cleavage, which promotes p53 stability^16, 50^. More broadly, caspase-2 has been linked to cell death induced by metabolic imbalance^52^, endoplasmic reticulum (ER) stress^53^, and a mechanism of delayed mitosis-linked cell death^54–55^. Independent of its pro-apoptotic functions, caspase-2 has also been tied to cell cycle progression^13^ and regulation of cell proliferation^56^, including by functioning as a checkpoint for supernumerary centrosomes^55^. Pointing towards possible therapeutic opportunities for caspase-2 selective inhibitors, caspase-2 activity has been implicated in neurodegenerative diseases, most notably in the context of Alzheimer’s disease (AD) and other tauopathies, through cleavage of the tau protein, which impairs memory in a murine AD disease model^57^. Intriguingly, caspase-2 depletion has recently been reported as protective in a murine model of NAFLD/NASH^53, 58^, suggestive of possible further therapeutic utility for caspase-2 inhibitors.

Here, using a high coverage cysteine chemoproteomic platform, we identified a highly reactive and ligandable non-catalytic cysteine (C370) that is unique to caspase-2. We found that covalent modification at C370 blocks caspase-2 activity in a proteoform specific manner, occurring only in the monomeric zymogen or precursor form of caspase-2. By combining gel-based activity-based protein profiling (ABPP) with a *tobacco etch virus* (TEV)^59^ protease activation assay, we assessed the inhibitory activity of a panel of electrophilic compounds, which guided our discovery of active and control compounds spanning multiple electrophilic chemotypes. Competitive cysteine chemoproteomic analysis of compound-treated lysates and cells confirmed caspase-2 target engagement and high caspase-2 selectivity for prioritized leads. A heretofore unreported pan-caspase reactive chemotype is also revealed. Providing evidence to support compensatory caspase activity, cell-based treatment with C370-directed inhibitors failed to afford protection from etoposide-mediated apoptosis. Taken together, our study provides a comprehensive toolbox of chemical probes, proteomic platforms, and functional assays to guide ongoing and future proteoform-selective caspase inhibitor campaigns.

## RESULTS

### Mass spectrometry-based chemoproteomics identifies a highly reactive non-catalytic cysteine residue (C370) in caspase-2

Prior studies, including our own, revealed that mass spectrometry-based chemoproteomic measurements of intrinsic cysteine reactivity towards the pan-cysteine reactive probe iodoacetamide alkyne **(IAA**; **Figure 1A** and **Figure S1**) correlate with residue functionality^42, 60^. Therefore, our first step to identify functional and ligandable cysteine residues in caspase-2 was to deploy a high coverage isotopic tandem orthogonal proteolysis activity-based protein profiling (isoTOP-ABPP)^60–61^ to assay caspase cysteine reactivity. Following the workflow shown in **Figure 1A**, we assayed the IAA-reactivity of cysteine residues in lysates derived from Jurkat cells, a primary acute lymphoblastic leukemia (ALL) cell line, in which many caspases show elevated expression^62^. By pairing the isoTOP-ABPP workflow with single-pot solid-phase-enhanced sample-preparation (SP3) and high field asymmetric ion mobility spectrometry (FAIMS)^63–64^, we generated high coverage reactivity datasets that compared cysteine reactivity with 100 μM versus 10 μM **IAA**—highly reactive cysteines, termed “hyper-reactive” residues were classified based on the calculated isoTOP-ABPP ratio values near to zero [Log_2_(R_heavy:light_) = Log_2_(R_10:1_) = -1.0–1.0], consistent with saturation of labeling at low concentrations of iodoacetamide alkyne.

**Figure 1.**
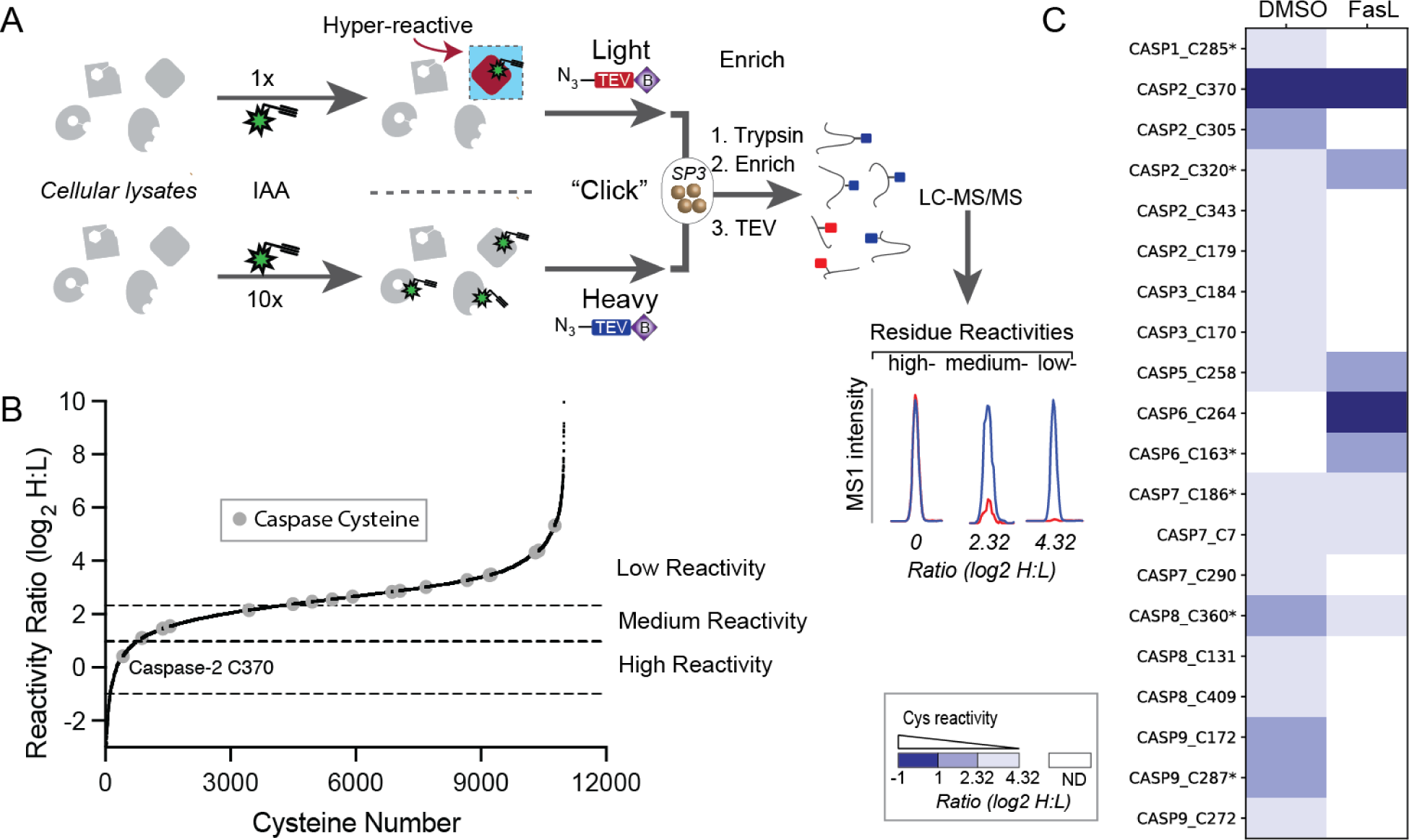
Isotopic tandem orthogonal proteolysis–activity-based protein profiling (isoTOP-ABPP) to stratify the reactivity of caspase cysteine residues. (A) IsoTOP-ABPP workflow used here, which was modified from the previously published methods^60–61^ to incorporate single-pot, solid-phase-enhanced sample-preparation (SP3) cleanup^72^. Lysates are treated with either 10 or 100 μM IAA followed by copper-catalyzed azide-alkyne cycloaddition (CuAAC) conjugation to the previously reported isotopically labeled *tobacco etch virus* (TEV)-cleavable biotinylated peptide tags^60^. The samples are then combined, subjected to SP3 cleanup and on-resin trypsin digest, enrichment on streptavidin resin followed by TEV proteolysis, and liquid chromatography– tandem mass spectrometry (LC-MS/MS) analysis. isoTOP-ABPP ratios (log_2_ H:L) are calculated from the MS1 ion intensity ratios for heavy-versus light-labeled peptides for peptides from non-apoptotic Jurkat cells. Ratio thresholds: high” reactivity: Log_2_(R_heavy:light_) = -1.0 – 1.0, “medium” reactivity: Log_2_(R_heavy:light_) = = 1 - 2.32, “low” reactivity Log_2_(R_heavy:light_) = >2.32. ND-Not detected. (B) Waterfall plot showing isoTOP-ABPP reactivity analysis of cell lysates derived from viable non-apoptotic Jurkat cells. (C) Comparison of the measured isoTOP-ABPP reactivity ratios for caspase cysteines identified in analysis of vehicle treated (DMSO) and apoptotic (50 ng/μL FasL, 4h) cellular lysates. Non-apoptotic experiments (n = 6) and Apoptotic experiments (n = 5). All MS data can be found in Table S2.

In aggregate the **IAA**-reactivity was quantified for 11457 total cysteines from 4153 total proteins (**Table S2**). Eighteen total caspase cysteines from seven caspases were quantified across all replicate experiments. Unexpectedly, given the established nucleophilicity of protease catalytic thiolate nucleophiles^65^, non-catalytic cysteine (C370) in caspase-2 was found to be the most highly reactive residue and only hyper-reactive caspase cysteine across all caspase cysteines quantified (**Figure 1B** and **1C** and **Figure S2A**). As caspase activation is tightly regulated, with protease activity only unleashed during apoptosis, we also extended this analysis to cellular lysates derived from cells undergoing extrinsic [Fas-Fas ligand (FasL)-induced] apoptosis^66–67^. This analysis revealed that C370 remains highly reactive towards IAA in lysates derived from apoptotic cells (**Figure 1C** and **Figure S2B**).

### C370 is highly conserved, active-site proximal, and modestly impacts the activity of recombinant active caspase-2

Both the catalytic cysteine (C320) and active-site proximal C370 in caspase-2 are highly conserved across species (**Figure S3A,B**)^1^. Additionally, C370 is uniquely found in caspase-2 (**Figure S3C**), unlike the catalytic C320, which is shared across all human caspases (**Figure S3D**).

As our prior study had shown that mutation of an IAA-reactive non-catalytic cysteine in caspase-8 nearly completely abolished protein function^42^, we postulated that C370 in caspase-2 may serve a similar regulatory role. To assess whether mutation of C370 adversely affected caspase-2 activity, we recombinantly expressed caspase-2. Comparison of the relative activity of wild-type active caspase-2 with C370A mutated protein revealed a modest ∼10% decrease in enzyme activity, measured under saturating conditions with the Ac-VDVAD-AFC fluorogenic substrate (**Figure S4A**).

Given the seeming contradiction between the elevated intrinsic reactivity of C370 and its seeming modest impact on the active enzyme activity (**Figure S4A**), we speculated that the C370 IAA-reactivity might be favored in the precursor or zymogen form of the enzyme. Consistent with this hypothesis, in the x-ray crystal structure of active caspase-2 (PDB: 1PYO), Cys370 appears buried (**Figure S4B,C**). Homology modeling using I-TASSER^68^, revealed that in a more pro caspase-like conformation^37, 69–71^, C370 is located on the C-terminal loop adjacent to the active site region (**Figure S4C**). The conformational changes caused by the 180° flip of the C-terminal loop and subsequent orientational shift of the sulfhydryl putatively affording increased solvent accessibility for C370.

### Zymogen- and caspase-2 specific labeling of C370 by electrophilic alkyne-probes

The homology models together with previous findings that both catalytic and non-catalytic caspase cysteines are amenable to covalent modification in the protease’s zymogen form^33, 44, 73^ prompted us to establish a gel-based activity-based assay to further probe the labeling specificity of C370 in caspase-2. Recombinant pro-caspase-2 (proCASP2) was expressed harboring D333A and D347A mutations, which prevent caspase cleavage and activation, together with single and dual C370A and C320A mutations to assess cysteine specificity (See **Table S1** for a summary of all constructs). We subjected each protein to gel-based ABPP comparing the labeling of each protein with **IAA** or caspase-directed **Rho-DEVD-AOMK** [Asp-Glu-Val-Asp-(2,6-dimethyl benzoyloxy)-methyl ketone] probe (**Figure S1**)^74^, to assay general cysteine reactivity and caspase activity, respectively (**Figure 2A,B**). We observed that the widely cysteine-reactive **IAA** labels both C370 and C320. In contrast with the relatively robust zymogen labeling, the active form of caspase-2 only shows trace labeling by **IAA (Figure 2B**). Consistent with prior studies, we observed that the **Rho-DEVD-AOMK** labeled the catalytic nucleophile C320, with mutation of this residue completely blocking labeling. We were initially surprised to observe labeling of the zymogen form of the enzyme by **Rho-DEVD-AOMK**, as this probe is tailored to react with catalytically competent caspases. As prior studies have shown that the zymogen form of caspase-2 exists as a mixture of inactive monomer and catalytically competent dimeric forms^45^, this observed labeling may be rationalized in part by a mixed population of active and inactive enzyme.

**Figure 2.**
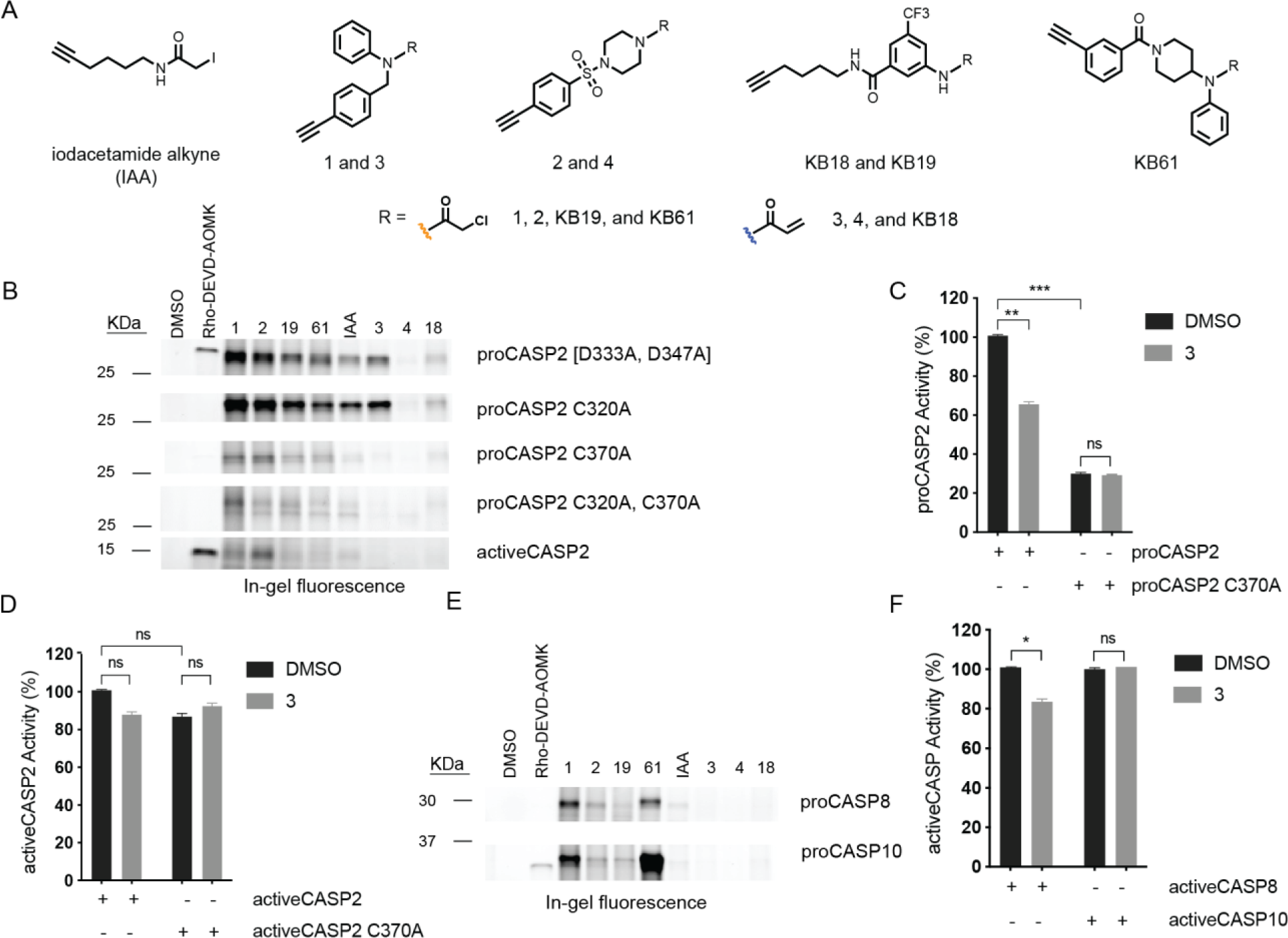
Alkyne probe 3 preferentially labels C370 over other caspase cysteines to afford partial blockade in pro-caspase activity. (A) Structures of cysteine reactive clickable probes including previously reported **IAA**^60^ and **KB18**, **KB19**, and **KB61**^33^, as well as compounds **1**–**4,** unique to this study. (B) Gel-based ABPP analysis of recombinant caspase-2 constructs harboring the indicated mutations in Jurkat whole cell lysates treated for 1h with the indicated compounds (10 µM for all click probes except for IAA, which was analyzed at 1 µM, and **Rho-DEVD-AOMK** at 2 µM). (C) Assessment of compound-induced changes in protease activity for proCASP2 and proCASP3_C370A proteins subjected to **3** (100 µM, 2h, 30 °C) followed by multimode plate reader fluorescence analysis of caspase activity using the fluorogenic substrate Ac-VDVAD-AFC (1 µM) under kosmotropic conditions (sodium citrate)^76–77^. (D) Relative activities of activeCASP2 in the absence (Black) or presence of **3** (Gray) (100 µM) assessed using fluorogenic substrate Ac-VDVAD-AFC. (E) Gel-based ABPP analysis of recombinant pro-caspase-8 (proCASP8) and pro-caspase-10 (proCASP10) subjected to the indicated compounds (10 µM for all click probes except for IAA, which was analyzed at 1 µM, and **Rho-DEVD-AOMK** at 2 µM). (F) Relative protease activities of activeCASP8 and activeCASP10 in the absence (Black) or presence of **3** (Gray) (100 µM, 2h, 30 °C) assessed using fluorogenic substrate Ac-VDVAD-AFC. For Figure 2D, **2E** and **2F**, experiments were performed in triplicate. Full length gels shown in **Figure S8** and **S9**. For C, D, F Data represent mean values and standard deviation. Statistical significance was calculated with unpaired Student’s t-tests, ns, not significant, * p<0.05, ** p<0.01, ** *p<0.001, NS p>0.05.

Given the established value of cysteine-reactive probes as tool compounds, and even as lead candidates for potential therapies, we next set out to determine whether more elaborated compounds featuring more attenuated electrophiles could label C370 selectively. To this end, we obtained a small panel of structurally diverse aliphatic and aromatic cysteine-reactive alkyne-containing scout fragments, including newly synthesized analogues (**1-4)** and previously reported compounds (**KB18**, **KB19**, **KB61**)^33^ (**Figure 2A** and **Figure S1**).

We labeled our panel of recombinant caspase-2 proteins with these additional cysteine-reactive probes (**Figure 2B**). In contrast with active caspase-2, which labeled poorly with nearly all probes tested, we found that the D333A, D347A mutant form of the protein (zymogen-like) labeled robustly with a number of probes. Mutation of C370A nearly completely blocked labeling, which supports our hypothesis that the primary reactive cysteine in the zymogen protein is the non-catalytic C370 residue, which was further supported by the robust probe labeling of the C320A mutated protein. C370 was labeled robustly by most chloroacetamide probes, including **1** and **2**. As chloroacetamides generally show heighted proteome-wide reactivity, we also tested several acrylamide probes (**3**, **4**, **KB18**) and gratifyingly found that acrylamide **3** labeled caspase-2 at a level comparable to many of the chloroacetamide compounds.

Given the aforementioned catalytic activity of pro-caspase-2 (proCASP2), we next sought to assess whether modifications to C370 in the zymogen form of caspase-2 by **3** would impact activity. Pre-treatment of proCASP2 with **3** partially blocked enzyme activity in a cysteine-dependent manner (**Figure 2C**). While zymogen forms of caspases generally show low activity^75^, the addition of kosmotropes (e.g. sodium citrate) can increase activity^76–77^. Consistent with these prior studies, we found that addition of both Dithiothreitol (DTT) and sodium citrate afforded increased activity of the D330A, D347A mutant enzyme (**Figure S5**), as assayed using the fluorogenic substrate Ac-VDVAD-AFC^78^. We also observed that C370A mutation afforded a substantial increase in zymogen activity (**Figure 2C**), in contrast to our analysis of activeCASP2 C370A (**Figure 2D**). Treatment of activeCASP2 with **3** afforded no significant inhibition, consistent with a model whereby **3** labels the zymogen form of caspase-2 (**Figure 2D**).

The partial inhibition of caspase-2 by **3** can be rationalized by either incomplete labeling of the protein or by an allosteric inhibitory mechanism, where C370 modification affords only partial inactivation. Caspase-2 labeling by **3** saturated at approximately 25 µM, which would align with complete labeling (**Figure S6**). However, bottom-up LC-MS/MS analysis of recombinant protein labeled by **3** (100 µM) revealed only partial modification under conditions of excess probe (**Figure S7**). These data point to only partial modification of caspase-2 as the likely explanation for the incomplete inhibition.

Based on our prior cysteine chemoproteomic datasets^33, 79–80^, caspase cysteines generally react poorly with acrylamide-substituted compounds. Therefore, we postulated that the acrylamide in **3** might offer the additional benefit of improved caspase selectivity. As expected, no substantial labeling was observed for **3**-treated proCASP8 and proCASP10, as analyzed by gel-based ABPP (**Figure 2E**). Furthermore, only modest inhibition of activeCASP8 was observed after treatment with **3** (**Figure 2F**).

### Assessment of the scope of chemotypes that label Caspase-2 at C370.

Given the challenges associated with lead optimization for electrophilic compounds, including idiosyncratic structure–activity relationship (SAR) that can be complicated by difference in pre- and post-alkylation binding poses, and the absence of high-resolution co-crystal structures, we opted to pursue multiple parallel lines of SAR analysis, including synthesis of analogues of **3 (Figure 3A**) and broadening the chemotypes evaluated by obtaining a small panel of putative electrophilic scaffolds (**Figure S1**, **P01**, **P02**, **P03, P04, P04, P05,** and **P06**). Gel-based ABPP analysis revealed that methylphenyl propiolate **P01**, beta-nitrostyrene **P03**, and nitrovinyl benzodioxole **P06** all label C370 (**Figure S10**). As the potency of these compounds compares favorably to **3**, with near complete competition of C370 at 10 µM dose (**Figure S10**), we additionally chose to synthesize and test a focused set of methyl phenylpropiolate **P01** analogues (**Figure 3A** and **Figure S1**).

**Figure 3.**
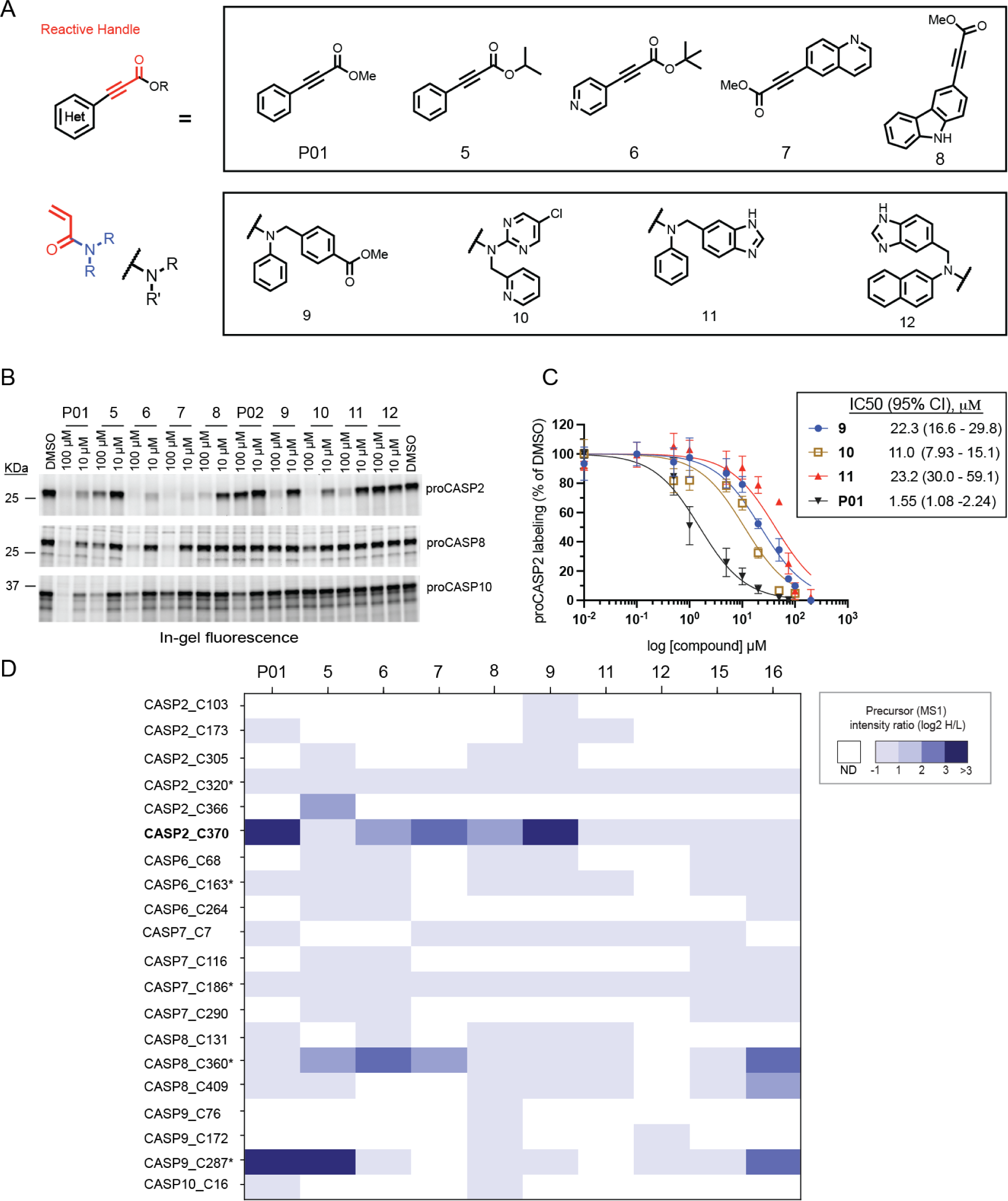
Fragment electrophiles selectively label proCASP2 at C370. (A) Structures of electrophilic compound library of **3** and **P01** analogues. (B) Gel-based ABPP analysis of the indicated recombinant proteins in whole cell lysates treated for 1h with the indicated compounds followed by click conjugation to compound **3** (10 µM) for proCASP2 and **KB61** 10 µM) for proCASP8 and proCASP10. Full length gel images in supporting information **Figure S12** (C) Apparent IC50 curve for blockade of **3** labeling of proCASP2 by **the indicated compounds**. CI, 95% confidence intervals. (D) Caspase cysteines quantified in competitive isoTOP-ABPP analysis of Jurkat cell lysates treated with **P01**, **5** and **6** (25 µM) and **7**, **8**, **9**, **11**, **12**, **14**, **15**, and **16** (100 µM). Competition ratios are calculated from the mean quantified precursor intensity (log_2_H/L) for DMSO (H, heavy) versus compound-treatment (L, light). * indicates catalytic cysteine residue. ND-Not detected. For C, data represent mean values ± STDEV for at least three independent experiments. For D, experiments were conducted in at least two biological replicates. All MS data can be found in **Table S3**.

Using a gel-based ABPP screening format, we compared the potency of library members **5-16** (**Figure 3B**). We find that the N-phenyl-N-benzyl analogue modifications to the benzyl moiety (e.g. methyl ester **9** and benzimidazole **11**) show similar (∼20 µM) apparent IC50 values (**Figure 3C**). Replacement of the aniline portion with a more electron withdrawing chloropyrimidine group **10** (**Figure 3C**) afforded an increase in potency (apparent IC50 11.0 µM). Naphthyl compound **12** showed no appreciable competition of probe labeling for caspase-2, indicating that large modifications to the aniline ring are not tolerated. For the **P01** analogues, we find that benzyl, phenyl, and isopropyl ester modifications decrease compound potency, consistent with a potential steric clash with the bulkier substituents (**Figure S11)**. Curiously, the bulky tert-butyl pyridine analogue **6** did not suffer from decreased potency, hinting that the pyridine species may alter the electrophilicity and/or binding pose relative to the non-heterocyclic analogues. **P01** analogues **7** and **8**, which feature larger aromatic substituents (quinoline and carbazole, respectively) both maintained caspase-2 labeling, with the carbazole analogue showing modestly decreased potency.

We then extended our SAR analysis to assess labeling of recombinant proCASP8, proCASP10 and activeCASP3 (**Figure 3B, Figure S12, Figure S13 and Figure S14)**. Gratifyingly, none of the compound **3** analogues (**9-12**) showed any appreciable caspase-8/10 labeling. In contrast, we were surprised to observe, given our aforementioned observations that caspase catalytic cysteines generally react poorly with Michael acceptors, that **P01** together with analogues **5** and **6** afford marked labeling of both caspase-8 and -10 (**Figure 3B** and **Figure S12**, **Figure S13 and Figure S14**). The bulkier analogues **7** and **8** showed, respectively, partial and near complete selectivity for C370 over the catalytic cysteines in caspase-8 and caspase-10 (**Figure 3B and Figure S12**). These same compounds did not block labeling of recombinant caspase-3 as visualized using the **Rho-DEVD-AOMK** probe (**Figure S15**) and had minimal effects on recombinant activeCASP2 and activeCASP2_C370A activity (**Figure S16**).

To assess whether these selectivity profiles extended to endogenous caspases, we next subjected our compound library to competitive isoTOP-ABPP analysis in Jurkat cell lysates, following the workflow shown in **Figure S17**, as has been reported previously^33^. While we initially struggled to consistently detect C370, we found that the use of a FAIMS device during acquisition (**Figure S18**) afforded a marked increase in coverage of a number of quantified caspase cysteines, including C370. This observation aligns with prior reports of online gas phase fractionation improving coverage for both cysteine chemoproteomics^81^ as well as other low abundance ions^63, 82^. In aggregate across 10 compounds screened, we quantified 11238 cysteines from 4003 proteins (**Table S3**). 20 total cysteines from six caspases (**Figure 3D**). Notably we only detect C264 in a handful of datasets, which is consistent with previous reports indicating difficulties with detection of this peptide from endogenous protein^44^.

Gratifyingly and consistent with our gel-based ABPP analysis, we observed that compounds **P01, 6-9, and 11)** all afford high precursor intensity ratios for peptides harboring C370, consistent with compound modification at C370. Inactive control compound **12** showed no appreciable labeling for any caspase cysteine detected. Again, consistent with our gel-based analysis we observe that a number of the **P01 analogues** are reactive towards initiator caspase-8 and -9 catalytic cysteines. This finding further supports the likelihood of this chemotype as broadly caspase reactive.

### Establishing TEV-protease activatable caspase-2

Our functional studies using recombinant proCASP2 **(Figure 2C**) hinted that C370 labeling afforded inhibition of the pro-enzyme. However, these assays were complicated by the relatively low overall activity of the pro-enzyme, particularly in the absence of a kosmotropic agent (i.e. citrate). Therefore, our next step was to establish an assay that could test whether labeling of C370 in the inactive form of the enzyme would inhibit the active enzyme. We were motivated by previous reports that utilized engineered *tobacco etch virus* (TEV) protease-activatable caspase constructs^83–85^ for cell-based studies. To test whether this technology would extend to *in vitro* caspase-2 inhibition studies, we generated proCASP2TEV and proCASP2TEV_C370A by replacing both caspase-2 cleavage sites (D333 and D347) with TEV (ENLYFQG) cleavage motifs (**Figure S19)**. Gratifyingly, upon addition of TEV protease, we observed marked TEV-dependent increased catalytic activity (**Figure 4A**) and near complete conversion to active caspase-sized species as detected by gel-based ABPP (**Figure 4B-E**). The activation was rapid, occurring within 5-10 minutes, and only required modest (2.5 µM) concentrations of TEV protease to achieve complete activation. Comparison of the kinetic parameters of proCASP2TEV and activeCASP2 revealed that the introduction of the TEV cleavage motifs afforded no substantial change in k_cat_ or K_m_ (**Figure 4F**). Consistent with our proCASP2 constructs, we found that proCASP2TEV was labeled efficiently by both **Rho-DEVD-AOMK** and **3**, with the latter showing C370-specific labeling, as assayed by gel-based ABPP (**Figure 4B, 4D-E** and **Figure S20**). DEVD-Rho labeling was only detectable for the 18 kDa cleavage product, which contains C320, whereas **3**-labeling was only observed for the 10 kDa cleavage product, which harbors C370 (arrows in **Figure 4D and 4E**). This striking probe-dependent labeling pattern provides additional evidence supporting the specificity of **3** for C370.

**Figure 4.**
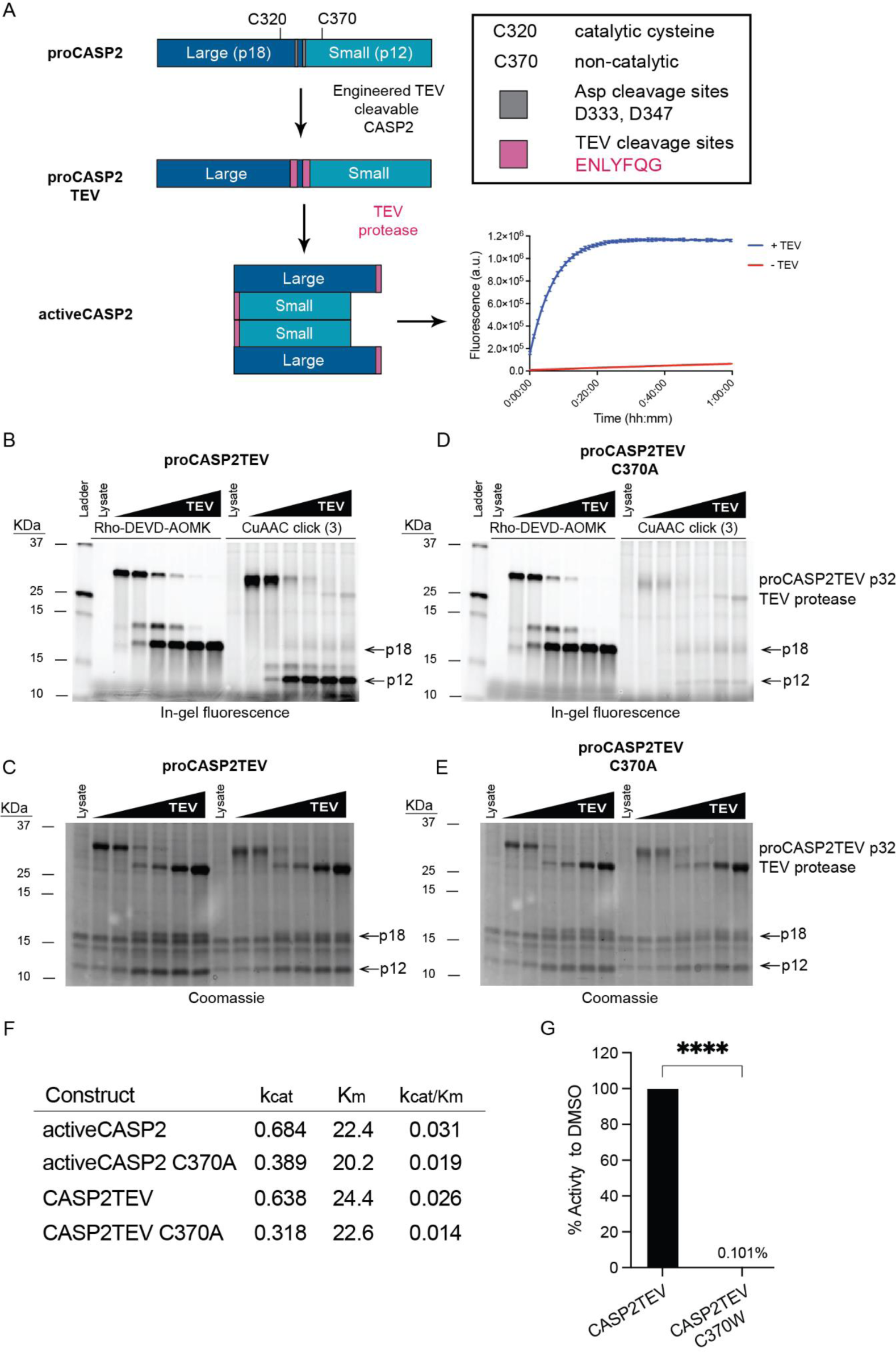
TEV-cleavable proCASP2 is activated by TEV protease with minimal effects on enzyme kinetics. (A) Replacement of caspase-2 cleavage sites (D333 and D347) with TEV (ENLYFQG) cleavage motifs affords engineered proCASP2TEV and proCASP2TEV_C370A proteins, which exhibit TEV-dependent increase in caspase activity towards Ac-VDVAD-AFC fluorogenic substrate. (B-E) Gel-based ABPP analysis of (B,C) proCASP2TEV and (D,E) proCASP2TEV_C370A, which were treated with either **Rho-DEVD-AOMK** (1 µM final concentration) or compound **3** (10µM final concentration) for 1h, followed by activation with TEV protease at increasing concentrations (0 < 0.1 < 0.5 < 1.0 < 2.5 < 5.0 µM), and click conjugation to Rho-Azide for **3**-treated samples. In-gel fluorescence analysis (B,D) and coomassie staining (C,E) were used to visualize compound labeling and protein abundance, respectively. Arrows indicate the ∼18 kDa **Rho-DEVD-AOMK**-labeled band and a ∼12 kDa **3**-labeled band. (F) Michaelis-Menten kinetics comparing proCASP2TEV activity to activeCASP2 activity. (G) Activity assay of proCASP2TEV with proCASP2TEV C370W mutant. For G, samples were prepared in three technical replicates. Statistical significance was calculated with unpaired Student’s t-tests, **** p<0.005. NS p>0.05. Arrows indicate cleaved or processed caspase.

### Autoactivation complicates TEV-dependent inhibition assays

Motivated by the TEV-dependent high activity of our engineered construct, we next asked whether treatment with our lead caspase-2 electrophilic compounds would block TEV dependent activity. We subjected our lead construct proCASP2TEV to treatment with excess of compounds **P01**, **8**, **9**, and **10** (100 µM), followed by activation by TEV protease in the presence of excess DTT (5 mM) to quench unreacted compound—as TEV protease is a cysteine protease, addition of DTT is required to prevent TEV catalytic cysteine alkylation, which could result in spurious measures of inhibition. Surprisingly, we observed minimal compound-dependent inhibition, both in the presence and absence of TEV protease (**Figure S21**). In contrast, gel-based ABPP analysis revealed comparable labeling of proCASP2TEV to that observed for our proCASP2 constructs (**Figure S22, S23**).

Somewhat perplexed by the seemingly inconsistency between the inhibition data and the gel-based ABPP analysis, we speculated that the recombinant protein might harbor a mixture of proteoforms, including both compound-sensitive and insensitive species. Our previous observation of the partial labeling of the C370 peptide by **3** (**Figure S7**) in proCASP2 was consistent with this hypothesis. Also consistent with this hypothesis, we observed that proCASP2TEV and proCASP2TEV_C370A exhibit autocleaving properties as indicated by the presence of an 18 kDa **Rho-DEVD-AOMK-**reactive form of caspase-2 detected in the immobilized metal affinity chromatography (IMAC)-purified protein (**Figure S24)** and observed TEV-independent activity (**Figure S25**). In contrast, autocleavage of proCASP2 and proCASP2_C370A was not detected (**Figure S26**). In light of these confounding factors, we opted to generate the C370W mutated protein to test whether genetic introduction of a bulky residue, which mimics small molecule labeling, would afford a complete blockade of enzyme activity. The proCASP2TEV_C370W protein, which was obtained in high yield and purity (**Figures S27**), showed no detectable activity (**Figure 4G**).

### Preferential labeling of the monomeric form of pro-caspase-2

Most initiator caspases (e.g. caspase-8, -9 and -10) exist primarily in an inactive monomeric state, with dimerization resulting in rapid activation^86^. While proCASP2 is thought to function as an initiator caspase, unique among caspases, its physiological pro-enzyme form (with CARD pro-domain) exists as a mixture of monomeric and dimeric states^45^. Therefore, we hypothesized that our recombinant pro-caspase, which lacks the CARD domain, might also exist as a mixture of oligomeric states. We subjected our proCASP2TEV and proCASP2TEV_C370A proteins to size exclusion chromatography (SEC), which revealed an elution profile for recombinant proCASP2TEV consistent with a mixed population of putative monomer, dimer, and a cleavage product (∼ 30.0 kDa; **Figure 5A, 5B**). SEC analysis of proCASP2TEV_C370A revealed two major species matching the molecular weights of monomeric and dimeric caspase-2 (**Figure 5C, 5D**). Notably, both proCASP2TEV and proCASP2TEV_C370A samples also harbored trace amounts of putative active caspase-2 species (∼18 kDa bands), which were not visualized by Coomassie staining but were detectable by the Rho-DEVD-AOMK activity-based probe (**Figure 5B, 5D**). Similarly, mixed putative monomer and dimer populations were observed for gel filtration analysis of the proCASP2 and proCASP2_C370A constructs (**Figure S28** and **Figure S29**), with the exception of the presence of previously noted ∼25 kDa intermediate species that was only observed for proCASP2_TEV.

**Figure 5.**
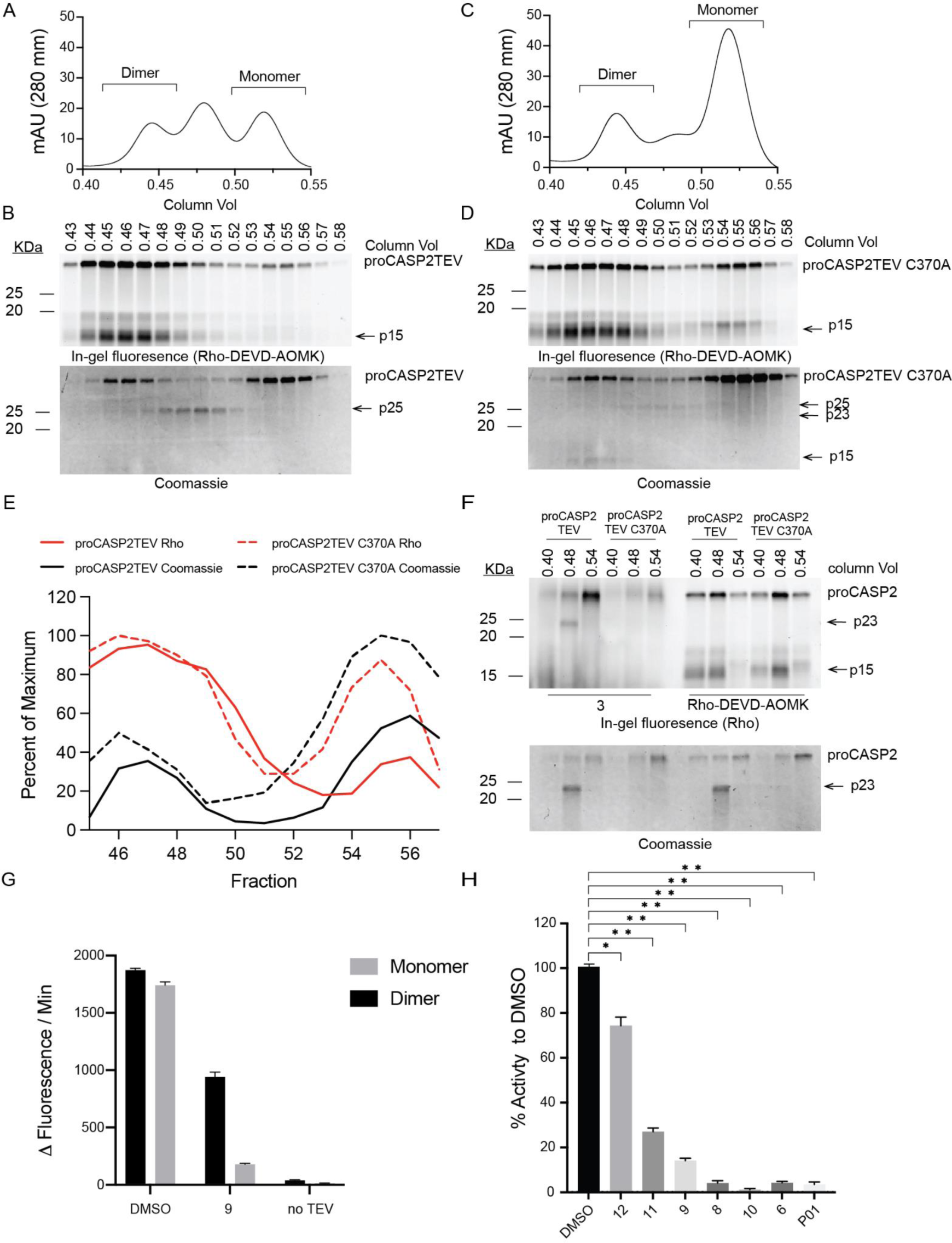
State-dependent inhibition of caspase-2. (A-D) Gel filtration analysis of (A,B) proCASP2TEV (C,D) proCASP2TEV_C370A showing (A,C) the UV absorbance at 280 nm (A280) for each sample and (B,D) gel-based ABPP analysis of the indicated fractions labeled with **Rho-DEVD-AOMK** (1 µM). (E) Percent of maximum dimeric and monomeric fractions of proCASP2TEV and proCASP2TEV C370A. Relative band intensities were determined for coomassie bands (black) and **Rho-DEVD-AOMK**-labeled bands (red). Percent components of total dimer and monomer forms calculated from the area under the curve (AUC) of dimeric column volume fractions (0.40 - 0.46) and monomeric column volume fractions (0.50 - 0.55) for proCASP2TEV activity assessed by labeling with **Rho-DEVD-AOMK** and total protein assessed with coomassie staining. (F) Gel-based ABPP analysis of the indicated gel filtration fractions for of proCASP2TEV and proCASP2TEV C370 with either **Rho-DEVD-AOMK** or **3,** with the latter samples conjugated to Rho-azide by click chemistry. (G) Relative activity of monomeric versus dimeric fractions of proCASP2TEV subjected to the indicated treatment by either compound **9** (100 µM, 1h, 30 ℃) or vehicle (DMSO) followed by addition of TEV protease or vehicle and activity analysis under kosmotropic conditions using **Ac-VDVAD-AFC** fluorogenic substrate (H) Measured percent activity for monomeric fractions of proCASP2TEV treated with the indicated compounds (100 µM, 1h, 30 ℃) followed by addition of TEV protease and kosmotropic buffer. Arrows indicate cleaved or processed caspase. For G and H data represent mean values and standard deviation, n = 3 and statistical significance was calculated with unpaired Student’s t-tests, * p<0.05, ** p<0.01.

Somewhat unexpectedly, we observe striking elution-dependent difference in the relative intensities of caspase-2 labeling by Rho-DEVD-AOMK. While proCASP2_TEV was observed to be a roughly 1:1 mixture of caspase-2 monomer and dimer, we observed ∼three-fold more labeling relative to protein abundance for the dimer-containing fractions (0.30 - 0.45) compared to the monomer-harboring fractions 0.48 - 0.55) (**Figure 5E**). These findings support that monomeric caspase-2 is generally less active towards the Rho-DEVD-AOMK activity-based probe.

Guided by these observations, we next asked whether these isolatable forms of caspase-2 would behave differently both in terms of measured catalytic activity and inhibition by our C370-reactive compounds. Comparative TEV-dependent activity was observed for both caspase-2 monomer/dimer species (**Figure 5G**). Consistent with our previous observation of partial inhibition of proCASP2 by **3 (Figure 2C),** the dimeric fractions only exhibited partial inhibition by **compound 9**. In contrast, the monomeric fractions showed near complete inhibition by **9** (**Figure 5G**). Extension of this analysis to our panel of electrophilic compounds, revealed, for the monomeric fractions, near complete inhibition of proCASP2TEV for all compounds that showed caspase-2 labeling as detected by gel-based ABPP (**Figure 5H**).

Given that we had observed a putative cysteine-mutant-dependent increase in the fraction of monomeric protein detected by gel filtration for the proCASP2TEV_C370A protein (**Figure 5C**), we additionally wondered whether compound labeling at C370 might similarly afford an increase in monomeric caspase-2. Consistent with this model, filtration analysis after compound labeling revealed a shift in the protein elution profile towards increased monomeric species (**Figure S30**). Alongside our inhibition and gel-based analysis data, this finding supports that compound labeling at C370 favors the monomeric zymogen form of caspase-2.

### In cell engagement of C370 is achieved by both broadly caspase reactive and caspase-2 selective compounds

As we postulated that the caspase-2 monomer-dimer equilibrium is likely sensitive to protein concentration, we next wondered whether caspase-2 sensitivity to compound labeling would be maintained in cell-based treatments. Using our established isoTOP-ABPP workflow (**Figure S17**), we subjected Jurkat cells to labeling with both active and inactive compounds followed by lysis, IAA labeling, enrichment, sequence specific proteolysis, and LC-MS/MS analysis. In aggregate, we identified 9089 cysteines from 3506 proteins (**Table S5**). As with our lysate-based labeling, we achieved high caspase coverage, quantifying 21 total cysteines from eight caspases (**Figure 6A**). The SAR observed across the library screen was generally comparable to that observed for our lysate-based labeling studies. Compound **P01** did, however, show modestly increased ratios for our cell-based analysis across all detected caspase cysteines.

**Figure 6.**
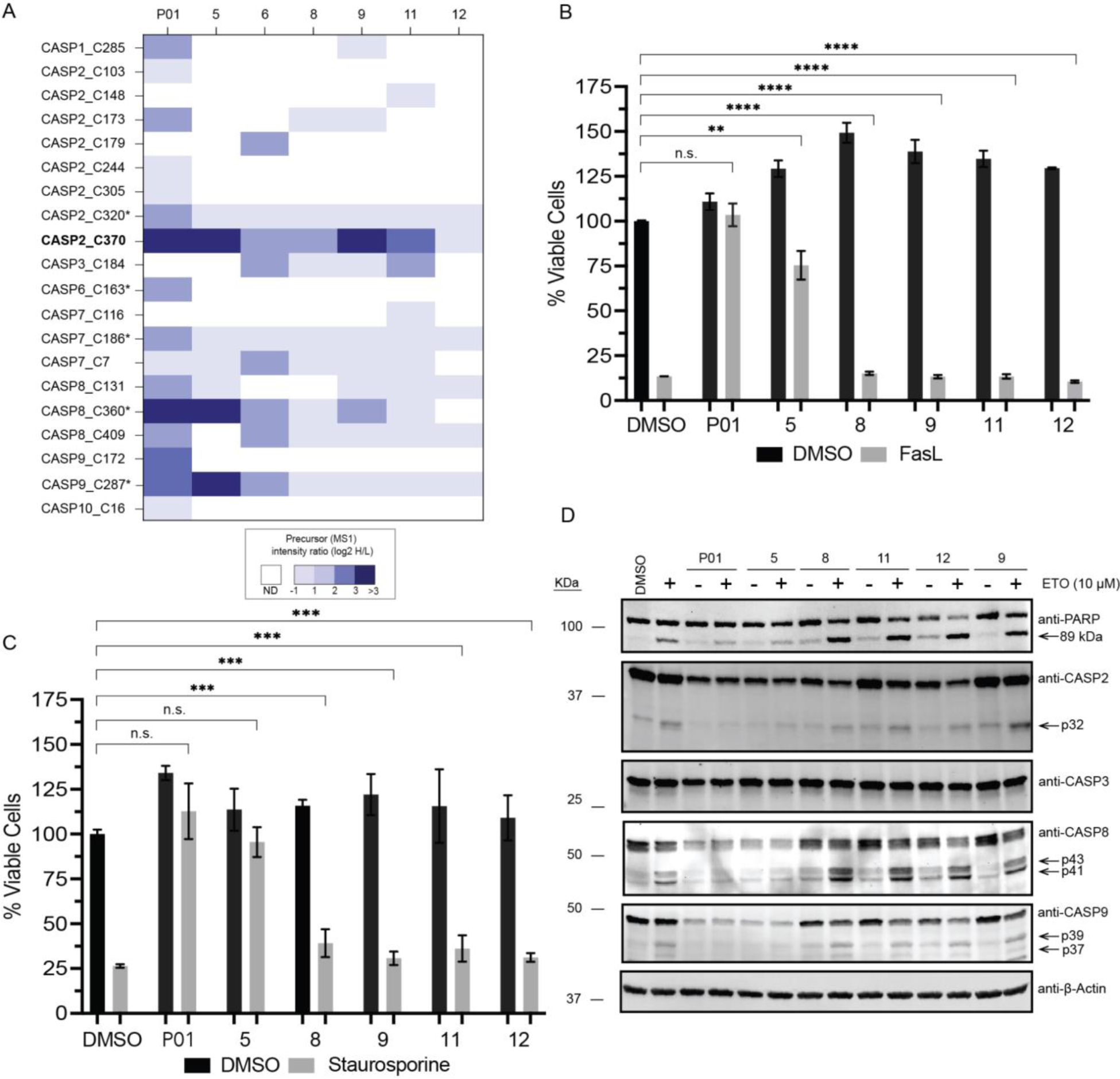
In situ labeling of caspase-2 by electrophilic lead compounds and blockade of extrinsic and intrinsic apoptosis by promiscuous caspase inhibitors. (A) Caspase cysteines quantified in competitive isoTOP-ABPP analysis of Jurkat cells treated with **P01**, **5**, **6** (25 µM) and **8**, **9**, **10**, **11**, and **12** (100 µM). Competition ratios are calculated from the mean quantified precursor intensity (log_2_ H/L) for DMSO (H, heavy) versus compound-treatment (L, light). * indicates catalytic cysteine residue. Experiments were conducted in at least two biological replicates. ND-Not detected. (B,C) Comparison of effects of the indicated compounds [1h pre-treatment with **P01**, **5**, and **6** (25 µM) and **8**, **9**, **10**, **11**, and **12** (100 µM)]. on (B) FasL (50 ng/mL, 4h)-induced apoptosis or (C) staurosporine (STS, 1µM, 4h)-induced apoptosis of Jurkat cells assayed by CellTiter-Glo^®^. (D) Comparison of the effects of the indicated compounds [1h pre-treatment with **P01**, **5**, and **6** (25 µM) and **8**, **9**, **10**, **11**, and **12** (100 µM)] on etoposide (10 µM, 10h)-induced apoptosis with cleavage of PARP, CASP2, CASP8, and CASP9 visualized by western blotting. Full blots shown in **Figure S31** and **Figure S32.** Viability experiments were performed in biological triplicate. All MS data can be found in **Table S5**. For B and C data represent mean values and standard deviation, n=3. Statistical significance was calculated with unpaired Student’s t-tests, * p<0.05, ** p<0.01, *** p<0.005. ****p<0.001. NS p>0.05. Arrows indicate PARP and caspase-cleaved bands.

Guided by our prior work using small molecule inhibitors to delineate the relative involvement of caspase-8 and -10 in initiating extrinsic apoptosis^33^ and the widespread use of the pan-caspase inhibitor DEVD-fmk for pinpointing caspase-dependent processes, we assessed compound-dependent changes to apoptosis. We observe that the broadly caspase reactive **P01** afforded a near completed blockade of both intrinsic and extrinsic apoptosis, induced with FasL and staurosporine (STS), respectively (**Figure 6B,C**). Compound **9,** which was observed to potently label catalytic cysteines in caspase-8 and -9 initiator caspases as well as C370 in caspase-2, also afforded a near complete blockade of both forms of cell death. While caspase-2 selective inhibitors, including compound **9** and **11**, robustly labeled C370 in situ as measured by isoTOP-ABPP, these compounds did not impact FasL-nor STS-induced apoptosis. These findings align with the established functions of caspase-8/9/3 as the primary drivers of these forms of apoptosis.

As previous reports have implicated caspase-2 as a key mediator of cellular response to DNA damage ^13, 16, 48–51^, we next investigated whether caspase-2 inhibitor treatment would block caspase processing induced by etoposide, a widely utilized inhibitor of DNA synthesis which causes double strand breaks. We found that only our non-selective caspase inhibitors (**P01** and **5)** afforded a detectable decrease in etoposide-induced processing of PARP, caspase-2, and caspase-8. These findings indicate that pro-caspase-2 inhibition is insufficient to fully block DNA-damage-induced apoptosis (**Figure 6D**). Intriguingly, treatment with these compounds also leads to a modest decrease in the immunoblot signal for caspase-2, caspase-8, and caspase-9, suggestive of compound-induced changes in protein abundance.

## Discussion

Despite being the most evolutionarily conserved caspase, caspase-2 remains enigmatic, implicated in numerous cellular processes, spanning DNA damage response, tumor suppression, response to oxidative stress, and has even been linked to nonalcoholic steatohepatitis (NASH)^53^, alzheimer’s^57, 87^, and Huntington’s diseases^88^. Thus, there is an increasing interest in the development of caspase-2 selective chemical probes with use for both functional biology and early-stage drug development, demonstrated by recent reports of selective peptide-based inhibitors^89–90^. Here, using a high coverage, FAIMS-enabled chemoproteomics approach, we identify a ligandable, potentially druggable, non-catalytic and active site proximal cysteine (C370). By combining gel-based ABPP and a novel TEV-activation assay, we identify both caspase-2 selective and broadly caspase-reactive lead compounds. We find that covalent modification of C370 occurs in recombinant protein, cells, and cell lysates in a proteoform-selective manner, as demonstrated by the observed zymogen-selectivity and preferential labeling of the monomeric form of caspase-2. To our knowledge, such proteoform-selective caspase-2 inhibitors are unprecedented.

While chemoproteomic analysis confirmed robust labeling of C370 for both cell- and lysate-based analysis, only the more promiscuous caspase lead compounds afforded marked protection from extrinsic and intrinsic apoptosis, including DNA-damage mediated processes. As genetic approaches have linked this latter pathway to caspase-2^13, 16, 50–51^, we attribute the disconnect between chemical inhibition and genetic depletion to compensatory activities of other caspases or non-catalytic functions of caspase-2, for example activities of its CARD domain^91^. As caspase-2 forms comparatively stable homodimers^45^, we cannot rule out the alternative possibility that a small fraction of dimerized caspase-2, which is resistant to covalent modification of C370, could be responsible for the observed sensitivity to etoposide-mediated apoptosis. Delineating these possibilities would benefit from the development of caspase-2 directed degrader molecules or through the use of the DTAG system^92^. Notably, we did observe depletion of caspase-2 after compound treatment, suggestive of compound-induced protein degradation, which could provide inroads into such degradation modalities.

Our findings contribute to the growing body of evidence regarding the presence of ligandable non-catalytic cysteine residues in other caspases, including the recently targeted cysteine residue C264 in caspase-6^44^. As demonstrated by the blockbuster success of targeting such conserved catalytic cysteines in kinases^93–95^, non-catalytic cysteines represent an exciting, yet still largely untapped opportunity for high selectivity caspase inhibitor development campaigns. However, despite our best efforts, we were unable to progress our best lead compounds beyond modest low micromolar potency. Despite this limitation, our use of a structurally matched inactive control compound **12**, allowed us to demonstrate that C370 does exhibit SAR towards electrophilic lead compounds. These findings provide a strong foundation for future efforts, likely enabled by caspase-2 co-crystal structures, to further increase the potency of caspase-2 directed leads. As C370 is uniquely found in caspase-2, these molecules are expected to retain high caspase-2 selectivity.

Identification of proteoform selective molecules is an area of rapidly emerging interest^33, 96–100^. As shown by type II inhibitors that target inactive forms of kinases^101–103^, we expect that targeting pro-caspases should emerge as a general strategy to selectively inactivate individual caspases in cells. An additional feature of pro-caspase inhibitors is that, when compared with active caspase inhibitors, they should fully block downstream caspase activation—active caspase inhibitors are ill equipped to block the rapid caspase activation (<30 min)^104^— and subsequent cleavage of effector caspases that is observed during apoptosis. Therefore, we expect that our study will provide a roadmap for the production of such molecules targeting caspases and related proteases, including through the implementation of activation assays, such as the TEV assay reported here. Looking more broadly towards the burgeoning area of covalent inhibitor development, our finding that the methylphenylpropiolate scaffold **P01** exhibits widespread cysteine reactivity, both towards caspases and more broadly, points towards untapped opportunities for its use as a scout electrophile. In contrast to similar chloroacetamide scout electrophiles, which have proven challenging to progress to more potent chemical probes, we expect this chemotype should progress more readily towards more attenuated acrylamide leads, both for caspases, and across the cysteinome.

## Acknowledgements

This study was supported by National Institutes of Health DP2 OD030950-01 (K.M.B.), Beckman Young Investigator Award (K.M.B.), UCLA DOE Institute (DE-FC02-02ER63421), NSF GRFP (1000235263), TRDRP T31DT1800 (T.Y.), NIGMS System and Integrative Biology 5T32GM008185-33 (L.M.B.), and UCLA’s Cellular and Molecular Biology (CMB) Training Program T32GM007185 (E.A.). We thank all members of the Backus lab for their helpful suggestions and Dennis Wolan for providing plasmids and for the Rho-DEVD-AOMK probe.

## DATA AVAILABILITY

The MS data have been deposited to the ProteomeXchange Consortium (http://proteomecentral.proteomexchange.org) via the PRIDE^105^ partner repository with the dataset identifiers PXD042362 and PXD046269.

## References

1. Kumar, S.; Kinoshita, M.; Noda, M.; Copeland, N. G.; Jenkins, N. A., Induction of apoptosis by the mouse Nedd2 gene, which encodes a protein similar to the product of the Caenorhabditis elegans cell death gene ced-3 and the mammalian IL-1 beta-converting enzyme. Genes & Development 1994, 8 (14), 1613–1626.

2. Yuan, J.; Shaham, S.; Ledoux, S.; Ellis, H. M.; Horvitz, H. R., The C. elegans cell death gene ced-3 encodes a protein similar to mammalian interleukin-1 beta-converting enzyme. Cell 1993, 75 (4), 641–52.

3. Wang, L.; Miura, M.; Bergeron, L.; Zhu, H.; Yuan, J., Ich-1, an Ice/ced-3-related gene, encodes both positive and negative regulators of programmed cell death. Cell 1994, 78 (5), 739–750.

4. Chun, H. J.; Zheng, L.; Ahmad, M.; Wang, J.; Speirs, C. K.; Siegel, R. M.; Dale, J. K.; Puck, J.; Davis, J.; Hall, C. G.; Skoda-Smith, S.; Atkinson, T. P.; Straus, S. E.; Lenardo, M. J., Pleiotropic defects in lymphocyte activation caused by caspase-8 mutations lead to human immunodeficiency. Nature 2002, 419 (6905), 395–399.

5. Alam, A.; Cohen, L. Y.; Aouad, S.; Sékaly, R.-P., Early Activation of Caspases during T Lymphocyte Stimulation Results in Selective Substrate Cleavage in Nonapoptotic Cells. The Journal of Experimental Medicine 1999, 190 (12), 1879–1890.

6. Fujita, J.; Crane, A. M.; Souza, M. K.; Dejosez, M.; Kyba, M.; Flavell, R. A.; Thomson, J. A.; Zwaka, T. P., Caspase activity mediates the differentiation of embryonic stem cells. Cell Stem Cell 2008, 2 (6), 595–601.

7. Baena-Lopez, L. A.; Arthurton, L.; Xu, D. C.; Galasso, A., Non-apoptotic Caspase regulation of stem cell properties. Semin Cell Dev Biol 2018, 82, 118–126.

8. Madadi, Z.; Akbari-Birgani, S.; Monfared, P. D.; Mohammadi, S., The non-apoptotic role of caspase-9 promotes differentiation in leukemic cells. Biochimica Et Biophysica Acta. Molecular Cell Research 2019, 1866 (12), 118524.

9. Sakthivel, D.; Brown-Suedel, A. N.; Keane, F.; Huang, S.; Sherry, K. M.; Charendoff, C. I.; Dunne, K. P.; Robichaux, D. J.; Le, B.; Shin, C. S.; Carisey, A. F.; Flanagan, J. M.; Bouchier-Hayes, L., Caspase-2 is essential for proliferation and self-renewal of nucleophosmin-mutated acute myeloid leukemia. bioRxiv: The Preprint Server for Biology 2023, 2023.05.29.542723.

10. Thornberry, N. A.; Bull, H. G.; Calaycay, J. R.; Chapman, K. T.; Howard, A. D.; Kostura, M. J.; Miller, D. K.; Molineaux, S. M.; Weidner, J. R.; Aunins, J.; Elliston, K. O.; Ayala, J. M.; Casano, F. J.; Chin, J.; Ding, G. J.-F.; Egger, L. A.; Gaffney, E. P.; Limjuco, G.; Palyha, O. C.; Raju, S. M.; Rolando, A. M.; Salley, J. P.; Yamin, T.-T.; Lee, T. D.; Shively, J. E.; MacCross, M.; Mumford, R. A.; Schmidt, J. A.; Tocci, M. J., A novel heterodimeric cysteine protease is required for interleukin-1βprocessing in monocytes. Nature 1992, 356 (6372), 768–774.

11. Salmena, L.; Lemmers, B.; Hakem, A.; Matysiak-Zablocki, E.; Murakami, K.; Au, P. Y. B.; Berry, D. M.; Tamblyn, L.; Shehabeldin, A.; Migon, E.; Wakeham, A.; Bouchard, D.; Yeh, W. C.; McGlade, J. C.; Ohashi, P. S.; Hakem, R., Essential role for caspase 8 in T-cell homeostasis and T-cell-mediated immunity. Genes & Development 2003, 17 (7), 883–895.

12. Wang, J.; Zheng, L.; Lobito, A.; Chan, F. K.-M.; Dale, J.; Sneller, M.; Yao, X.; Puck, J. M.; Straus, S. E.; Lenardo, M. J., Inherited Human Caspase 10 Mutations Underlie Defective Lymphocyte and Dendritic Cell Apoptosis in Autoimmune Lymphoproliferative Syndrome Type II. Cell 1999, 98 (1), 47–58.

13. Boice, A. G.; Lopez, K. E.; Pandita, R. K.; Parsons, M. J.; Charendoff, C. I.; Charaka, V.; Carisey, A. F.; Pandita, T. K.; Bouchier-Hayes, L., Caspase-2 regulates S-phase cell cycle events to protect from DNA damage accumulation independent of apoptosis. Oncogene 2022, 41 (2), 204–219.

14. Martinon, F.; Tschopp, J., Inflammatory caspases and inflammasomes: master switches of inflammation. Cell Death & Differentiation 2007, 14 (1), 10–22.

15. Oliver, T. G.; Meylan, E.; Chang, G. P.; Xue, W.; Burke, J. R.; Humpton, T. J.; Hubbard, D.; Bhutkar, A.; Jacks, T., Caspase-2-mediated cleavage of Mdm2 creates a p53-induced positive feedback loop. Molecular Cell 2011, 43 (1), 57–71.

16. Terry, M. R.; Arya, R.; Mukhopadhyay, A.; Berrett, K. C.; Clair, P. M.; Witt, B.; Salama, M. E.; Bhutkar, A.; Oliver, T. G., Caspase-2 impacts lung tumorigenesis and chemotherapy response in vivo. Cell Death & Differentiation 2015, 22 (5), 719–730.

17. Eskandari, E.; Eaves, C. J., Paradoxical roles of caspase-3 in regulating cell survival, proliferation, and tumorigenesis. Journal of Cell Biology 2022, 221 (6).

18. Rissman, R. A.; Poon, W. W.; Blurton-Jones, M.; Oddo, S.; Torp, R.; Vitek, M. P.; LaFerla, F. M.; Rohn, T. T.; Cotman, C. W., Caspase-cleavage of tau is an early event in Alzheimer disease tangle pathology. J Clin Invest 2004, 114 (1), 121–30.

19. Fuentes-Prior, P.; Salvesen, Guy S., The protein structures that shape caspase activity, specificity, activation and inhibition. Biochemical Journal 2004, 384 (Pt 2), 201–232.

20. Chéreau, D.; Kodandapani, L.; Tomaselli, K. J.; Spada, A. P.; Wu, J. C., Structural and Functional Analysis of Caspase Active Sites. Biochemistry 2003, 42 (14), 4151–4160.

21. McLuskey, K.; Mottram, Jeremy C., Comparative structural analysis of the caspase family with other clan CD cysteine peptidases. Biochemical Journal 2015, 466 (Pt 2), 219–232.

22. Julien, O.; Wells, J. A., Caspases and their substrates. Cell Death and Differentiation 2017, 24 (8), 1380–1389.

23. Lee, B. L.; Mirrashidi, K. M.; Stowe, I. B.; Kummerfeld, S. K.; Watanabe, C.; Haley, B.; Cuellar, T. L.; Reichelt, M.; Kayagaki, N., ASC- and caspase-8-dependent apoptotic pathway diverges from the NLRC4 inflammasome in macrophages. Scientific Reports 2018, 8 (1), 3788.

24. Zheng, T. S.; Flavell, R. A., Divinations and Surprises: Genetic Analysis of Caspase Function in Mice. Experimental Cell Research 2000, 256 (1), 67–73.

25. Hartung, I. V.; Rudolph, J.; Mader, M. M.; Mulder, M. P. C.; Workman, P., Expanding Chemical Probe Space: Quality Criteria for Covalent and Degrader Probes. Journal of Medicinal Chemistry 2023, 66 (14), 9297–9312.

26. Garbaccio, Robert M.; Parmee, Emma R., The Impact of Chemical Probes in Drug Discovery: A Pharmaceutical Industry Perspective. Cell Chemical Biology 2016, 23 (1), 10–17.

27. Garcia-Calvo, M.; Peterson, E. P.; Leiting, B.; Ruel, R.; Nicholson, D. W.; Thornberry, N. A., Inhibition of human caspases by peptide-based and macromolecular inhibitors. The Journal of Biological Chemistry 1998, 273 (49), 32608–32613.

28. Nicholson, D. W.; Ali, A.; Thornberry, N. A.; Vaillancourt, J. P.; Ding, C. K.; Gallant, M.; Gareau, Y.; Griffin, P. R.; Labelle, M.; Lazebnik, Y. A., Identification and inhibition of the ICE/CED-3 protease necessary for mammalian apoptosis. Nature 1995, 376 (6535), 37–43.

29. Enari, M.; Hug, H.; Nagata, S., Involvement of an ICE-like protease in Fas-mediated apoptosis. Nature 1995, 375 (6526), 78–81.

30. Slee, E. A.; Zhu, H.; Chow, S. C.; MacFarlane, M.; Nicholson, D. W.; Cohen, G. M., Benzyloxycarbonyl-Val-Ala-Asp (OMe) fluoromethylketone (Z-VAD.FMK) inhibits apoptosis by blocking the processing of CPP32. The Biochemical Journal 1996, 315 (Pt 1) (Pt 1), 21–24.

31. Thornberry, N. A.; Peterson, E. P.; Zhao, J. J.; Howard, A. D.; Griffin, P. R.; Chapman, K. T., Inactivation of Interleukin-1.beta. Converting Enzyme by Peptide (Acyloxy)methyl Ketones. Biochemistry 1994, 33 (13), 3934–3940.

32. Kato, D.; Boatright, K. M.; Berger, A. B.; Nazif, T.; Blum, G.; Ryan, C.; Chehade, K. A. H.; Salvesen, G. S.; Bogyo, M., Activity-based probes that target diverse cysteine protease families. Nature Chemical Biology 2005, 1 (1), 33–38.

33. Backus, K. M.; Correia, B. E.; Lum, K. M.; Forli, S.; Horning, B. D.; Gonzalez-Paez, G. E.; Chatterjee, S.; Lanning, B. R.; Teijaro, J. R.; Olson, A. J.; Wolan, D. W.; Cravatt, B. F., Proteome-wide covalent ligand discovery in native biological systems. Nature 2016, 534 (7608), 570–4.

34. Cuellar, M. E.; Yang, M.; Karavadhi, S.; Zhang, Y.-Q.; Zhu, H.; Sun, H.; Shen, M.; Hall, M. D.; Patnaik, S.; Ashe, K. H.; Walters, M. A.; Pockes, S., An electrophilic fragment screening for the development of small molecules targeting caspase-2. European Journal of Medicinal Chemistry 2023, 259, 115632.

35. Talanian, R. V.; Quinlan, C.; Trautz, S.; Hackett, M. C.; Mankovich, J. A.; Banach, D.; Ghayur, T.; Brady, K. D.; Wong, W. W., Substrate Specificities of Caspase Family Proteases *. Journal of Biological Chemistry 1997, 272 (15), 9677–9682.

36. Pop, C.; Salvesen, G. S., Human Caspases: Activation, Specificity, and Regulation *. Journal of Biological Chemistry 2009, 284 (33), 21777–21781.

37. Xu, J. H.; Eberhardt, J.; Hill-Payne, B.; González-Páez, G. E.; Castellón, J. O.; Cravatt, B. F.; Forli, S.; Wolan, D. W.; Backus, K. M., Integrative X-ray Structure and Molecular Modeling for the Rationalization of Procaspase-8 Inhibitor Potency and Selectivity. ACS chemical biology 2020, 15 (2), 575–586.

38. Hardy, J. A.; Lam, J.; Nguyen, J. T.; O’Brien, T.; Wells, J. A., Discovery of an allosteric site in the caspases. Proceedings of the National Academy of Sciences 2004, 101 (34), 12461–12466.

39. Tubeleviciute-Aydin, A.; Beautrait, A.; Lynham, J.; Sharma, G.; Gorelik, A.; Deny, L. J.; Soya, N.; Lukacs, G. L.; Nagar, B.; Marinier, A.; LeBlanc, A. C., Identification of Allosteric Inhibitors against Active Caspase-6. Sci Rep 2019, 9 (1), 5504.

40. Vance, N. R.; Gakhar, L.; Spies, M. A., Allosteric Tuning of Caspase-7: A Fragment-Based Drug Discovery Approach. 2017, 56 (46), 14443–14447.

41. Scheer, J. M.; Romanowski, M. J.; Wells, J. A., A common allosteric site and mechanism in caspases. Proceedings of the National Academy of Sciences 2006, 103 (20), 7595–7600.

42. Palafox, M. F.; Desai, H. S.; Arboleda, V. A.; Backus, K. M., From chemoproteomic-detected amino acids to genomic coordinates: insights into precise multi-omic data integration. Molecular Systems Biology 2021, 17 (2), e9840.

43. Jiang, Z. L.; Fletcher, N. M.; Diamond, M. P.; Abu-Soud, H. M.; Saed, G. M., S-nitrosylation of caspase-3 is the mechanism by which adhesion fibroblasts manifest lower apoptosis. Wound Repair and Regeneration 2009, 17 (2), 224–229.

44. Van Horn, K. S.; Wang, D.; Medina-Cleghorn, D.; Lee, P. S.; Bryant, C.; Altobelli, C.; Jaishankar, P.; Leung, K. K.; Ng, R. A.; Ambrose, A. J.; Tang, Y.; Arkin, M. R.; Renslo, A. R., Engaging a Non-catalytic Cysteine Residue Drives Potent and Selective Inhibition of Caspase-6. Journal of the American Chemical Society 2023, 145 (18), 10015–10021.

45. Baliga, B. C.; Read, S. H.; Kumar, S., The biochemical mechanism of caspase-2 activation. Cell Death & Differentiation 2004, 11 (11), 1234–1241.

46. Olsson, M.; Forsberg, J.; Zhivotovsky, B., Caspase-2: the reinvented enzyme. Oncogene 2015, 34 (15), 1877–1882.

47. Bouchier-Hayes, L.; Green, D. R., Caspase-2: the orphan caspase. Cell Death Differ 2012, 19 (1), 51–7.

48. Lim, Y.; Dorstyn, L.; Kumar, S., The p53-caspase-2 axis in the cell cycle and DNA damage response. Experimental & Molecular Medicine 2021, 53 (4), 517–527.

49. Sohn, D.; Budach, W.; Jänicke, R. U., Caspase-2 is required for DNA damage-induced expression of the CDK inhibitor p21(WAF1/CIP1). Cell Death Differ 2011, 18 (10), 1664–74.

50. Oliver, T. G.; Meylan, E.; Chang, G. P.; Xue, W.; Burke, J. R.; Humpton, T. J.; Hubbard, D.; Bhutkar, A.; Jacks, T., Caspase-2-mediated cleavage of Mdm2 creates a p53-induced positive feedback loop. Mol Cell 2011, 43 (1), 57–71.

51. Tinel, A.; Tschopp, J., The PIDDosome, a protein complex implicated in activation of caspase-2 in response to genotoxic stress. Science 2004, 304 (5672), 843–6.

52. Wilson, C. H.; Nikolic, A.; Kentish, S. J.; Keller, M.; Hatzinikolas, G.; Dorstyn, L.; Page, A. J.; Kumar, S., Caspase-2 deficiency enhances whole-body carbohydrate utilisation and prevents high-fat diet-induced obesity. Cell Death & Disease 2017, 8 (10), e3136.

53. Kim, J. Y.; Garcia-Carbonell, R.; Yamachika, S.; Zhao, P.; Dhar, D.; Loomba, R.; Kaufman, R. J.; Saltiel, A. R.; Karin, M., ER Stress Drives Lipogenesis and Steatohepatitis via Caspase-2 Activation of S1P. Cell 2018, 175 (1), 133–145.e15.

54. Dawar, S.; Lim, Y.; Puccini, J.; White, M.; Thomas, P.; Bouchier-Hayes, L.; Green, D. R.; Dorstyn, L.; Kumar, S., Caspase-2-mediated cell death is required for deleting aneuploid cells. Oncogene 2017, 36 (19), 2704–2714.

55. Fava, L. L.; Schuler, F.; Sladky, V.; Haschka, M. D.; Soratroi, C.; Eiterer, L.; Demetz, E.; Weiss, G.; Geley, S.; Nigg, E. A.; Villunger, A., The PIDDosome activates p53 in response to supernumerary centrosomes. Genes & Development 2017, 31 (1), 34–45.

56. Ho, L. H.; Taylor, R.; Dorstyn, L.; Cakouros, D.; Bouillet, P.; Kumar, S., A tumor suppressor function for caspase-2. Proceedings of the National Academy of Sciences of the United States of America 2009, 106 (13), 5336–5341.

57. Zhao, X.; Kotilinek, L. A.; Smith, B.; Hlynialuk, C.; Zahs, K.; Ramsden, M.; Cleary, J.; Ashe, K. H., Caspase-2 cleavage of tau reversibly impairs memory. Nature Medicine 2016, 22 (11), 1268–1276.

58. Machado, M. V.; Michelotti, G. A.; Pereira Tde, A.; Boursier, J.; Kruger, L.; Swiderska-Syn, M.; Karaca, G.; Xie, G.; Guy, C. D.; Bohinc, B.; Lindblom, K. R.; Johnson, E.; Kornbluth, S.; Diehl, A. M., Reduced lipoapoptosis, hedgehog pathway activation and fibrosis in caspase-2 deficient mice with non-alcoholic steatohepatitis. Gut 2015, 64 (7), 1148–57.

59. Kapust, R. B.; Tözsér, J.; Copeland, T. D.; Waugh, D. S., The P1’ specificity of tobacco etch virus protease. Biochemical and biophysical research communications 2002, 294 (5), 949–955.

60. Weerapana, E.; Wang, C.; Simon, G. M.; Richter, F.; Khare, S.; Dillon, M. B. D.; Bachovchin, D. A.; Mowen, K.; Baker, D.; Cravatt, B. F., Quantitative reactivity profiling predicts functional cysteines in proteomes. Nature 2010, 468 (7325), 790–795.

61. Weerapana, E.; Speers, A. E.; Cravatt, B. F., Tandem orthogonal proteolysis-activity-based protein profiling (TOP-ABPP)--a general method for mapping sites of probe modification in proteomes. Nature Protocols 2007, 2 (6), 1414–1425.

62. Abraham, R. T.; Weiss, A., Jurkat T cells and development of the T-cell receptor signalling paradigm. Nat Rev Immunol 2004, 4 (4), 301–8.

63. Saba, J.; Bonneil, E.; Pomiès, C.; Eng, K.; Thibault, P., Enhanced sensitivity in proteomics experiments using FAIMS coupled with a hybrid linear ion trap/Orbitrap mass spectrometer. J Proteome Res 2009, 8 (7), 3355–66.

64. Hebert, A. S.; Prasad, S.; Belford, M. W.; Bailey, D. J.; McAlister, G. C.; Abbatiello, S. E.; Huguet, R.; Wouters, E. R.; Dunyach, J.-J.; Brademan, D. R.; Westphall, M. S.; Coon, J. J., Comprehensive Single-Shot Proteomics with FAIMS on a Hybrid Orbitrap Mass Spectrometer. Analytical Chemistry 2018, 90 (15), 9529–9537.

65. Arad, D.; Langridge, R.; Kollman, P. A., A simulation of the sulfur attack in catalytic pathway of papain using molecular mechanics and semiempirical quantum mechanics. Journal of the American Chemical Society 1990, 112 (2), 491–502.

66. Itoh, N.; Yonehara, S.; Ishii, A.; Yonehara, M.; Mizushima, S.-I.; Sameshima, M.; Hase, A.; Seto, Y.; Nagata, S., The polypeptide encoded by the cDNA for human cell surface antigen Fas can mediate apoptosis. Cell 1991, 66 (2), 233–243.

67. Suda, T.; Takahashi, T.; Golstein, P.; Nagata, S., Molecular cloning and expression of the Fas ligand, a novel member of the tumor necrosis factor family. Cell 1993, 75 (6), 1169–78.

68. Yang, J.; Yan, R.; Roy, A.; Xu, D.; Poisson, J.; Zhang, Y., The I-TASSER Suite: protein structure and function prediction. Nature Methods 2015, 12 (1), 7–8.

69. Elliott, J. M.; Rouge, L.; Wiesmann, C.; Scheer, J. M., Crystal Structure of Procaspase-1 Zymogen Domain Reveals Insight into Inflammatory Caspase Autoactivation. Journal of Biological Chemistry 2009, 284 (10), 6546–6553.

70. Chai, J.; Wu, Q.; Shiozaki, E.; Srinivasula, S. M.; Alnemri, E. S.; Shi, Y., Crystal Structure of a Procaspase-7 Zymogen: Mechanisms of Activation and Substrate Binding. Cell 2001, 107 (3), 399–407.

71. Thomsen, N. D.; Koerber, J. T.; Wells, J. A. J. P. o. t. N. A. o. S., Structural snapshots reveal distinct mechanisms of procaspase-3 and-7 activation. 2013, 110 (21), 8477–8482.

72. Hughes, C. S.; Moggridge, S.; Müller, T.; Sorensen, P. H.; Morin, G. B.; Krijgsveld, J., Single-pot, solid-phase-enhanced sample preparation for proteomics experiments. Nature Protocols 2019, 14 (1), 68–85.

73. Palafox, M. F.; Desai, H. S.; Arboleda, V. A.; Backus, K. M., From chemoproteomic-detected amino acids to genomic coordinates: insights into precise multi-omic data integration. Mol Syst Biol 2021, 17 (2), e9840.

74. Vickers, C. J.; González-Páez, G. E.; Wolan, D. W., Selective detection and inhibition of active caspase-3 in cells with optimized peptides. J Am Chem Soc 2013, 135 (34), 12869–76.

75. Salvesen, G. S.; Dixit, V. M., Caspase activation: the induced-proximity model. Proc Natl Acad Sci U S A 1999, 96 (20), 10964–7.

76. Kruspig, B.; Nilchian, A.; Orrenius, S.; Zhivotovsky, B.; Gogvadze, V., Citrate kills tumor cells through activation of apical caspases. Cell Mol Life Sci 2012, 69 (24), 4229–37.

77. Boatright, K. M.; Renatus, M.; Scott, F. L.; Sperandio, S.; Shin, H.; Pedersen, I. M.; Ricci, J. E.; Edris, W. A.; Sutherlin, D. P.; Green, D. R.; Salvesen, G. S., A unified model for apical caspase activation. Mol Cell 2003, 11 (2), 529–41.

78. Ito, A.; Uehara, T.; Tokumitsu, A.; Okuma, Y.; Nomura, Y., Possible involvement of cytochrome c release and sequential activation of caspases in ceramide-induced apoptosis in SK-N-MC cells. Biochimica Et Biophysica Acta 1999, 1452 (3), 263–274.

79. Cao, J.; Boatner, L. M.; Desai, H. S.; Burton, N. R.; Armenta, E.; Chan, N. J.; Castellón, J. O.; Backus, K. M., Multiplexed CuAAC Suzuki-Miyaura Labeling for Tandem Activity-Based Chemoproteomic Profiling. Anal Chem 2021, 93 (4), 2610–2618.

80. Boatner, L. M.; Palafox, M. F.; Schweppe, D. K.; Backus, K. M., CysDB: a human cysteine database based on experimental quantitative chemoproteomics. Cell Chem Biol 2023, 30 (6), 683–698.e3.

81. Yan, T.; Desai, H. S.; Boatner, L. M.; Yen, S. L.; Cao, J.; Palafox, M. F.; Jami-Alahmadi, Y.; Backus, K. M., SP3-FAIMS Chemoproteomics for High-Coverage Profiling of the Human Cysteinome*. Chembiochem 2021, 22 (10), 1841–1851.

82. Hebert, A. S.; Prasad, S.; Belford, M. W.; Bailey, D. J.; McAlister, G. C.; Abbatiello, S. E.; Huguet, R.; Wouters, E. R.; Dunyach, J. J.; Brademan, D. R.; Westphall, M. S.; Coon, J. J., Comprehensive Single-Shot Proteomics with FAIMS on a Hybrid Orbitrap Mass Spectrometer. Anal Chem 2018, 90 (15), 9529–9537.

83. Morgan, C. W.; Julien, O.; Unger, E. K.; Shah, N. M.; Wells, J. A., Turning on caspases with genetics and small molecules. Methods Enzymol 2014, 544, 179–213.

84. Julien, O.; Zhuang, M.; Wiita, A. P.; O’Donoghue, A. J.; Knudsen, G. M.; Craik, C. S.; Wells, J. A., Quantitative MS-based enzymology of caspases reveals distinct protein substrate specificities, hierarchies, and cellular roles. Proc Natl Acad Sci U S A 2016, 113 (14), E2001–10.

85. Gray, D. C.; Mahrus, S.; Wells, J. A., Activation of specific apoptotic caspases with an engineered small-molecule-activated protease. Cell 2010, 142 (4), 637–46.

86. Salvesen, G. S.; Dixit, V. M., Caspase activation: The induced-proximity model. Proceedings of the National Academy of Sciences 1999, 96 (20), 10964–10967.

87. Pozueta, J.; Lefort, R.; Ribe, E. M.; Troy, C. M.; Aran-cio, O.; Shelanski, M., Caspase-2 is required for dendritic spine and behavioral alterations in J20 APP transgenic mice. Nature communications 2013, 4, 1939.

88. Hermel, E.; Gafni, J.; Propp, S. S.; Leavitt, B. R.; Wellington, C. L.; Young, J. E.; Hackam, A. S.; Logvinova, A. V.; Peel, A. L.; Chen, S. F.; Hook, V.; Singaraja, R.; Krajewski, S.; Goldsmith, P. C.; Ellerby, H. M.; Hayden, M. R.; Bredesen, D. E.; Ellerby, L. M., Specific caspase interactions and amplification are involved in selective neuronal vulnerability in Huntington’s disease. Cell Death & Differentiation 2004, 11 (4), 424–438.

89. Bosc, E.; Anastasie, J.; Soualmia, F.; Coric, P.; Kim, J. Y.; Wang, L. Q.; Lacin, G.; Zhao, K.; Patel, R.; Duplus, E.; Tixador, P.; Sproul, A. A.; Brugg, B.; Reboud-Ravaux, M.; Troy, C. M.; Shelanski, M. L.; Bouaziz, S.; Karin, M.; El Amri, C.; Jacotot, E. D., Genuine selective caspase-2 inhibition with new irreversible small peptidomimetics. Cell Death & Disease 2022, 13 (11), 959.

90. Poreba, M.; Rut, W.; Groborz, K.; Snipas, S. J.; Salvesen, G. S.; Drag, M., Potent and selective caspase-2 inhibitor prevents MDM-2 cleavage in reversine-treated colon cancer cells. Cell Death Differ 2019, 26 (12), 2695–2709.

91. Bouchier-Hayes, L.; Martin, S. J., CARD games in apoptosis and immunity. EMBO Rep 2002, 3 (7), 616–21.

92. Nabet, B.; Roberts, J. M.; Buckley, D. L.; Paulk, J.; Dastjerdi, S.; Yang, A.; Leggett, A. L.; Erb, M. A.; Lawlor, M. A.; Souza, A.; Scott, T. G.; Vittori, S.; Perry, J. A.; Qi, J.; Winter, G. E.; Wong, K.-K.; Gray, N. S.; Bradner, J. E., The dTAG system for immediate and target-specific protein degradation. Nature Chemical Biology 2018, 14 (5), 431–441.

93. Zhang, J.; Yang, P. L.; Gray, N. S., Targeting cancer with small molecule kinase inhibitors. Nature Reviews Cancer 2009, 9 (1), 28–39.

94. Zhao, Z.; Bourne, P. E., Progress with covalent small-molecule kinase inhibitors. Drug Discovery Today 2018, 23 (3), 727–735.

95. Abdeldayem, A.; Raouf, Y. S.; Constantinescu, S. N.; Moriggl, R.; Gunning, P. T., Advances in covalent kinase inhibitors. Chemical Society Reviews 2020, 49 (9), 2617–2687.

96. Cravatt, B. F., Activity-Based Protein Profiling – Finding General Solutions to Specific Problems. 2023, 63 (3-4), e202300029.

97. Kemper, E. K.; Zhang, Y.; Dix, M. M.; Cravatt, B. F., Global profiling of phosphorylation-dependent changes in cysteine reactivity. Nat Methods 2022, 19 (3), 341–352.

98. Ostrem, J. M.; Peters, U.; Sos, M. L.; Wells, J. A.; Shokat, K. M., K-Ras(G12C) inhibitors allosterically control GTP affinity and effector interactions. Nature 2013, 503 (7477), 548–51.

99. Dhawan, N. S.; Scopton, A. P.; Dar, A. C., Small molecule stabilization of the KSR inactive state antagonizes oncogenic Ras signalling. Nature 2016, 537 (7618), 112–116.

100. Dai, S. A.; Hu, Q.; Gao, R.; Blythe, E. E.; Touhara, K. K.; Peacock, H.; Zhang, Z.; von Zastrow, M.; Suga, H.; Shokat, K. M., State-selective modulation of heterotrimeric Gαs signaling with macrocyclic peptides. Cell 2022, 185 (21), 3950–3965.e25.

101. Wang, W.; Krosky, D.; Ahn, K., Discovery of Inactive Conformation-Selective Kinase Inhibitors by Utilizing Cascade Assays. Biochemistry 2017, 56 (34), 4449–4456.

102. Ayala-Aguilera, C. C.; Valero, T.; Lorente-Macías, Á.; Baillache, D. J.; Croke, S.; Unciti-Broceta, A., Small Molecule Kinase Inhibitor Drugs (1995–2021): Medical Indication, Pharmacology, and Synthesis. Journal of Medicinal Chemistry 2022, 65 (2), 1047–1131.

103. Schindler, T.; Bornmann, W.; Pellicena, P.; Miller, W. T.; Clarkson, B.; Kuriyan, J., Structural mechanism for STI-571 inhibition of abelson tyrosine kinase. Science 2000, 289 (5486), 1938–42.

104. Tyas, L.; Brophy, V. A.; Pope, A.; Rivett, A. J.; Tavaré, J. M., Rapid caspase-3 activation during apoptosis revealed using fluorescence-resonance energy transfer. EMBO Rep 2000, 1 (3), 266–70.

105. Perez-Riverol, Y.; Csordas, A.; Bai, J.; Bernal-Llinares, M.; Hewapathirana, S.; Kundu, D. J.; Inuganti, A.; Griss, J.; Mayer, G.; Eisenacher, M.; Pérez, E.; Uszkoreit, J.; Pfeuffer, J.; Sachsenberg, T.; Yilmaz, S.; Tiwary, S.; Cox, J.; Audain, E.; Walzer, M.; Jarnuczak, A. F.; Ternent, T.; Brazma, A.; Vizcaíno, J. A., The PRIDE database and related tools and resources in 2019: improving support for quantification data. Nucleic Acids Res 2019, 47 (D1), D442–d450.

