## Supplementary Information for "Chemoproteomics identifies proteoform-selective caspase-2 inhibitors"

**Table of Contents**

| | |
|---|---|
| **(A) Supplementary Figures** | **3 - 31** |
| **(B) Supplementary Tables** | **31 - 38** |
| **(C) Biology Methods** | **38 - 46** |
| **(D) Chemistry Methods** | **46 - 56** |
| **(E) NMR Spectra** | **56 - 78** |
| (F) **References** | **78 - 79** |

### (A) Supplementary Figures

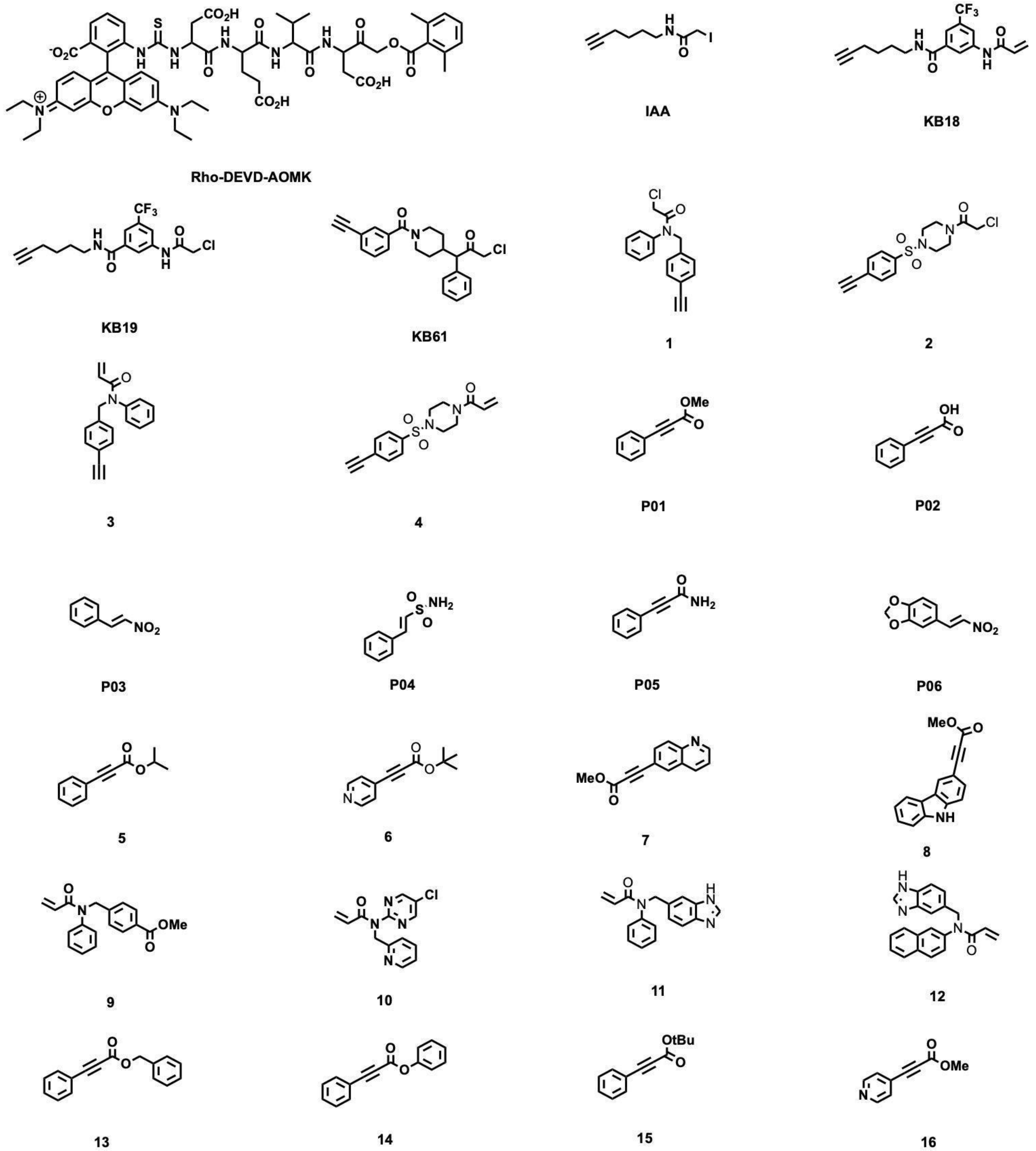

**Figure S1**. **Structures of compounds used in this study**, including previously reported compounds **KB18**, **KB19**, and **KB61** reported by Backus, KM., et al. 2016[1], **Rho-DEVD-AOMK** probe generated by Wolan *et al*.[1] iodoacetamide alkyne (**IAA**) first reported by Weerapana et al. 2010[2] and purchased compounds **P01** (Fisher Scientific, AC334590050), **P02** (Fisher Scientific, P06105G), **P03** (Combi-blocks, QB-5712), (**P04** (Combi-blocks, ST-8644), **P05** (Combi-blocks, QC-2990) and **P06** (Combi-blocks, QF-4549).

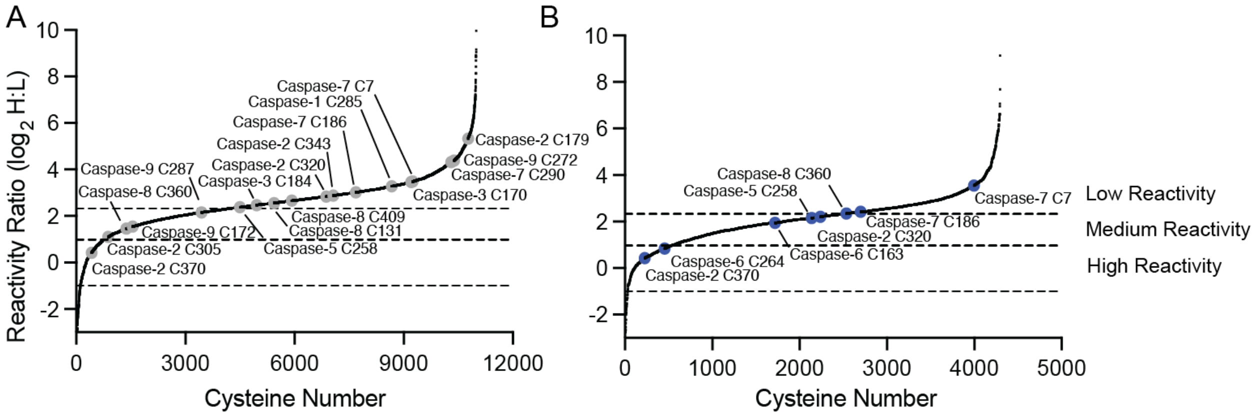

**Figure S2**. **Isotopic tandem orthogonal proteolysis–activity-based protein profiling (isoTOP-ABPP) reactivity analysis of (A) non-apoptotic cell lysates and (B) apoptotic lysates.** Cell lysates obtained for Jurkat cells treated with FasL (50 ng/µL FasL, 4h) and subjected to isoTOP-ABPP ABPP[2,3] comparing concentration-dependent cysteine labeling by **IAA (**10 µM vs 100 µM, 1h) analysis following the workflow shown in **Figure 1A**. IsoTOP-ABPP ratio ($R_{10:1}$) reactivity thresholds, calculated from the MS1 ion intensity ratios: "high" reactivity: $Log_2(R_{heavy:light})$ = -1.0 – 1.0, "medium" reactivity: $Log_2(R_{heavy:light})$ = = 1 - 2.32, "low" reactivity $Log_2(R_{heavy:light})$ = >2.32. Non-apoptotic experiments (n=6). Apoptotic experiments (n = 5). All MS data can be found in **Table S2**.

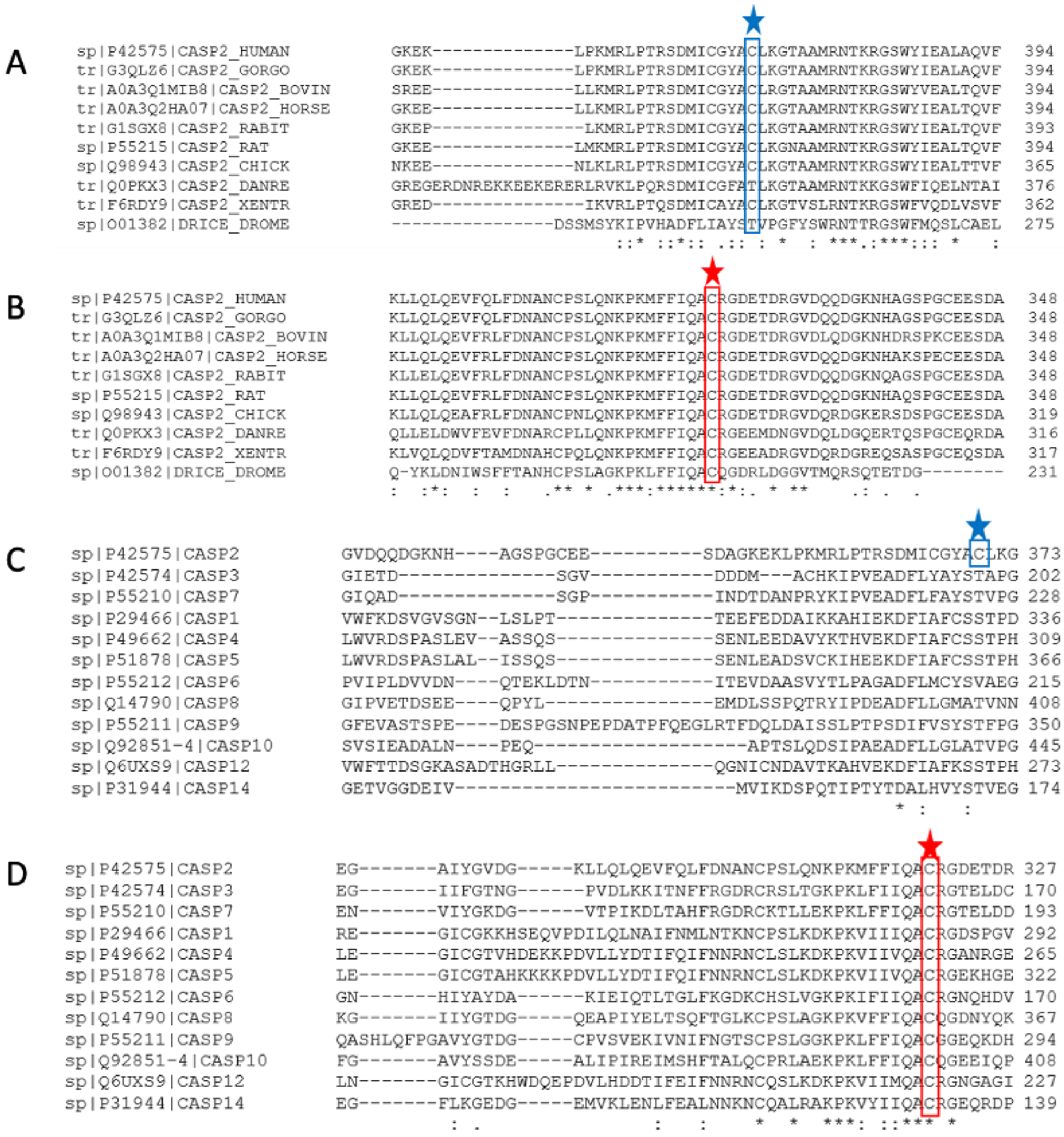

**Figure S3. C370 is highly conserved and unique to caspase-2**. (A,B) Model organism caspase-2 orthologue protein sequence alignments (Clustal Omega[4,5]. (A) C370 in blue and (B) catalytic C320 in red. (C,D) Protein sequence alignment of 12 human caspases. (C) C370 in blue and (D) Catalytic C320 in red.

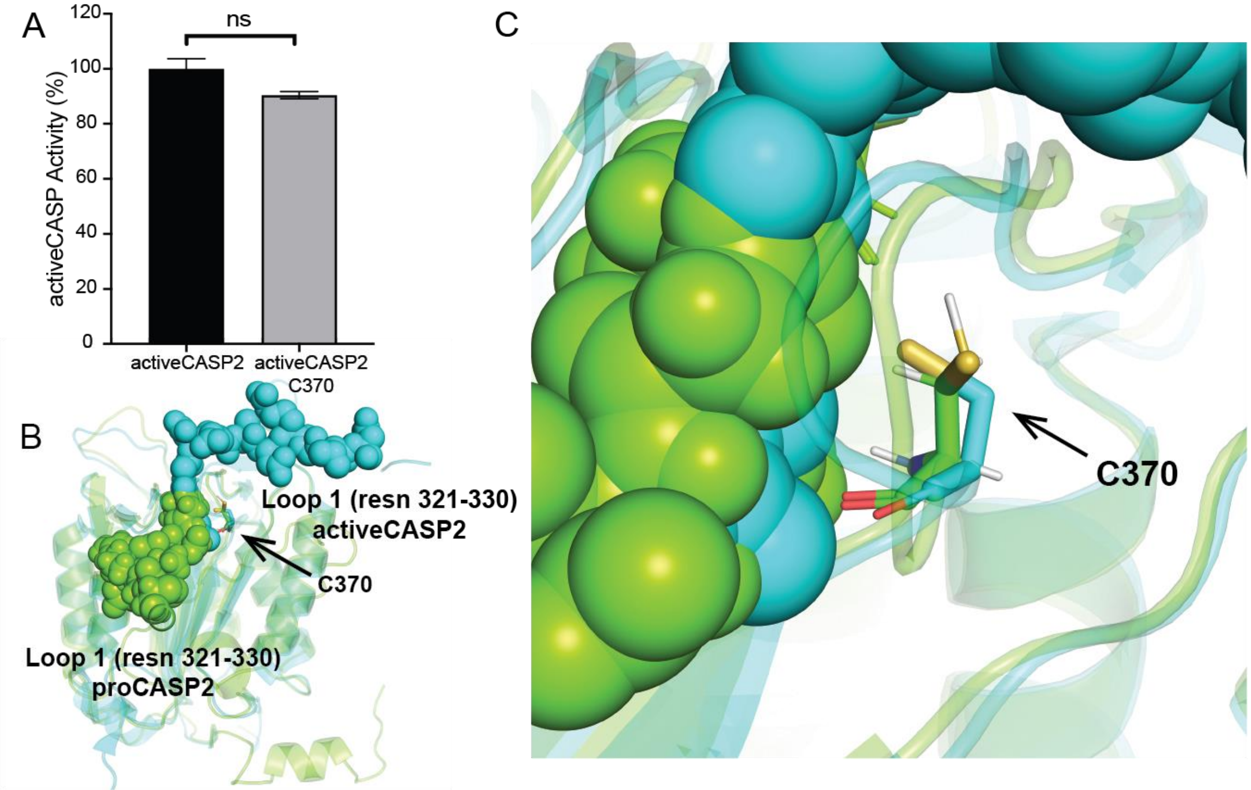

**Figure S4**. **C370 is an active-site proximal non-catalytic cysteine residue.** (A) Comparison of the activity of recombinant activeCASP2 and activeACSP2_C370A (1 µM recombinant protein, 5 mM DTT, in PBS supplemented with 333 mM citrate buffer pH 7.4) using fluorogenic substrate Ac-VDVAD-AFC (10 µM substrate) and fluorescence emission spectra ( $\lambda_{ex}$ = 400 nm and $\lambda_{em}$ = 505 nm) monitored by multimodal plate reader with percentage activity calculated from the linear range of the reaction curves. (B) Crystal structure of active-caspase-2 (PDB: 1PYO; complexed with Acetyl-Leu-Asp-Glu-Ser-Asp-cho) in cyan highlighting active site adjacent loop (Loop 1) overlaid with predicted (I-TASSER[6–8] structure of pro-caspase-2 in green. (C) 180° flip of loop 1 repositions the C370 Sulfhydryl. For A) data represents mean activity ± STDEV for two technical replicate experiments. Statistical significance was calculated with unpaired Student's t-tests, comparing activeCASP2- to activeCASP2_C370A; ns, not significant, n.s. p>0.05.

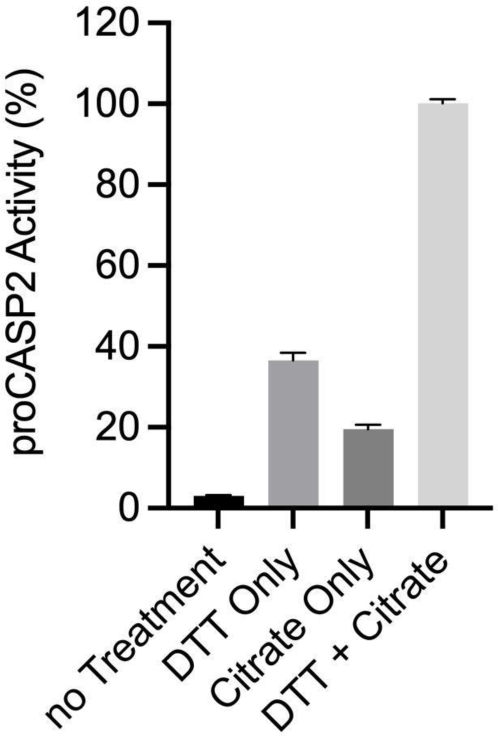

**Figure S5. Assessment of recombinant proCASP2 activity sensitivity to additives.** Additives: DTT (10 mM pre-treatment for 30 mins) and citrate (333 mM added immediately prior to assay). Activity of recombinant proCASP2 (1 µM recombinant protein in PBS supplemented with 333 mM citrate buffer pH 7.4) monitored using fluorogenic substrate Ac-VDVAD-AFC (10 µM substrate) and fluorescence emission spectra ( $\lambda_{ex}$ = 400 nm and $\lambda_{em}$ = 505 nm) monitored by multimodal plate reader with percentage activity relative to DTT+Citrate conditions calculated from the linear range of the reaction curves. Data represent mean values ± STDEV for three technical replicate experiments.

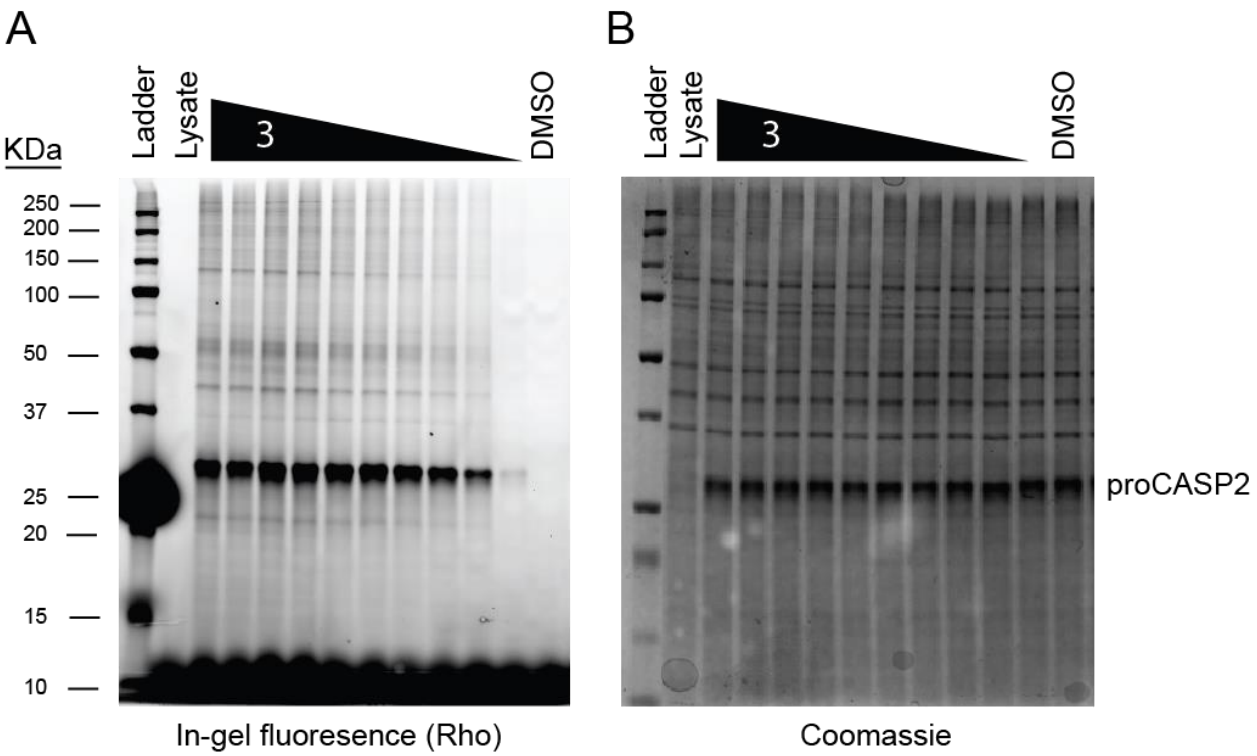

**Figure S6. Dose-dependent labeling of proCASP2 by alkyne probe 3.** Gel-based ABPP analysis of recombinant proCASP2 in cell lysates subjected to labeling a dose range of **3** (1000 μM > 750 μM > 500 μM > 250 μM > 100 μM > 75 μM > 50 μM> 25 μM> 10 μM > 1 μM) for 1h followed by click conjugation to rhodamine azide and SDS-PAGE analysis. (A) In-gel fluorescence and (B) Coomassie InstantBlue visualization of protein loading.

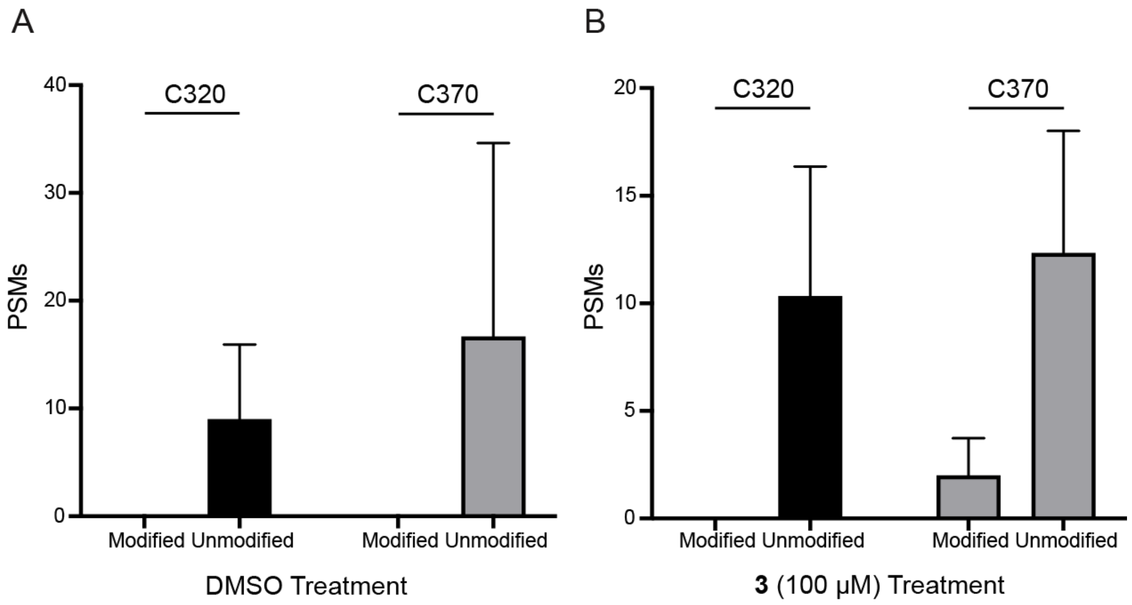

**Figure S7**. **Bottom-up proteomic analysis of recombinant proCASP2 labeled by 3**. Recombinant proCASP2 in PBS was treated with either (A) DMSO vehicle or (B) compound **3** (100 μM final concentration) for 1h at 30℃ followed by sample cleanup (SP3), sequence specific digest and LC-MS/MS analysis. Data represent mean values ± STDEV for three biological replicates. All MS Data can be found in **Table S4.**

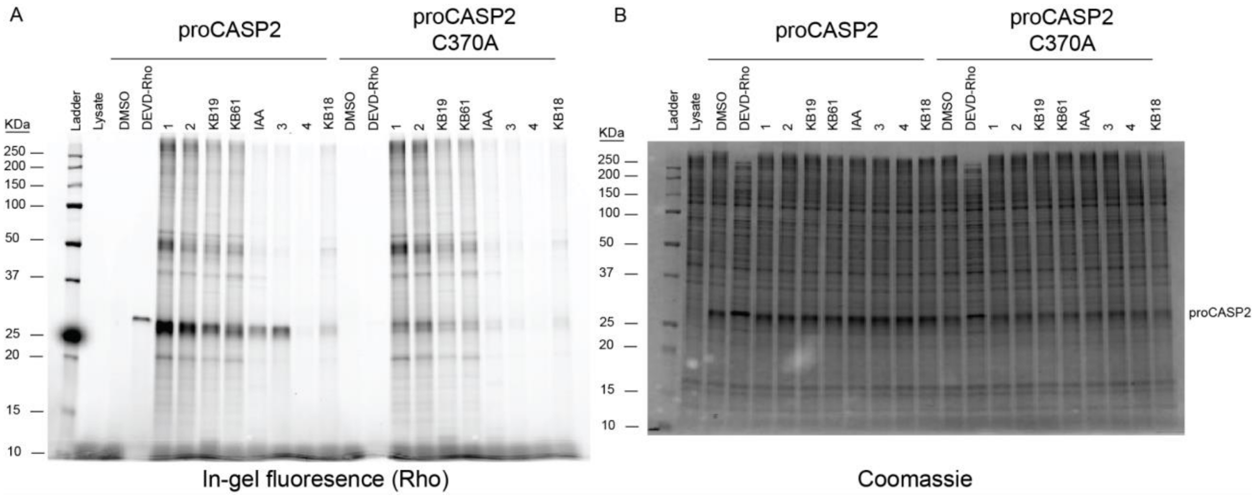

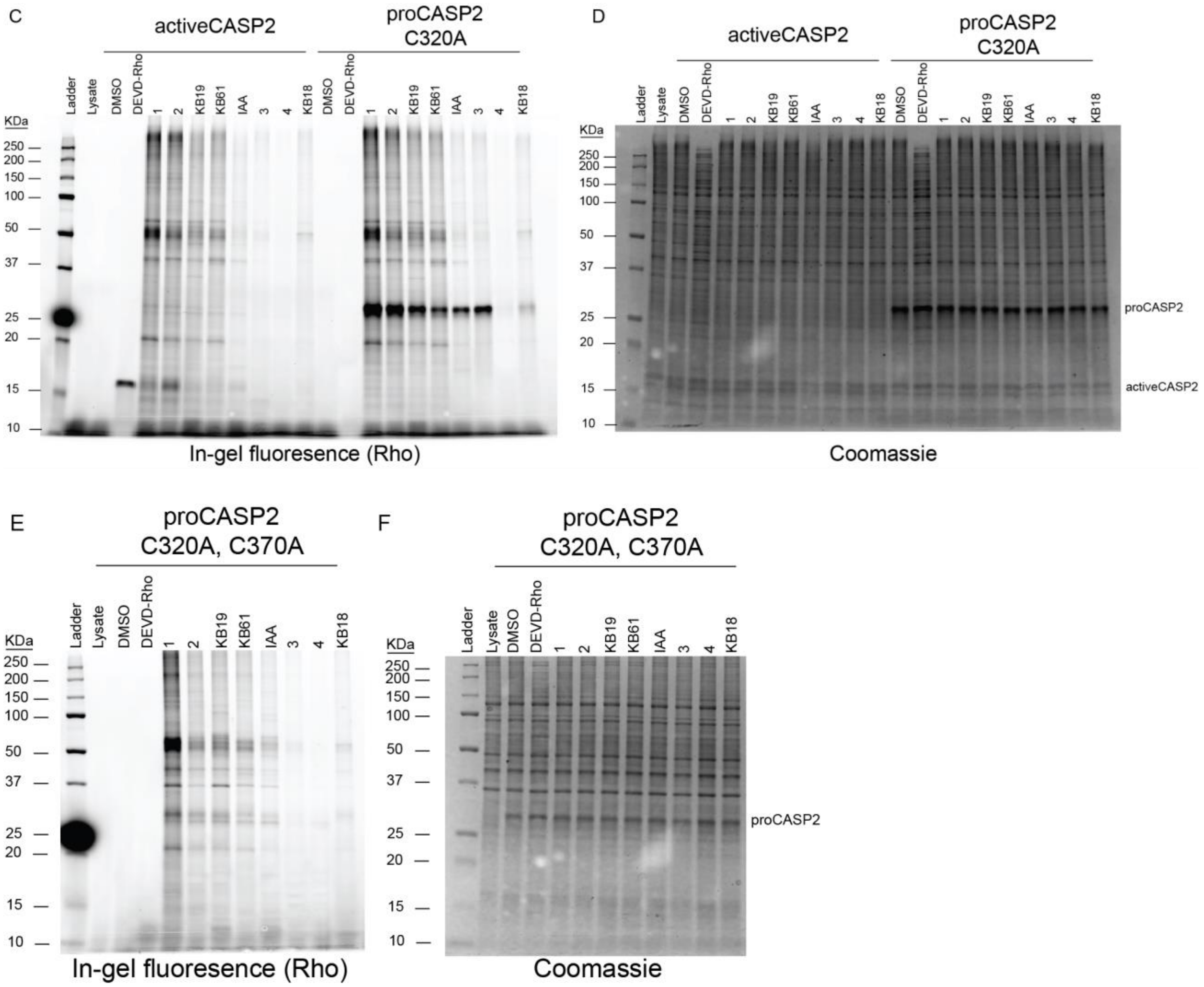

**Figure S8. Full length gels for Figure 2B, labeling of proCASP2 by alkyne probes.** Gel-based ABPP analysis of recombinant caspase-2 constructs harboring the indicated mutations in whole cell lysates treated for 1h with the indicated compounds (10 µM for all click probes, 2 µM final for **Rho-DEVD-AOMK** and 2 µM final for **IAA**), followed by click conjugation to rhodamine azide for alkyne probes and in-gel analysis. (A,C,E) in-gel fluorescence and (B,D,F) Coomassie InstantBlue visualization of protein loading for (A,B) proCASP2 and proCASP2_C370A (C, D) activeCASP2 and proCASP2_C320A, (E, F) proCASP2_C320A,C370A and proCASP2_C320A.

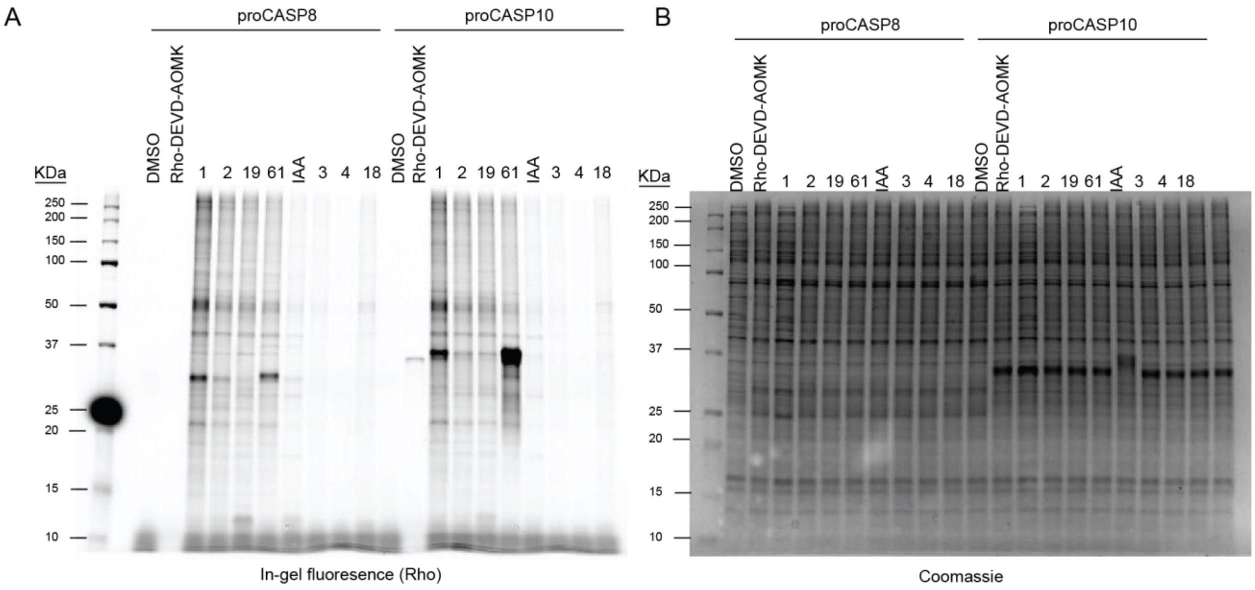

**Figure S9. Full length gels for Figure 2E, labeling of proCASP8 and proCASP10 by alkyne probes**. Gel-based ABPP analysis of recombinant proCASP8 and proCASP10 in Jurkat cell lysates treated for 1h with the indicated compounds (10 µM for all click probes and 2 µM final for **Rho-DEVD-AOMK**), followed by click conjugation to rhodamine azide for alkyne probes and in-gel analysis. (A) in-gel fluorescence and (B) Coomassie InstantBlue visualization of protein loading.

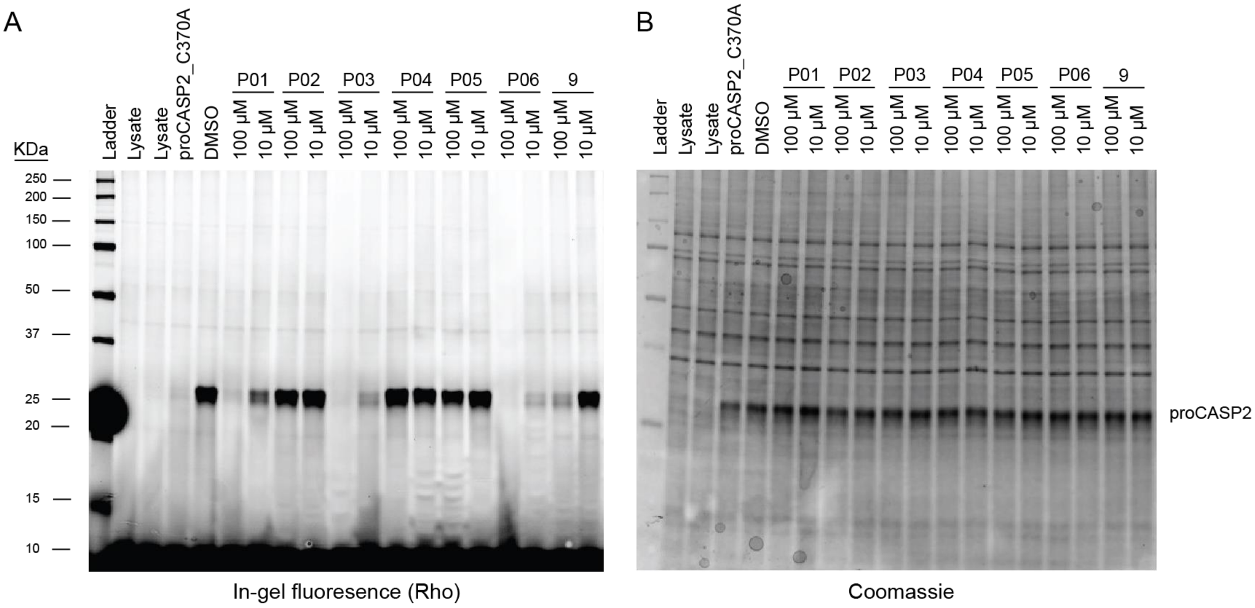

**Figure S10**. **Competitive gel-based ABPP analysis of proCASP2 labeling by diverse electrophilic fragments.** Recombinant proCASP2 in Jurkat cell lysates treated for 1h with the indicated compounds at the indicated concentrations, followed by **3** (10 µM, 1h), click conjugation to rhodamine azide and in-gel analysis. (A) in-gel fluorescence and (B) Coomassie InstantBlue visualization of protein loading.

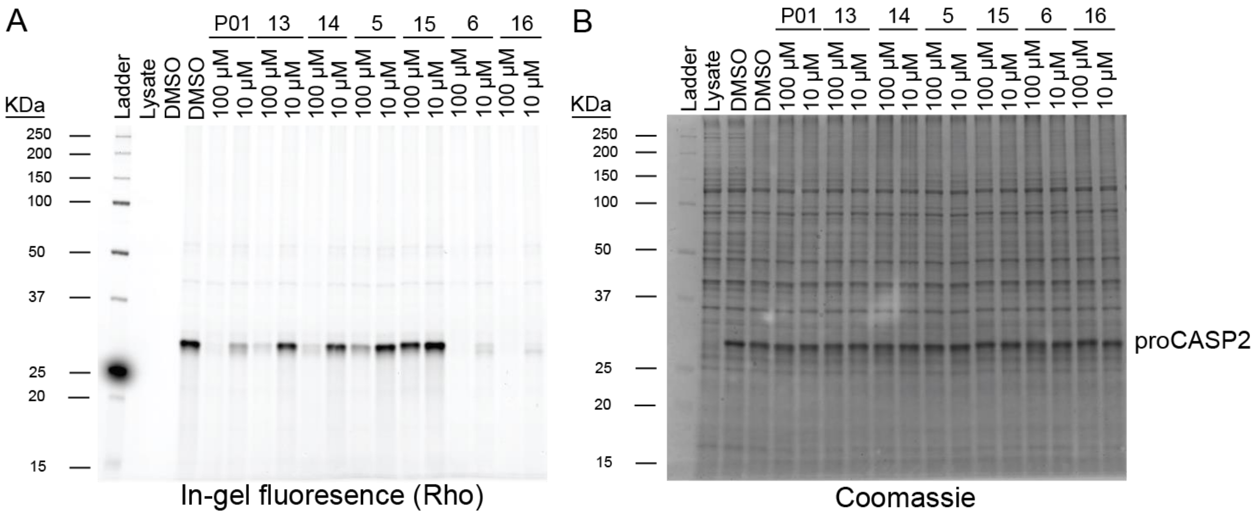

**Figure S11**. **Competitive gel-based ABPP analysis of proCASP2 labeling by methyl phenylpropiolate (P01) analogues.** Recombinant proCASP2 in Jurkat cell lysates treated for 1h with the indicated compounds at the indicated concentrations, followed by **3 (**10 µM, 1h), click conjugation to rhodamine azide and in-gel analysis. (A) in-gel fluorescence and (B) Coomassie InstantBlue visualization of protein loading.

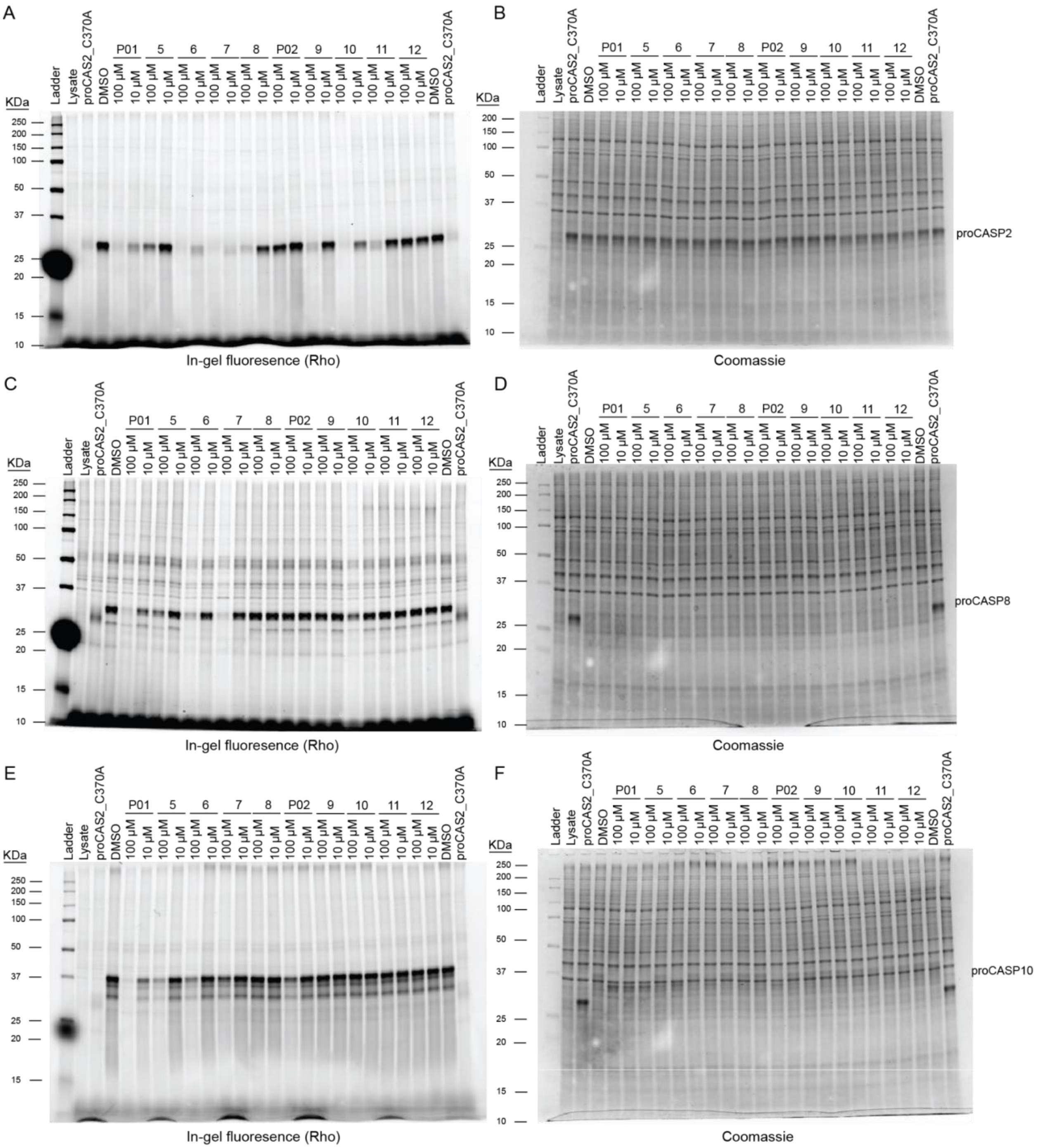

**Figure S12. Competitive gel-based ABPP analysis of procaspases labeling by electrophilic fragments.** Gel-based ABPP analysis of recombinant (A,B) proCASP2, (C,D) proCASP8, and (E,F) proCASP10 in whole cell lysates treated for 1h with the indicated compounds at the indicated concentrations, followed by either **3 (**10 µM, 1h) for proCASP2 or **KB61** (10 µM, 1h) for proCASP8 and proCASP10, click conjugation to rhodamine azide and in-gel analysis. (A,C,E) in-gel fluorescence and (B,D,F) Coomassie InstantBlue visualization of protein loading.

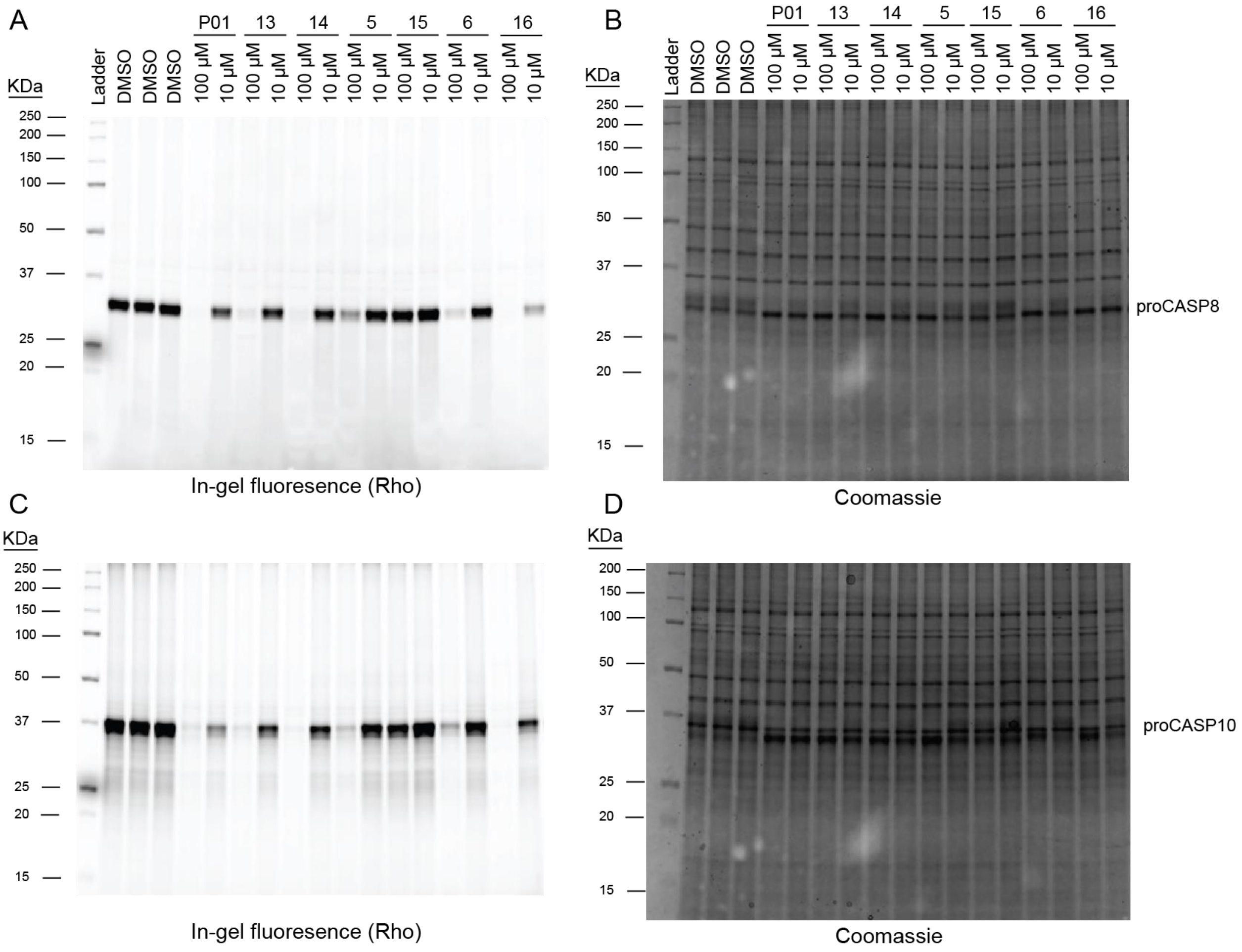

**Figure S13**. **Competitive gel-based ABPP analysis of proCASP8 and proCASP10 labeling by electrophilic fragments.** Gel-based ABPP analysis of recombinant (A,B) proCASP8 and (C,D) proCASP10) in whole cell lysates treated for 1h with the indicated compounds at the indicated concentrations, followed by **KB61** (10 µM, 1h) for proCASP8 and proCASP10, click conjugation to rhodamine azide and in-gel analysis. (A,C) in-gel fluorescence and (B,D) Coomassie InstantBlue visualization of protein loading.

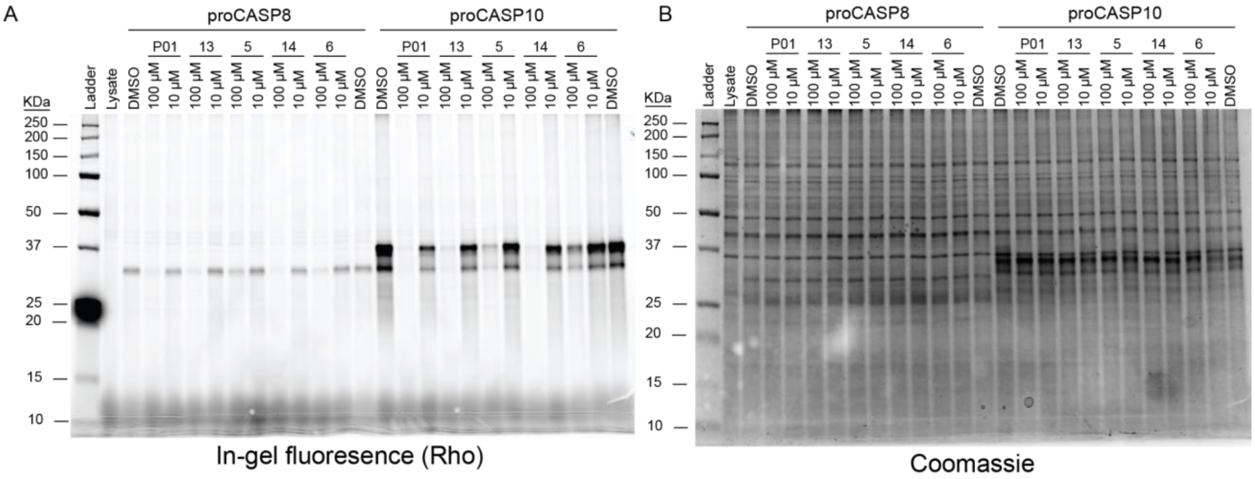

**Figure S14**. **Competitive gel-based ABPP analysis of proCASP8 and proCASP10 labeling by P01 analogues.** The indicated procaspases in whole cell lysates were treated for 1h with the indicated compounds at the indicated concentrations, followed by **KB61** (10 µM, 1h), click conjugation to rhodamine azide and in-gel analysis. (A) in-gel fluorescence and (B) Coomassie InstantBlue visualization of protein loading.

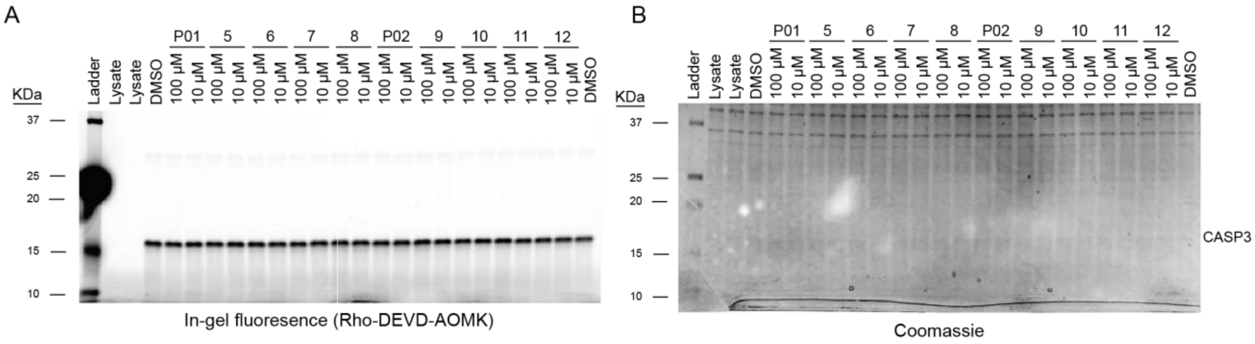

**Figure S15**. **Competitive gel-based ABPP analysis of activeCASP3 labeling by electrophilic fragments.** ActiveCASP3 in whole cell lysates was treated for 1h with the indicated compounds at the indicated concentrations, followed by **Rho-DEVD-AOMK** (0.2 µM), and in-gel analysis. (A) in-gel fluorescence and (B) Coomassie InstantBlue visualization of protein loading.

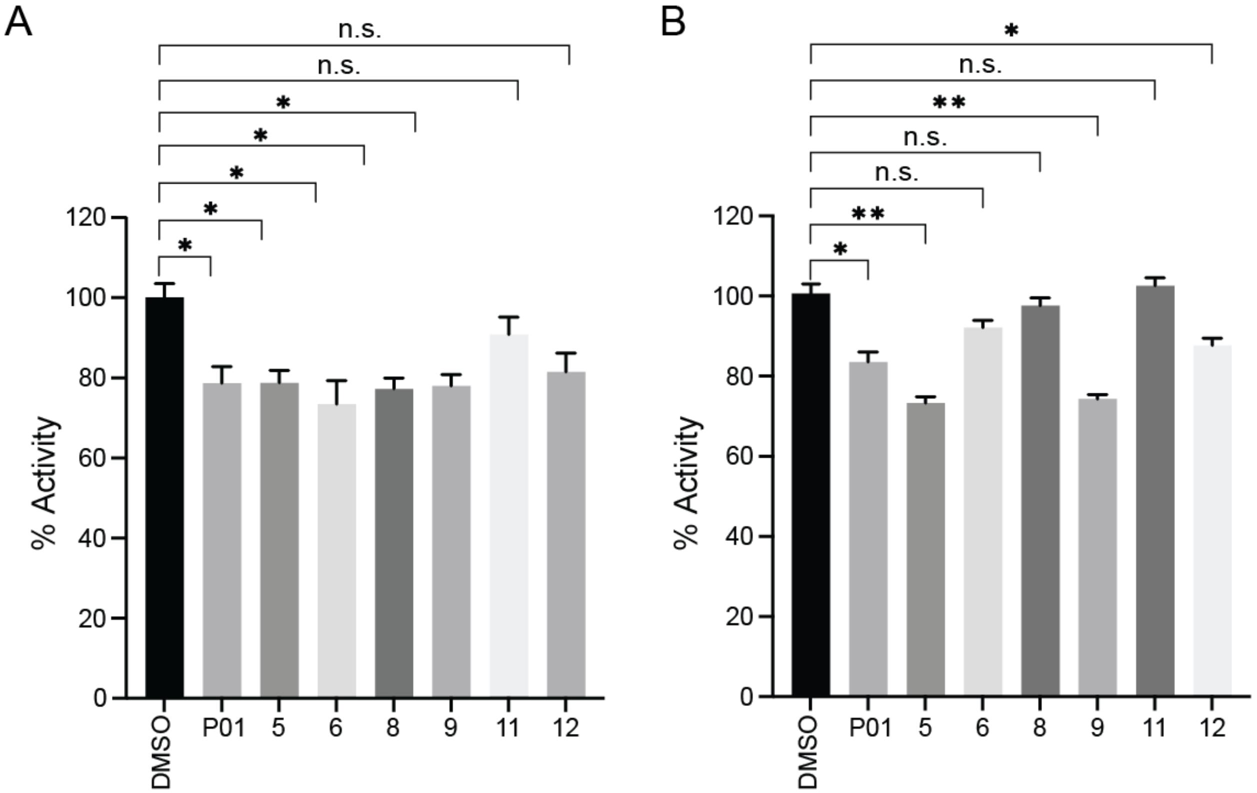

**Figure S16. Assessment of recombinant (A) activeCASP2 and (B) activeCASP2_C370A sensitivity to electrophilic fragments.** Activity of recombinant protein assessed using fluorogenic substrate Ac-VDVAD-AFC (10 μM) with fluorescence emission spectra ($\lambda_{ex}$ = 400 nm and $\lambda_{em}$ = 505 nm) monitored by multimodal plate reader with percentage activity relative to DMSO calculated from the linear range of the reaction curves. Data represent mean values ± STDEV for three technical replicates. Statistical significance was calculated with unpaired Student's t-tests, n.s., not significant, * p<0.05, ** p<0.01, n.s. p>0.05.

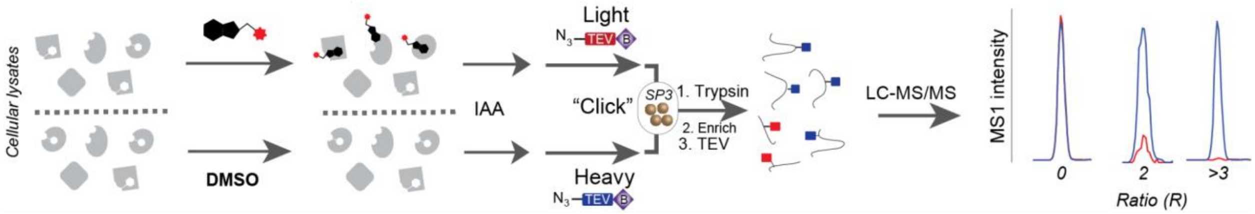

**Figure S17. Workflow for competitive isoTOP-ABPP analysis of electrophilic fragments.** IsoTOP-ABPP workflow used here, which was modified from the previously published method to incorporate single-pot, solid-phase-enhanced sample-preparation (SP3) cleanup[9]. Jurkat lysates are treated with either compound (100 µM) or vehicle (DMSO) followed by **IAA** cysteine capping at 200 µM final concentration for 1h. Samples were subjected to Cu(I)-catalyzed azide–alkyne cycloaddition (CuAAC) conjugation to isotopically labeled tobacco etch virus (TEV)-cleavable biotinylated peptide tags[3]. The samples are then combined, subjected to SP3 cleanup, enrichment on streptavidin and sequential trypsin and TEV digests, the combined samples were acquired by LC-MS/MS and the relative areas for the MS1 chromatographic peaks quantified. IsoTOP-ABPP ratios (R = $\log_2$DMSO:compound) are calculated from the MS1 ion intensity ratios for heavy- versus light-labeled peptides. Competition H:L ratios (R = $\log_2$H:L) ≥ 2 designate fragment electrophile labeling sites.

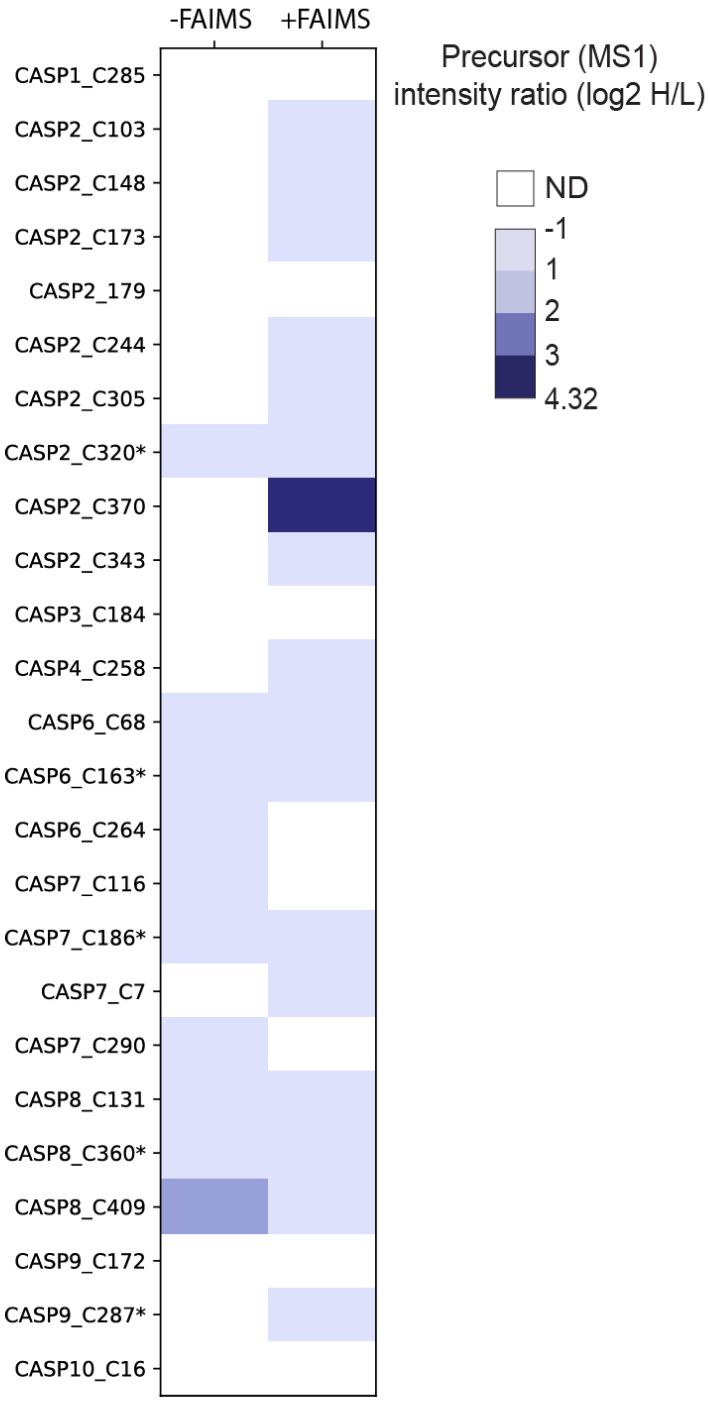

**Figure S18**. **Assessment of impact of FAIMS acquisition on caspase peptide coverage.** Competitive isoTOP-ABPP samples prepared as shown in **Figure S17** with Jurkat lysates treated with **9** (100 μM for 1h) or vehicle (DMSO). Shown are the mean values for precursor ion intensity ratios for biological replicates (n = 2) acquired ± FAIMS. MS Data can be found in **Table S4**. * Indicate catalytic cysteine residues.

```
proCASP2            DNANCPSLQNKPKMFFIQACRGDETDRGVDQQ-----DGKNHAGSPGCEES        346
proCASP2TEV         DNANCPSLQNKPKMFFIQACRGDETDRGVDQQENLYFQGKNHAGSPGCEES        351
proCASP2TEV_C370A   DNANCPSLQNKPKMFFIQACRGDETDRGVDQQENLYFQGKNHAGSPGCEES        351
                    ********************************       *************

proCASP2            D-----AGKEKLPKMRLPTRSDMICGYACLKGTAAMRNTKRGSWYIEALAQVFSERA  398
proCASP2TEV         ENLYFQGGKEKLPKMRLPTRSDMICGYACLKGTAAMRNTKRGSWYIEALAQVFSERA  403
proCASP2TEV_C370A   ENLYFQGGKEKLPKMRLPTRSDMICGYAALKGTAAMRNTKRGSWYIEALAQVFSERA  403
                          .*********************.****************************
```

**Figure S19**. **Sequences of engineered proCASP2TEV cleavable constructs**. Sequence alignments of truncated caspase-2 UniProtKB-1 isoform (truncated residue from D301 and earlier) aligned with the indicated proCASP2TEV cleavable constructs. Highlighted in yellow is the catalytic cysteine residue, C320 and the non-catalytic cysteine residue, C370. Highlighted in green are the protease cleavable residues D333 and D347 that were replaced with a TEV recognition sequence, ENLYFQG.

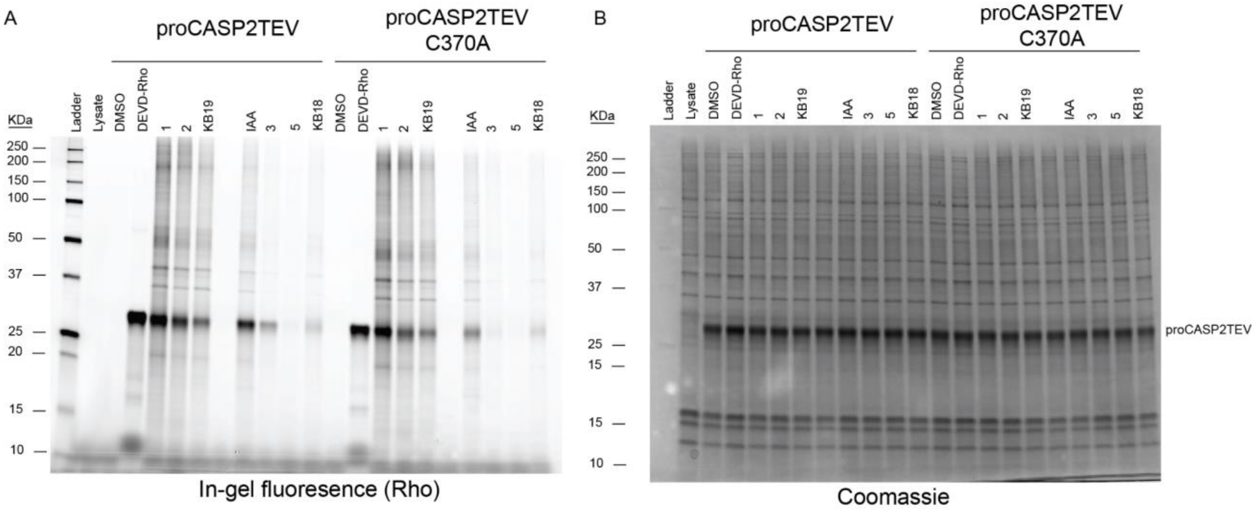

**Figure S20. Click compound 3 does not label proCASP2TEV_C370A**. Labeling of **proCASP2TEV and proCASP2TEV_C370A by alkyne probes.** Gel-based ABPP analysis of the indicated procaspases in whole cell lysates treated for 1h with the indicated compounds (10 µM for all click probes and 2 µM final for Rho-DEVD-AOMK), click conjugation to rhodamine azide and in-gel analysis. (A) in-gel fluorescence and (B) Coomassie InstantBlue visualization of protein loading.

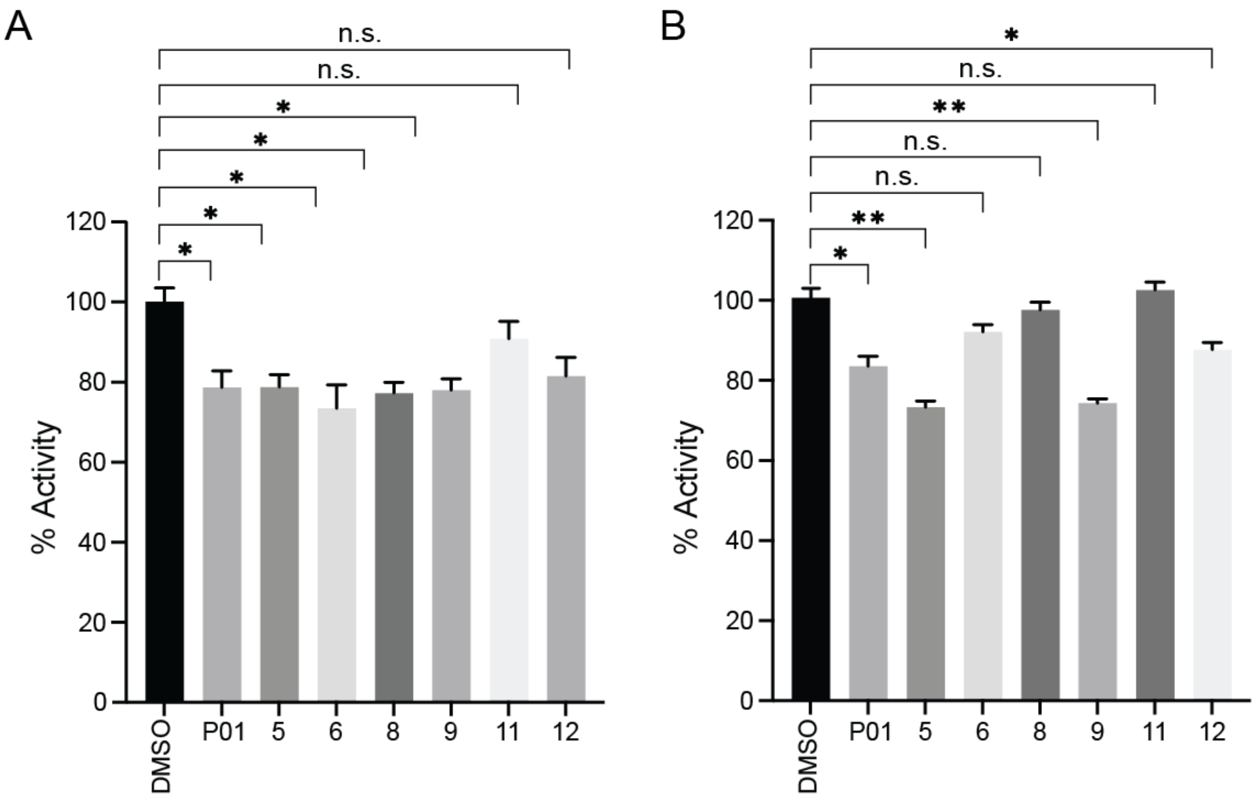

**Figure S21. Activity of proCASP2TEV and proCASP2TEV_C370A with P01 analogues showing little to no inhibition prior to gel filtration.** Recombinant (A) proCASP2TEV and (B) proCASP2TEV_C370A were treated with DMSO vehicle or the indicated compounds at 100 µM final concentration for 1h. Subsequently, samples were then treated with 5 µM Ac-VDVAD-AFC 5 mM DTT, and 333 mM citrate in PBS and analyzed via spectrophotometer monitoring 7-amino-4-trifluoromethylcoumarin (AFC) release by substrate cleavage was detected at $\lambda_{ex}$ = 400 nm and $\lambda_{em}$ = 505 nm every minute after substrate incubation for 1h. Percent activities calculated from the slope of the linear range calculated from the reaction progress curves. Experiments were performed in triplicates with mean relative activity compared to DMSO ± STDEV shown.

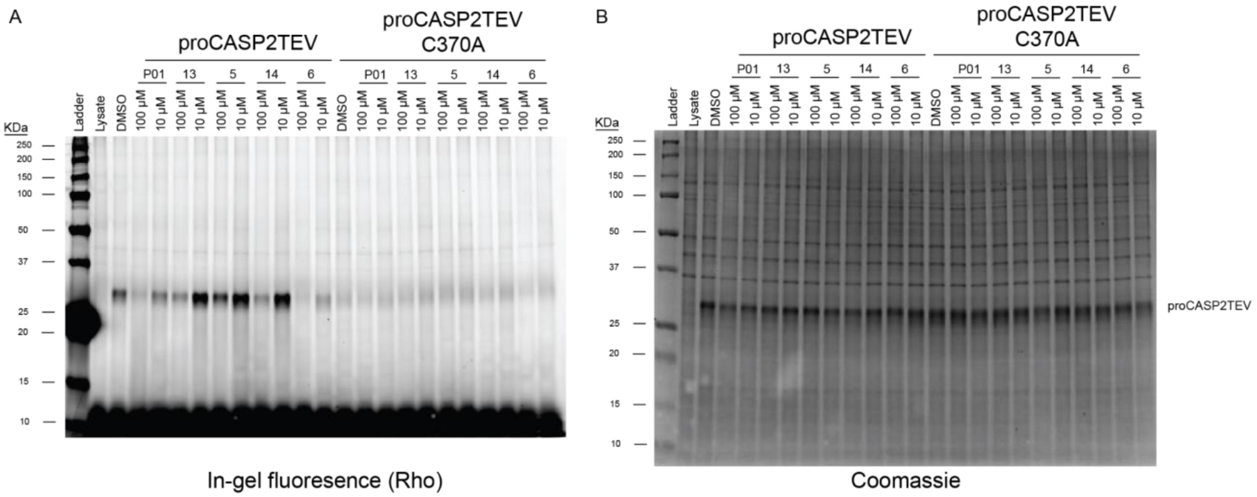

**Figure S22**. **P01 analogues do not compete for labeling of compound 3 similar to proCAS2 ABPP assay.** The indicated procaspases in whole cell lysates were treated for 1h with the indicated compounds at the indicated concentrations, followed by **3** (10 µM, 1h), click conjugation to rhodamine azide and in-gel analysis. (A) in-gel fluorescence and (B) Coomassie InstantBlue visualization of protein loading.

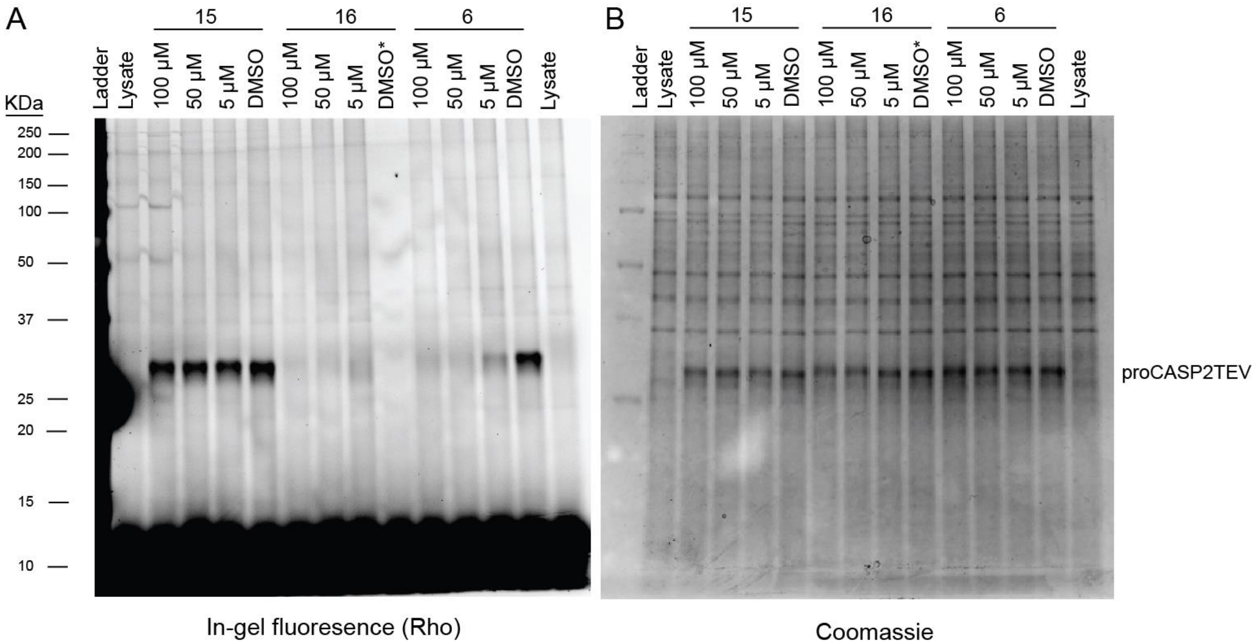

**Figure S23**. **Dase-dependent labeling by electrophilic fragments for proCASP2TEV.** proCASP2TEV in whole cell lysates were treated for 1h with the indicated compounds at the indicated concentrations, followed by **3** (10 µM, 1h), click conjugation to rhodamine azide and in-gel analysis. (A) in-gel fluorescence and (B) Coomassie InstantBlue visualization of protein loading. *DMSO indicates no treatment with compound **3**.

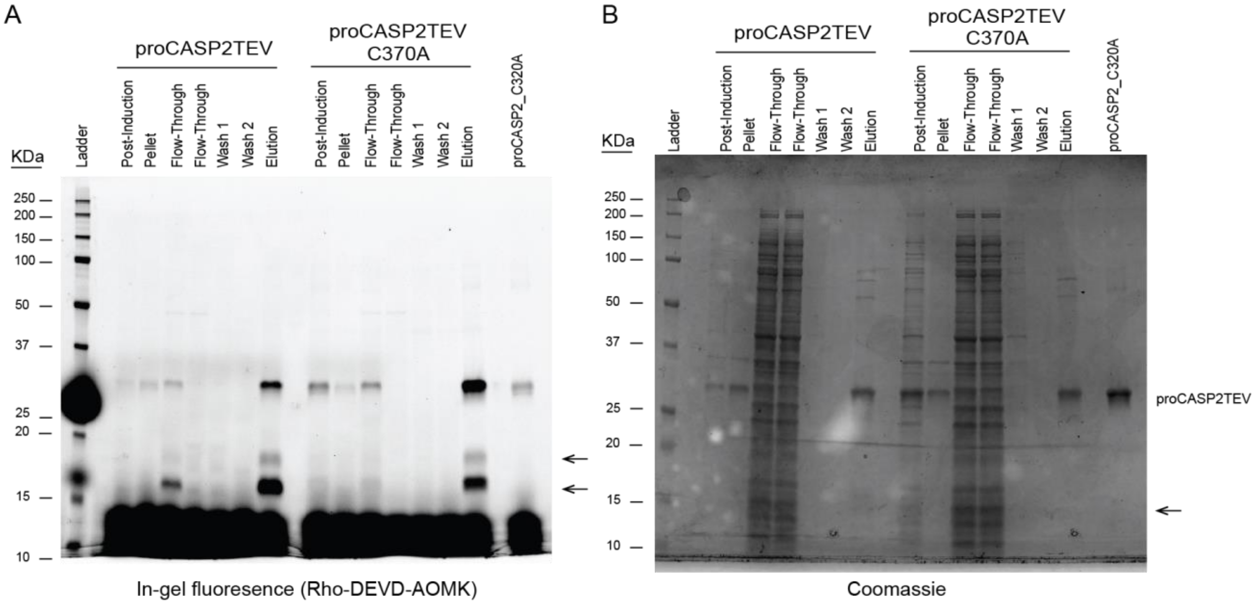

**Figure S24**. **Gel-Based ABPP assessment of recombinant proCASP2TEV and proCASP2TEV_C370A auto-cleaving properties**. Protein purification after IPTG induction and overexpression in BL21 (DE3) *E.coli* cells. Lysate or eluant from each purification step was labeled by **Rho-DEVD-AOMK** (2 µM, 1h) followed by (A) in-gel fluorescence and (B) Coomassie InstantBlue visualization of protein loading. Arrows indicate cleaved caspase-2 as labeled by **Rho-DEVD-AOMK**.

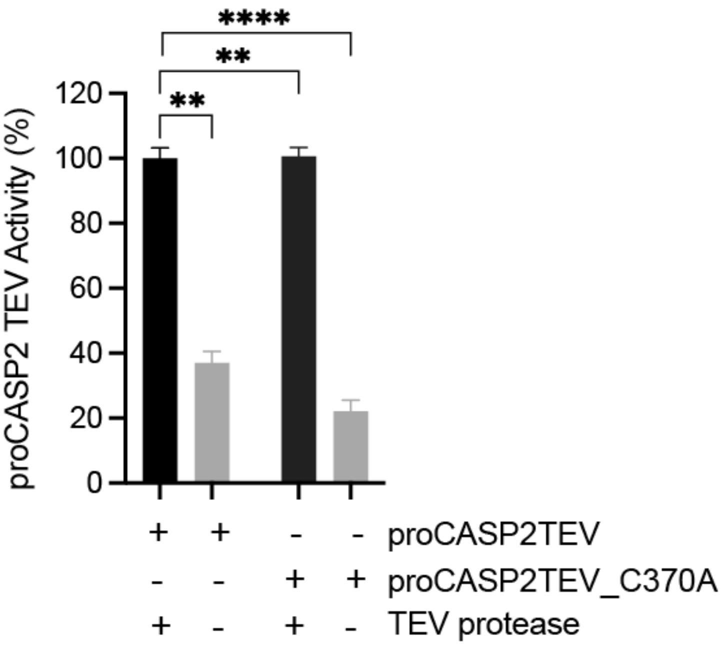

**Figure S25**. **TEV-induced activity of proCASPTEV proteins.** Activity of recombinant protein assessed using fluorogenic substrate Ac-VDVAD-AFC (10 µM substrate) with fluorescence emission spectra ($\lambda_{ex}$ = 400 nm and $\lambda_{em}$ = 505 nm) monitored by multimodal plate reader with percentage activity relative to +TEV protease samples calculated from the linear range of the reaction curves. Data represent mean values ± STDEV for three technical replicates. Statistical significance was calculated with unpaired Student's t-tests, ** p<0.001, ****p<0.0001.

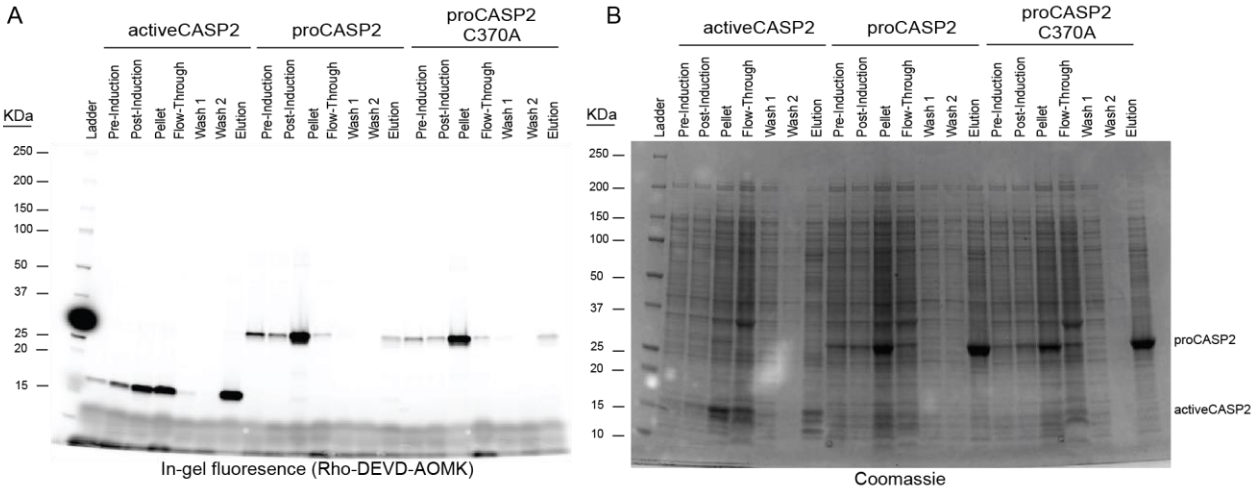

**Figure S26. Gel-Based ABPP assessment of recombinant activeCASP2, proCASP2, and proCASP2_C370A auto-cleaving properties**. Protein purification after IPTG induction and overexpression in BL21 (DE3) *E.coli* cells. Lysate or eluant from each purification step was labeled by **Rho-DEVD-AOMK** (2 µM, 1h) followed by (A) in-gel fluorescence and (B) Coomassie InstantBlue visualization of protein loading.

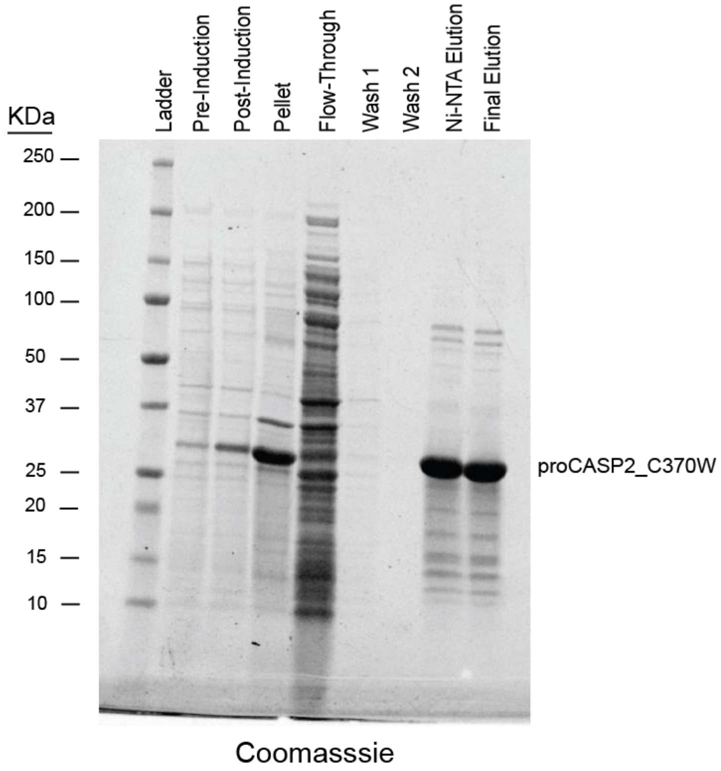

**Figure S27. Gel-Based ABPP assessment of recombinant proCASP2_C370W auto-cleaving properties**. Protein purification after IPTG induction and overexpression in BL21 (DE3) *E.coli* cells. Lysate or eluant from each purification step was visualized by Coomassie InstantBlue.

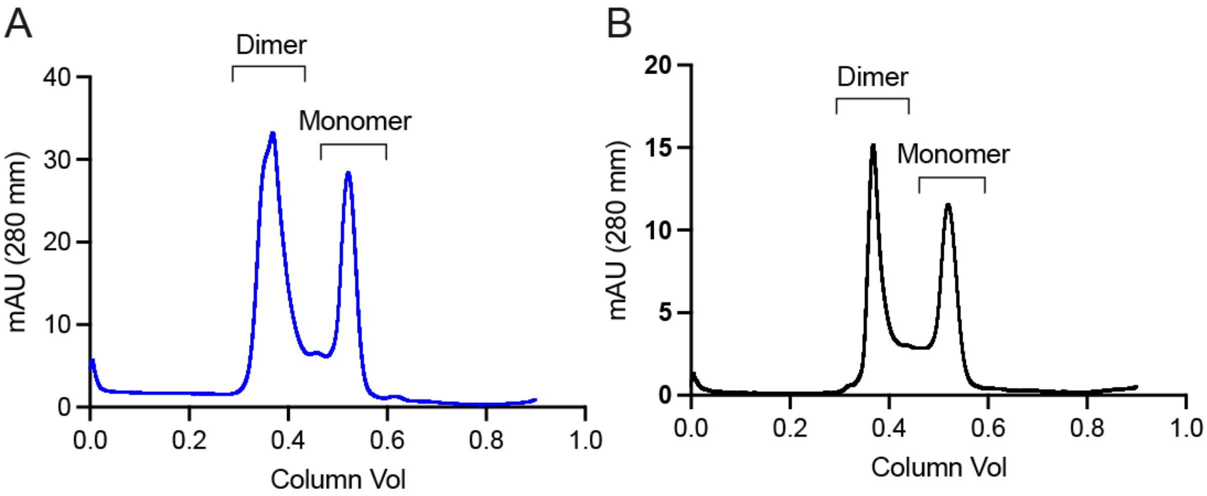

**Figure S28**. **Gel-filtration analysis of recombinant proCASP2 and proCASP2_C370A**. Superdex 75 gel filtration fast protein liquid chromatography (FPLC) elution profiles of recombinant proCASP2 constructs. (A) proCASP2 and (B) proCASP2_C370A, with peaks corresponding to molecular weights consistent with dimeric (column volumes 0.30 - 0.45) and monomeric (column volumes 0.48 - 0.55) proteins.

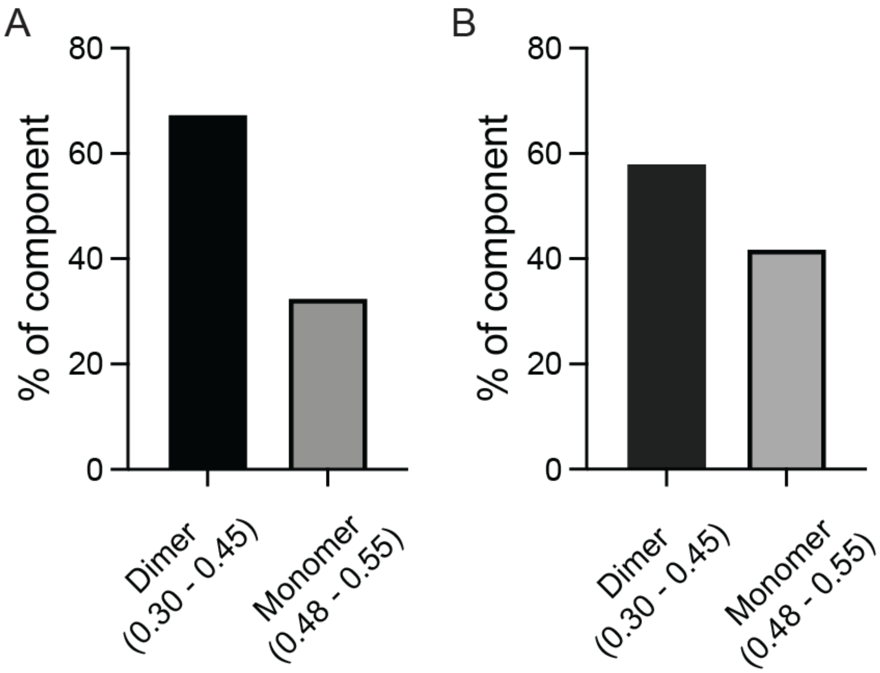

**Figure S29. Quantification of the relative peak areas for Figure S28.** Percent of dimer and monomer (% of component) for (A) proCASP2 and (B) proCASP2_C370A. Percent of component was determined by area-under-the-curve (AUC) calculations (AUC integrated measurement tool from GraphPad Prism 9) of dimeric gel filtration column volume fractions (0.30 - 0.45) and monomeric fractions (0.48 - 0.55) from proCASP2 (0.3 mg/mL).

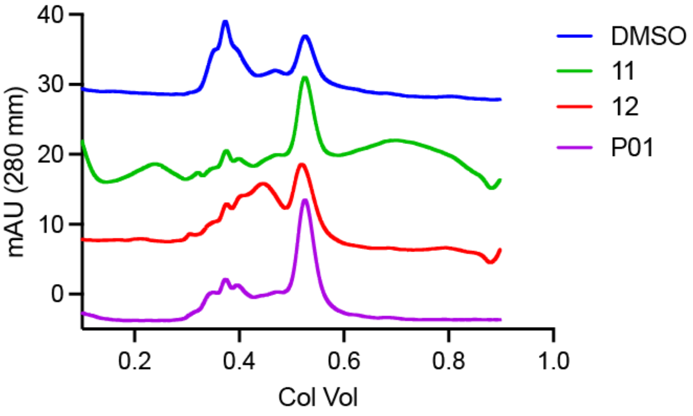

**Figure S30. Assessment of compound-induced changes to proCASP2TEV gel filtration elution profile.** proCASP2TEV was treated for 1h with the indicated compounds (100 μM, 30 °C, 1h) followed by Superdex 75 gel filtration fast protein liquid chromatography (FPLC).

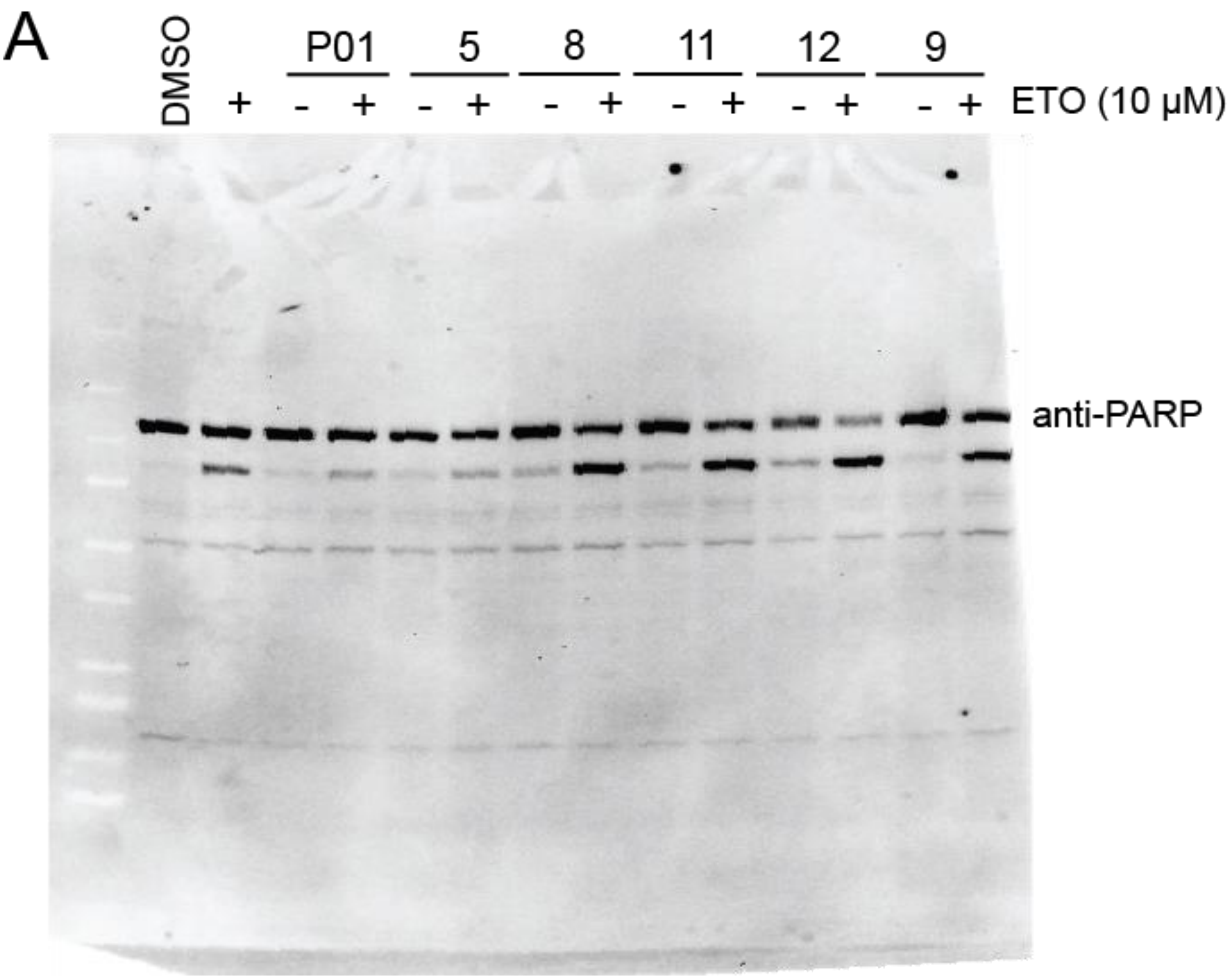

B

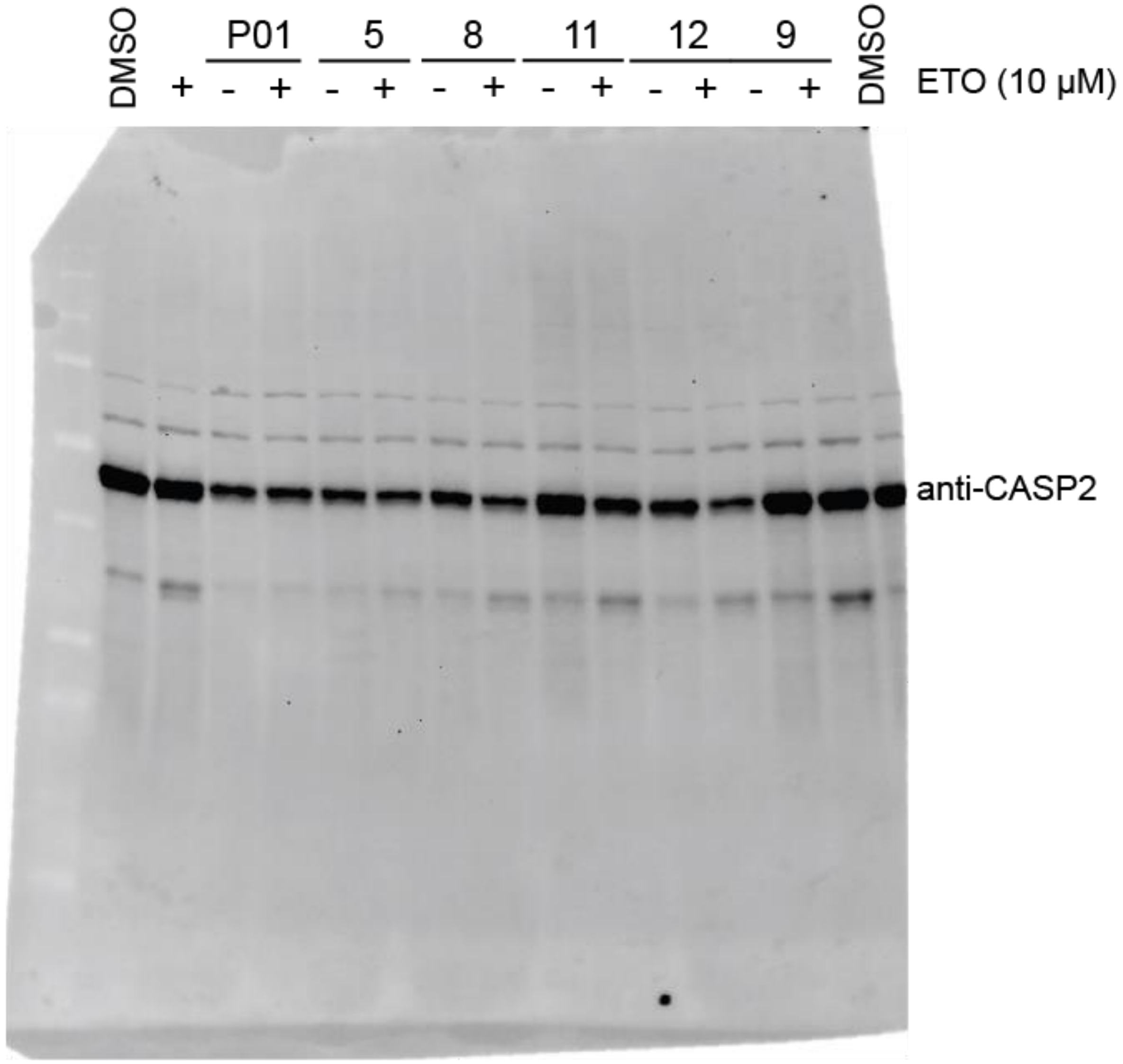

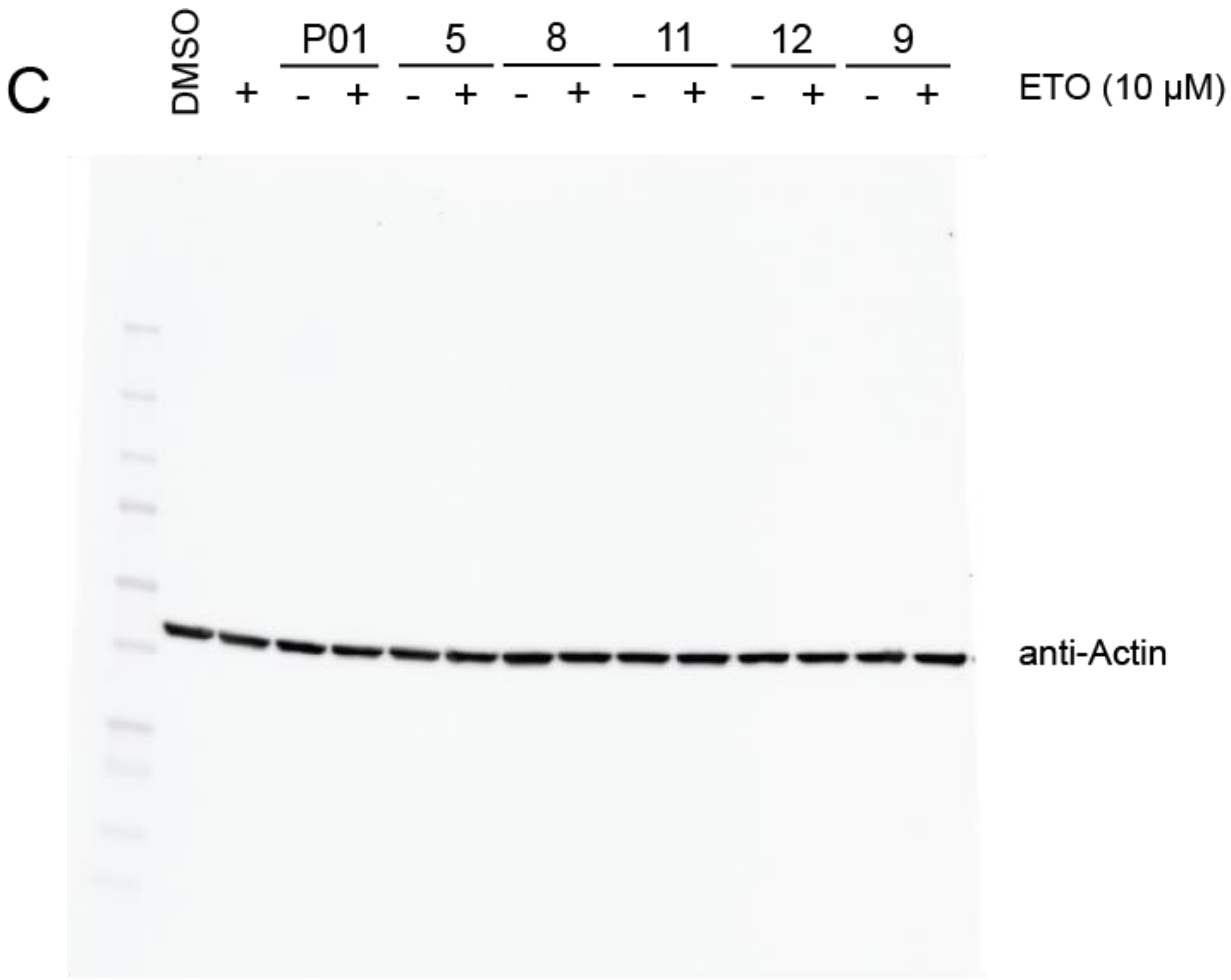

**Figure S31. Full Length immunoblots corresponding to Figure 6D**. Comparison of the effects of the indicated compounds [1h pre-treatment with **P01**, **5**, and **6** (25 μM) and **8**, **9**, **10**, **11**, and **12** (100 μM) on etoposide (10 μM, 10h)-induced apoptosis with cleavage of PARP, and CASP2 visualized by western blotting. (A) anti-PARP (1:1000). (B) anti-CASP2 (1:1000). (C) anti-β-actin (1:5000) dilution.

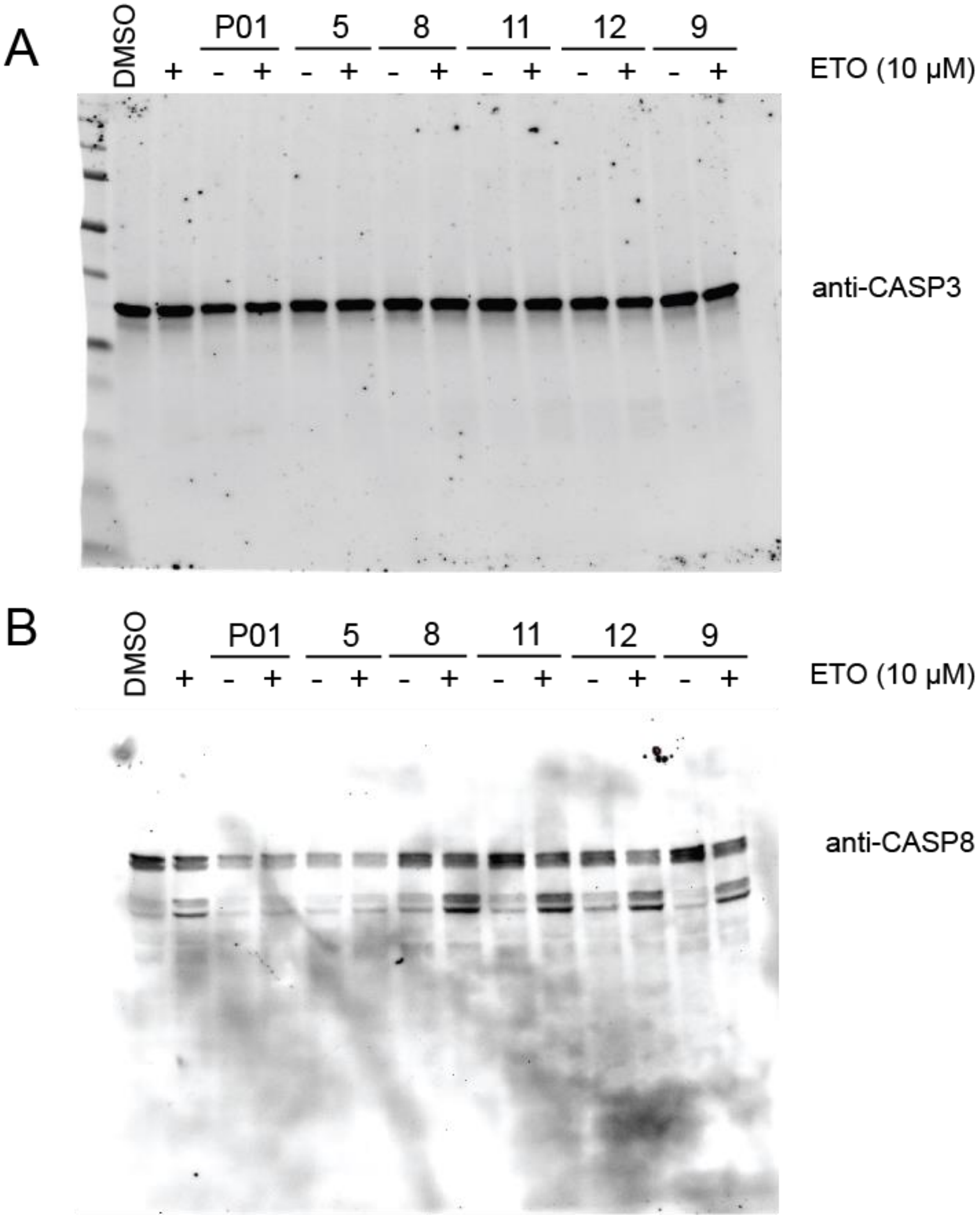
A
DMSO
P01
5
8
11
12
9
+ - + - + - + - + - + - +
ETO (10 µM)
anti-CASP3
B
DMSO
P01
5
8
11
12
9
+ - + - + - + - + - + - +
ETO (10 µM)
anti-CASP8

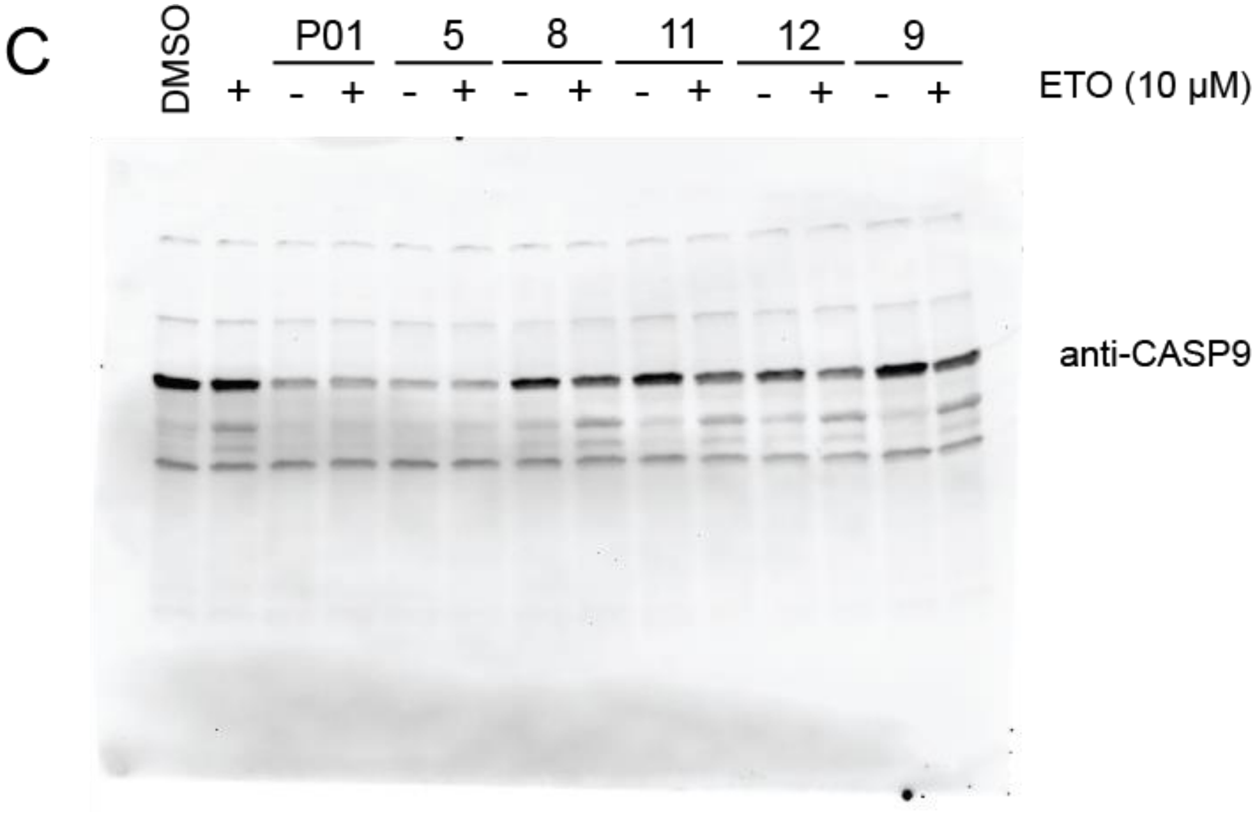
C
DMSO
P01 5 8 11 12 9
+ - + - + - + - + - + - +
ETO (10 μM)
anti-CASP9

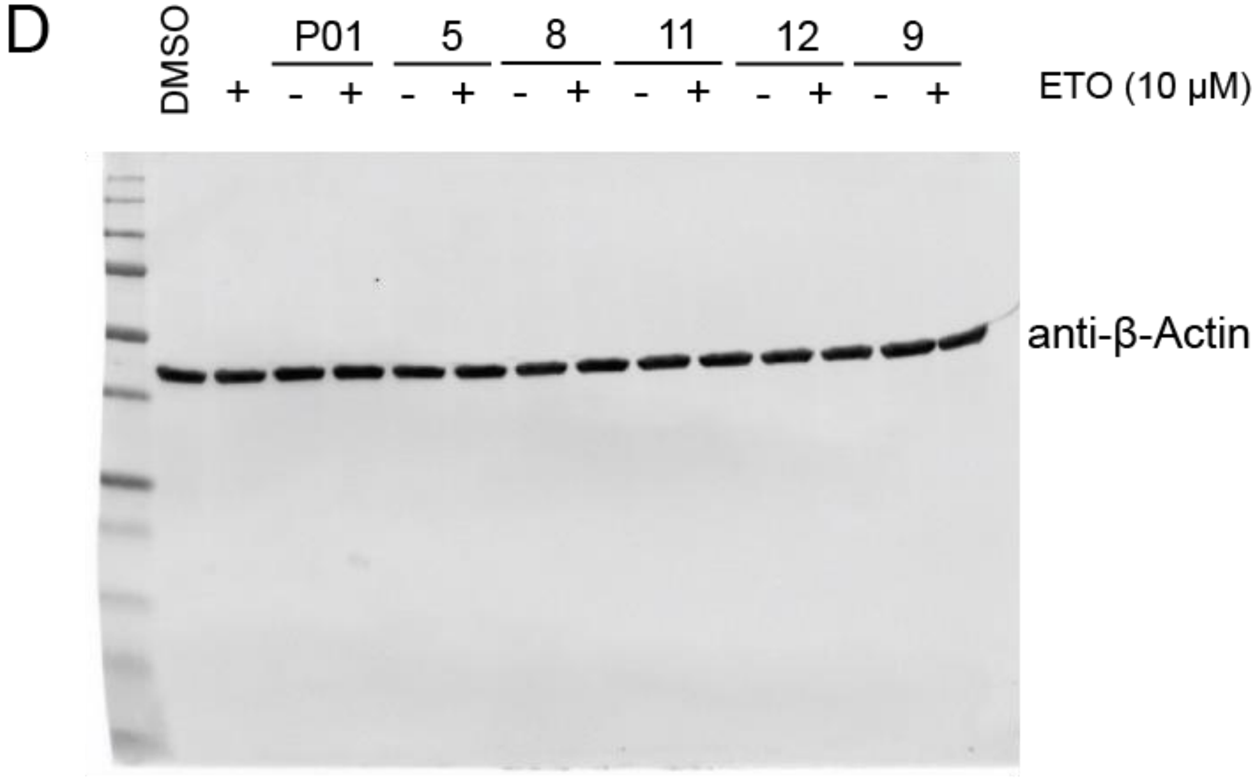
D
DMSO
P01 5 8 11 12 9
+ - + - + - + - + - + - +
ETO (10 μM)
anti-β-Actin

**Figure S32. Full Length immunoblots corresponding to Figure 6D**. Comparison of the effects of the indicated compounds [1h pre-treatment with **P01**, **5**, and **6** (25 µM) and **8**, **9**, **10**, **11**, and **12** (100 µM)] on etoposide (10 µM, 10h)-induced apoptosis monitoring cleavage of CASP3, CASP8, and CASP9. (A) anti-CASP3 (1:1000). (B) anti-CASP8 (1:1000). (C) anti -CASP9 (D) anti-β-actin (1:5000) dilution. Blots represent re-analysis of cell lysates analyzed in **Figure S31**.

### (B) Supplementary Tables

**Table S1.** Summary list of recombinant pET23B caspase-2 plasmids (Residues 170-452) used in this study.

| Construct | Mutation | Source |
|---|---|---|
| activeCASP2 | n/a | Current study |
| activeCASP2_C370A | C370A | Current study |
| activeCASP2_C320A | C320A | Current study |
| activeCSP2C320A_C370A | C320A, C370A | Current study |
| proCASP2_D326A | D326A, D333A, D347A | Current study |
| proCASP2 | D333A, D347A | Current study |
| proCASP2_C370A | C370A, D333A, D347A | Current study |
| proCASP2TEV | D333ENLYFQG, D347ENLYFQG | Current study |
| proCASP2TEV_C370A | D333ENLYFQG, D347ENLYFQG, C370A | Current study |

**Table S2-5.** Datasets corresponding to each figure, provided in the attached supplementary files.

**Table S6.** Summary list of recombinant caspase-8 and caspase-10 as well as purchased caspase-3 constructs used in this study.

| Construct | Mutation | Source |
|---|---|---|
| activeCASP3 | n/a | Vickers, C.J., *et al*. 2013[1] |
| activeCASP8 | n/a | Backus, K.M., *et al*. 2016[10] |

| | | |
|---|---|---|
| activeCASP10 | n/a | Backus, K.M., *et al*. 2016[10] |
| proCASP8 | D347A, D384A, C409S, C433S | Backus, K.M., *et al*. 2016[10] |
| proCASP10 | D415A | Backus, K.M., *et al*. 2016[10] |

**Table S7.** Files in Proteomics Identification Database (PRIDE) datasets. PRIDE IDENTIFIER:PXD042362 and PXD046269.

| Figure | File name | Compound | Experiment |
|---|---|---|---|
| Figure 1B | 2020-10-07-JOC-70min_FAIMS_3cv_-35_-45_-60_OTIT_SP3_B2-4.csv_results.raw | | isoTOP_nonApop |
| | 2020-10-07-JOC-70min_FAIMS_3cv_-35_-45_-60_OTIT_SP3_B3-5.csv_results.raw | | isoTOP_nonApop |
| | 2020-10-07-JOC-70min_FAIMS_3cv_-35_-45_-60_OTIT_SP3_S1-1.csv_results.raw | | isoTOP_nonApop |
| | 2020-10-07-JOC-70min_FAIMS_3cv_-35_-45_-60_OTIT_SP3_S2-2.csv_results.raw | | isoTOP_nonApop |
| | 2020-10-07-JOC-70min_FAIMS_3cv_-35_-45_-60_OTIT_SP3_S3-3.csv_results.raw | | isoTOP_nonApop |
| | 2020-12-05-kb-70min_FAIMS_100micron_3cv_-35_-45_-55--OTIT_JOC_7.csv_results.raw | | isoTOP_nonApop |
| | 2020-12-05-kb-70min_FAIMS_100micron_3cv_-35_-45_-55--OTIT_JOC_8.csv_results.raw | | isoTOP_nonApop |
| Figure 1C | 2020-10-24-JOC-70min_FAIMS_3cv_-35_-45_-60_OTIT_1-2-rerun-1.csv_results.raw | | isoTOP_Apop |
| | 2020-10-24-JOC-70min_FAIMS_3cv_-35_-45_-60_OTIT_1-3-rerun-2.csv_results.raw | | isoTOP_Apop |
| | 2020-10-24-JOC-70min_FAIMS_3cv_-35_-45_-60_OTIT_2-3.csv_results.raw | | isoTOP_Apop |

| | | | |
|---|---|---|---|
| | 2020-10-24-JOC-70min_FAIMS_3cv_-35_-45_-60_OTIT_3-4.csv_results.raw | | isoTOP_Apop |
| | 2020-12-05-kb-70min_FAIMS_100micron_3cv_-35_-45_-55--OTIT_JOC_1.csv_results.raw | | isoTOP_Apop |
| | 2020-12-05-kb-70min_FAIMS_100micron_3cv_-35_-45_-55--OTIT_JOC_2.csv_results.raw | | isoTOP_Apop |
| | 2020-12-6-kb-70min_FAIMS_100micron_3cv_-35_-45_-55--OTIT-JOC_3 | | isoTOP_Apop |
| Figure 3E | 2022-05-04-KB-JOC_isoTOP_EA312_6_quant.raw | 6 | isoTOP_lysates |
| | 2021-08-03-KB-ET-EA312-4-steep.raw | 6 | isoTOP_lysate |
| | 2022-05-15-KB-JOC_isoTOP_EA326_1_quant.raw | 5 | isoTOP_lysates |
| | 2022-05-15-KB-JOC_isoTOP_EA326_2_quant.raw | 5 | isoTOP_lysates |
| | 2022-04-27-KB-JOC_isoTOP_EA329C_10_quant.raw | 15 | isoTOP_lysates |
| | 2022-05-04-KB-JOC_isoTOP_EA329C_9_quant.raw | 15 | isoTOP_lysates |
| | 2022-05-04-KB-JOC_isoTOP_EA329D_7_quant.raw | 16 | isoTOP_lysates |
| | 2022-05-04-KB-JOC_isoTOP_EA329D_8_quant.raw | 16 | isoTOP_lysates |
| | 2021-12-30-KB-JOC-338-A-2_quant.raw | 7 | isoTOP_lysates |
| | 2021-12-30-KB-JOC-338-B-6_quant.raw | 7 | isoTOP_lysates |
| | 2021-12-9-KB-JOC-70min-isoTOP-EA339-8_quant.raw | 8 | isoTOP_lys |

| | | | |
|---|---|---|---|
| | | | ates |
| | 2021-12-9-KB-JOC-70min-isoTOP-EA339-7_quant.raw | 8 | isoTOP_lysates |
| | 2022-09-18-KB-FAIMS-JOC-isoTOP-lysate-SO339-8.pep.raw | 8 | isoTOP_lysates |
| | 2021-12-9-KB-JOC-70min-isoTOP-MPP-4.raw | P01 | isoTOP_lysates |
| | 2021-12-9-KB-JOC-70min-isoTOP-MPP-3.raw | P01 | isoTOP_lysates |
| | 2021-12-9-KB-JOC-70min-isoTOP-SO22-2_quant.raw | 10 | isoTOP_lysates |
| | 2022-05-04-KB-JOC_isoTOP_SO22_5_quant.raw | 10 | isoTOP_lysates |
| | 2022-09-18-KB-FAIMS-JOC-isoTOP-lysate-SO86-1.raw | 12 | isoTOP_lysates |
| | 2022-09-18-KB-FAIMS-JOC-isoTOP-lysate-SO86-2.raw | 12 | isoTOP_lysates |
| | 2022-09-18-KB-FAIMS-JOC-isoTOP-lysate-SO103-4.raw | 11 | isoTOP_lysates |
| | 2022-09-23-KB-FAIMS-JOC-isoTOP-lysate-SO103-3_rerun091822.raw | 11 | isoTOP_lysates |
| | 2022-09-23-KB-FAIMS-JOC-isoTOP-lysate-SO139-5_rerun091822.raw | 9 | isoTOP_lysates |
| | 2022-09-23-KB-FAIMS-JOC-isoTOP-lysate-SO139-6_rerun091822.raw | 9 | isoTOP_lysates |
| Figure S7 | 2021-09-12-KB-JOC-DMSO-1.raw | | Recombinant protein |
| | 2021-09-1-KB-JOC-DMSO-2.raw | | Recombinant protein |

| | | | |
|---|---|---|---|
| | 2021-09-1-KB-JOC-DMSO-3.raw | | Recombinant protein |
| | 2021-09-1-KB-JOC-AZ12-1.raw | 3 | Recombinant protein |
| | 2021-09-1-KB-JOC-AZ12-2.raw | 3 | Recombinant protein |
| | 2021-09-1-KB-JOC-AZ12-3.raw | 3 | Recombinant protein |
| Figure S18 | 2022-06-18-KB-noFAIMS-JOC--IsoTOP-SO139_RT_5.raw | 9 | isoTOP |
| | 2022-06-18-KB-noFAIMS-JOC--IsoTOP-SO139_RT_6.raw | 9 | isoTOP |
| | 2022-09-23-KB-FAIMS-JOC-isoTOP-lysate-SO139-5_rerun091822.raw | 9 | isoTOP |
| | 2022-09-23-KB-FAIMS-JOC-isoTOP-lysate-SO139-6_rerun091822.raw | 9 | isoTOP |
| Figure 6A | 2022-09-10-KB-FAIMS-JOC-isoTOP-incell-MPP_frozen_2.raw | P01 | isoTOP_in-cell |
| | 2022-09-09-KB-FAIMS-JOC-isoTOP-incell-MPP_frozencells_3_rerun0828.raw | P01 | isoTOP_in-cell |
| | 2022-09-09-KB-FAIMS-JOC-isoTOP-incell-MPP_freshcells_4_rerun0828.raw | P01 | isoTOP_in-cell |
| | 2022-12-07-KB-JOC-isoTOP_incell_SO22_9.raw | 10 | isoTOP_in-cell |
| | 2022-12-07-KB-JOC-isoTOP_incell_SO22_10.raw | 10 | isoTOP_in-cell |
| | 2022-12-06-KB-JOC-isoTOP_incell_SO86_1.raw | 12 | isoTOP_in-cell |
| | 2022-12-06-KB-JOC-isoTOP_incell_SO86_2.raw | 12 | isoTOP_in-cell |

| | | | |
|---|---|---|---|
| | 2022-12-06-KB-JOC-isoTOP_incell_SO103_3.raw | 11 | isoTOP_in-cell |
| | 2022-12-06-KB-JOC-isoTOP_incell_SO103_4.raw | 11 | isoTOP_in-cell |
| | 2022-12-06-KB-JOC-isoTOP_incell_SO139_5.raw | 9 | isoTOP_in-cell |
| | 2022-12-08-KB-JOC-isoTOP_incell_SO139_6_rerun.raw | 9 | isoTOP_in-cell |
| | 2023_02_14_KB_FAIMS_incell_isoTOP_EA339_JOC_1.raw | 8 | isoTOP_in-cell |
| | 2023_02_14_KB_FAIMS_incell_isoTOP_EA339_JOC_2.raw | 8 | isoTOP_in-cell |
| | 2023_02_14_KB_FAIMS_incell_isoTOP_EA339_JOC_3.raw | 8 | isoTOP_in-cell |
| | 2023_02_14_KB_FAIMS_incell_isoTOP_EA312_JOC_4.raw | 6 | isoTOP_in-cell |
| | 2023_02_14_KB_FAIMS_incell_isoTOP_EA312_JOC_5.raw | 6 | isoTOP_in-cell |
| | 2023_02_14_KB_FAIMS_incell_isoTOP_EA312_JOC_6.raw | 6 | isoTOP_in-cell |
| | 2023_02_14_KB_FAIMS_incell_isoTOP_EA326_JOC_7.raw | 5 | isoTOP_in-cell |
| | 2023_02_14_KB_FAIMS_incell_isoTOP_EA326_JOC_8.raw | 5 | isoTOP_in-cell |
| | 2023_02_14_KB_FAIMS_incell_isoTOP_EA326_JOC_9.raw | 5 | isoTOP_in-cell |

**Table S8.** List of caspase-2 primer sequences used in this study.

| Purpose | Primer Sequence |
|---|---|

| | | |
|---|---|---|
| Subclone Caspase-2 into pET23b (Residues 170-452) | 5'-AA**<u>CATATG</u>**GGTCCTGTCTGCCTTCAGGTG-3' | Forward |
| Subclone Caspase-2 into pET23b | 5'-AA**<u>CTCGAG</u>**TGTGGGAGGGTGTCC-3' | Reverse |
| Mutation C370A | 5'-GCGGCTATGCCGCCCTCAAAGGGACTGCCGCCATGCGG-3' | Forward |
| | 3'-GTCCCTTTGAGGGCGGCATAGCCGCATATCATGTCTG-5' | Reverse |
| Mutation C320A | 5'-CTTCATCCAGGCCGCCCGTGGAGATGAGACTGATCG-3' | Forward |
| | 3'-CATCTCCACGGGCGGCCTGGATGAAGAACATTTTTG-5' | Reverse |
| Mutation D333A | 5’-GAGATGAGACTGCTCGTGGGGTTGACCAACAAGCTGGAAAGAACCACG-3’ | Forward |
| | 5’-GTGGAGATGAGACTGCTCGTGGGGTTGACCAACAAGCTGGAAAGAACCACGCAGG-3’ | Reverse |
| Mutation D347A | 5’-CCCCTGGGTGCGAGGAGAGTGCTGCCGGTAAAGAAAAGTTG-3’ | Forward |
| | 5’CAACTTTTCTTTACCGGCAGCACTCTCCTCGCACCCAGGGG-3’ | Reverse |
| Mutation C370W | 5'-GCGGCTATGCCTGGCTCAAAGGGACTGCCGCCATGCGG-3' | Forward |
| | 3'-CAGTCCCTTTGAGCCAGGCATAGCCGCATATCATGTCTG-5' | Reverse |
| | 5'-GAAAATCTCTACTTCCAGGGCGGTAAAGAAAAGTTGCCG-3' | Forward |

| | | |
|---|---|---|
| ENLYFQG insertion | 3'-CTTTACCGCCCTGGAAGTAGAGATTTTCACTCTCCTCGC-5' | Reverse |

**(C) Biology Methods**

*Cell lines, culture conditions*
Jurkat cells were cultured in Roswell Park Memorial Institute (RPMI) media (Fisher Scientific, 11875119) with 10% fetal bovine serum (Avantor Seradigm Lot # 214B17) and 100U/mL penicillin and 100U/mL streptomycin at 37℃ and 5% $CO_2$. Jurkat cells were obtained from ATCC (TIB-152). Cell culture reagents including RPMI 1640 media, trypsin-EDTA and penicillin/streptomycin (Pen/Strep) were purchased from Fisher Scientific.

*Mycoplasma testing*
Mycoplasma testing was conducted monthly using the MycoAlert® kit (LT07-703, Lonza Rockland, Rockland, ME) following the manufacturer's instructions.

*Cell Harvesting*
Suspension cells were centrifuged at 1,400 *g* for 5 minutes and the supernatant was aspirated. The pellets were then washed in 10 mL PBS and centrifuged at 1,400 *g* for 5 minutes. PBS wash was repeated, and the subsequent pellet was then resuspended in 1 mL PBS in a microcentrifuge tube and centrifuged at 1,400 *g* for 5 minutes. The supernatant was aspirated, and the cells were stored at -80 ℃.

*Cell Lysis*
Jurkat lysates used for competitive ABPP gel analysis were lysed using an Ultrasonic Probe Sonicator at Power 2 for 10 pulses, 1 second pulse, 1 second off on ice. For western blotting samples, cells were reconstituted and lysed using cold 100 uL of 0.3% 3-[(3-Cholamidopropyl) dimethylammonio]-1-propanesulfonate (CHAPS) buffer in PBS. The reconstituted cell pellet was left to incubate with the CHAPS buffer on ice for 15 min. The samples were then harvested by centrifugation (1,400 x g, 10 min) and the clarified supernatant was then transferred to a new tube.

*Plasmids*
Caspase-2 (Residues 170-452) was subcloned into pET23b (+) and the point mutations and insertions generated by site-directed mutagenesis using the primer sequences listed in Table S8. Plasmids were propagated in chemically competent TOP10 cells.

*Protein Concentration*
Protein concentrations were determined using a Bio-Rad DC protein assay kit following the manufacturer's instructions using reagents from Bio-Rad Life Science (DC Reagent A and B, 5000113 and 5000114).

*isoTOP-ABPP proteomic sample preparation*

**Cell processing:** Apoptotic Jurkat cells (treated with 50 ng/mL FasL for 4h) or non-apoptotic Jurkat cells (treated with DMSO for 4 h) were harvested at 1.0 x$10^6$ cells/mL and harvested by centrifugation (1,400 x *g*). Collected cells were then resuspended in 10 mL of cold PBS and subsequently harvested by centrifugation (1,400 x *g*), and this wash step was repeated 2 cycles. Cell pellets were then reconstituted in 500 uL of cold PBS and subjected to Ultrasonic Probe Sonicator lysis (Power 2, 10x pulses; 1 second pulse, 1 second off) on ice. Lysate concentrations were then adjusted to 2 mg/mL.

**Electrophilic fragment and IA-alkyne labeling:** For cysteine reactivity analysis, 200 µL lysates (2 mg/mL) were then labeled with either 10 µM or 100 µM iodoacetamide alkyne (**IAA**) for 1h at ambient temperature. For lysate-based compound screening, samples were treated with either DMSO or fragment electrophiles at 25 µM (**P01**, **5**, **6**) or 100 µM (**3**, **7-16**) for 1h followed by subsequent labeling with 200 µM **IAA** for 1h. For the in-cell treated samples, 10 ml of 1.2 x $10^6$ cells/mL were treated with DMSO or electrophilic fragments (25 or 100 µM) for 1h. Cells were subsequently harvested as detailed for *Cell Harvesting* and lysed as detailed for *Cell Lysis*. Samples were prepared in at least biological duplicate.

**Click chemistry:** Samples were subjected to bioorthogonal copper(I)-catalyzed azide-alkyne cycloaddition (CuAAC)[11] or "click" conjugation to previously reported isotopically labeled *tobacco etch virus* (TEV)-cleavable biotinylated peptide tags[2]. To 200 µL cell lysates (2 mg/mL), samples were combined with a premixed cocktail of click reagents consisting of TEV tags (4µl of 5 mM stock, final concentration= 100 µM), TCEP (4 µl of fresh 50 mM stock in water, final concentration 1 mM), TBTA (12 µl of 1.7 mM stock in DMSO/t-butanol 1:4, final concentration = 100 µM), and CuSO4 (4 µl of 50 mM stock in water, final concentration = 1 mM). After 1h, the samples were then combined pairwise (400 µL total) and treated with 40 µL of 10% SDS (1% SDS final) followed by 0.5 µL of benzonase (Fisher Scientific, 707464). Samples were left to incubate for 30 min at 37 ℃. Following benzonase treatment, samples were subjected to Single-Pot Solid-Phase-enhanced sample-preparation (SP3).

**Single-Pot Solid-Phase-enhanced sample-preparation (SP3):** Following the previously reported protocol[9], for each 400 µL sample (400 µg protein), 40 µL Sera-Mag SpeedBeadsTM Carboxyl Magnetic Beads, hydrophobic (Thermo Scientific™, 09-981-123) and 40 µL Sera-Mag SpeedBeadsTM Carboxyl Magnetic Beads, hydrophilic (Thermo Scientific™, 09-981-121) were gently mixed and washed with 1 mL distilled water. Beads were combined using a magnetic rack (Sergi Lab Supplies, 1005a) and water was carefully aspirated. Washes were repeated for a total of 3 times. 80 µL of mixed beads were then added to the 400 µL of combined samples. The bead-sample mixture was then incubated for 5 min at ambient conditions with shaking (1000 rpm). 200 proof ethanol was added to each sample such that each sample contained ≥55% ethanol by volume (for 480 µL of combined sample/SP3 beads, 600 µL of ethanol was added). The samples were incubated for 5 min at ambient conditions with shaking (1000 rpm). The beads were then washed three times with 600 µL 80 % ethanol in water. After washes, ethanol was removed using the magnetic rack, and beads were then resuspended in 200 μL 0.5 % SDS in PBS containing 2 M urea. 10 μL of 200 mM DTT (10 μM final concentration) was then added to each sample and the samples incubated at 65 ° C for 15 min. Following reduction, 10 μL of 400 mM iodoacetamide (20 μM final concentration) was added to each sample and the samples incubated for 30 min at 37 ° C with shaking at 300 rpm. Subsequently, 600 μL of 200 proof ethanol was added to each sample, and the samples were incubated for 5 min at ambient conditions with shaking (500 rpm). Beads were then washed three times with 80 % ethanol in water. Samples were then diluted with 150 μL 2 M urea in PBS followed by the addition of reconstituted MS grade trypsin (2 µg, Promega, V5111). The samples were subjected to water bath sonication for 1 min and subsequently left to digest overnight (16 - 18hr) at 37℃ and shaking at 200 rpm. The digested peptide solution and SP3 beads were then transferred into 15 mL falcon tubes. Peptides were then rebound to SP3 beads via the addition of 3.8 mL of 100% acetonitrile for a final percentage of ≥95% acetonitrile by volume and the peptides were subjected to shaking at 1000 rpm for 10 minutes at ambient temperature. Beads were collected using a magnet and solution was discarded. Samples were washed with 1 mL of 100% acetonitrile three times. Digested peptides were then eluted with 100 µL of 2% DMSO in water, shaking at 1000 rpm for 30 min at 37℃. Supernatant was collected in a 1.5 mL centrifuge tube on ice after separating SP3 beads with a magnetic rack. SP3 beads were resuspended with an additional 100 µL of 2% DMSO in water, shaking at 1000 rpm for 45 min at 37℃. Supernatant was collected after separating SP3 beads with a magnetic rack and combined with the previous elution volume (200 µL total).

**Streptavidin Enrichment:** Pierce™ Streptavidin Agarose resin (Thermo Scientific™, PI20353) (100 µL of resin) was first washed 3x in 10 mL of PBS by centrifugation at 1,800 x *g* for 3 min per wash. Solution was aspirated, making sure not to disturb spun down resin. After washing, resin was resuspended in 1 mL PBS/sample and re-distributed into 1.5 mL microcentrifuge tubes. The 200 µL peptide elution from previously prepared SP3 method was then added to the 1 mL of PBS/streptavidin resin. Samples were enriched by rotation for 2h at ambient conditions. After enrichment, the resin was collected by centrifugation at 1,400 x g for 5 min, and supernatant was aspirated and discarded. The resin was then subjected to washes 2x in 1 mL of PBS and 2x in 1 mL of water by centrifugation at 1,400 x g for 5 min per wash. After carefully aspirating and discarding the water, the resin/peptide slurry was then treated with a TEV protease following the "Tev digestion" protocol.

**Tev Digest: Following streptavidin enrichment,** samples were resuspended in 75 µL TEV buffer (50 mM Tris, pH 8, 0.5 mM EDTA, 1 mM DTT). To the resuspended samples, 1.5 µL TEV protease (2 mg/mL or 70 µM; MacroLab, UC Berkeley) was added and the reactions were rotated for 7h at 30°C. The samples were then harvested by centrifugation (3,000 x *g*, 1 min) and the supernatant was collected. Samples were then subjected to a final cleanup following the "Sample cleanup" protocol.

**Sample Cleanup:** The collected peptides were then desalted using Pierce™ C18 100 µL Tips (Thermo Scientific™, 87784) following the manufacturer's protocol. Briefly, 10 mL of the following four solutions were prepared; A) 100% acetonitrile, B) 50:50 acetonitrile:ultra pure water, C) Ultra pure water with 0.1% trifluoroacetic acid, and D) 60% acetonitrile with 0.1% trifluoroacetic acid in ultrapure water. Each C18 100 µL tip was first equilibrated with 100 µL with solution A for a total of two times followed by equilibration with solution B for a total of two times. The tips were then washed with 100 µL of solution C for a total of three times. Finally, 100 µL of samples were loaded into the tips and subsequently washed 2x with solution C. Samples were then eluted with 100 µL of solution D. Following desalting, each 100 µL sample was dried by speedvac and reconstituted in 20 µL of 5% acetonitrile and 1% formic acid in water.

*Recombinant proCASP2 proteomic sample preparation*

25 µL recombinant purified proCASP2 (0.5 mg/mL) was first treated with 10 mM DTT followed by buffer exchange using Zeba desalting columns. Samples were treated with DMSO or compound **3** (100 µM) for 1h at 30 ℃. Samples were then subjected to *SP3 clean-up* using 5 µL of mixed SP3 beads. 30 uL beads/sample were then washed with 200 proof ethanol, followed by 2x 80% ethanol in water, and reduced with 10 µL of 200 mM DTT (10 mM final concentration) in samples reconstituted with 200 µL of 2M urea in 0.5% SDS in PBS. Samples were then incubated with 10 µL of 400 mM iodoacetamide (20 mM final concentration) and subjected to on-bead Trypsin digest following the SP3 protocol. Reconstituted samples in 100 µL of 2% DMSO in water were analyzed by LC-MS/MS.

*Proteomics acquisition*

The samples were analyzed by liquid chromatography tandem mass spectrometry (LC-MS/MS) using a Thermo Scientific™ Orbitrap Eclipse™ Tribrid™ mass spectrometer (Thermo Scientific™) coupled to an Easy-nLC™ 1200 pump and to a High Field Asymmetric Waveform Ion Mobility Spectrometry (FAIMS) Interface. Peptides were fractionated online using a 16 cm long, 100 µM inner diameter (ID) fused silica capillary packed in-house with bulk C18 reversed phase resin (particle size, 1.9 µm; pore size, 100 Å; Dr. Maisch GmbH). The 70-minute water-acetonitrile gradient was delivered using a EASY-nLC™ 1200 system at different flow rates (Buffer A: water with 3% DMSO and 0.1% formic acid and Buffer B: 80% acetonitrile with 3% DMSO and 0.1% formic acid). The detailed 70 minute gradient includes 0 – 5 min from 3 % to 10 % at 300 nL/min, 5 – 15 min from 10 % to 20 % at 220 nL/min, 15 – 64 min at 20 to 47% at 220nL/min, 64-66 min from 47 % to 95 % at 250 nL/min, and 66 to 70 min at 95% buffer B in buffer A. Data was collected with charge exclusion (1, 8,>8). Data was acquired using a Data-Dependent Acquisition (DDA) method consisting of a full orbitrap MS1 scan (Resolution = 120,000) followed by sequential MS2 scans (Resolution = 15,000) to utilize the remainder of the 1 second cycle time. Precursor isolation window was set to 1 m/z and high energy c-trap dissociation (HCD) normalized collision energy was set to 30%. Run time 70 minutes Injection volume 5 uL. For data acquired using a FAIMS device 3 compensation voltages (CV; -35, -45, -55V) were used as determined in our previous study[9].

*Data analysis*

For isoTOP ABPP experiments, the Proteomic workflow of FragPipe and its collection of tools were set as default. FragPipe output data was then compiled using our in-house python script. Custom python scripts were implemented to compile modified_peptide_label_quant.tsv (quantification) outputs from FragPipe (v19.0)[12–14]. These scripts were used to calculate an average of median logged ratios for each peptide, cysteine and protein per replicate and experiment condition. As a preprocessing step, logged ratios from singleton peptides were removed before further analysis, and unpaired heavy or light-identified peptides were kept by setting ratios to $\log_2(20)$ or $\log_2(1/20)$. Median heavy over light ratios for the same cysteine residue from cysteine peptides of different charges and miss cleavages in the same replicate were computed. Then, for each experimental condition (apoptotic vs nonapoptotic, DMSO vs treatment, etc.), an average of medians was calculated to obtain a final "average_of_medians" metric. This "average_of_medians" value was utilized to compare the isotopical quantification of the same peptide/cysteine/protein across various experimental conditions or across replicates within the same experiment.

**Protein, peptide, and cysteine identification.** RAW files were searched with MSFragger (v3.7) and FragPipe (v19.0). The proteomic workflow and its collection of tools was set as default and PTMprophet was enabled[15] . Precursor and fragment mass tolerance was set as 20 ppm. Missed cleavages were allowed up to 1. Peptide length was set 7 - 50 and peptide mass range was set 500 - 5000. For identification, cysteine residues were searched with differential modification C+. For ligandability quantification, MS1 labeling quant was enabled with Light set as C+521.30745 and Heavy set as C+527.3213. MS1 intensity ratio of heavy and light labeled cysteine peptides were reported with Ionquant (v1.8.10). Calibrated and deisotoped spectrum files produced by FragPipe were retained and reused for this analysis. The MS search data have been deposited to the ProteomeXchange Consortium via the PRIDE partner repository with the dataset identifiers PXD042403 and PXD046269. File details can be found in **Table S7**. Custom python scripts were implemented to compile labeled peptide datasets. Unique proteins, unique cysteines, and unique peptides were quantified for each dataset. Unique proteins were established based on UniProt protein IDs. Unique peptides were found based on sequences containing a modified cysteine residue. Unique cysteines were classified by an identifier consisting of a UniProt protein ID and the amino acid number of the modified cysteine (ProteinID_C#); residue numbers were found by aligning the peptide sequence to the corresponding UniProt protein sequences found in protein.fas FragPipe output files. For cases where multiple cysteines were labeled on the same peptide, a new identifier for each modified cysteine residue number was created. Multiplexed peptide identifiers were reported as ProteinID_C#1 and ProteinID_C#2, instead of ProteinID_C#1_C#2.

*Protein expression and purification*

The sequence encoding caspase-2, lacking the prodomain (residue numbers 32-121) was subcloned into the pET23b(+) with a c-terminal hexa his-tag. Point mutations (C320A, C370A, D333A, D347A) were created by PCR-based site-directed mutagenesis. Plasmids were propagated in TOP10 chemically competent cells. Single colonies from TOP10 grown cells were collected in 5 mL of LB supplemented with 100 µg/ml ampicillin and grown overnight (16h). Cells were harvested the following day and subjected to Zyppy Plasmid Miniprep following the manufacturer's protocol (Zymo Research, D4037). Following sequencing of plasmids, validated plasmids were transformed to BL21(DE3) *e.coli* cells. Single colonies were picked from an LB agar plate and grown in 10 mL of LB media supplemented with 100 µg/mL ampicillin. The cell culture was then transferred and grown in 1 L of Miller LB medium at 37 °C to an optical density (OD600) of 0.5. The culture was then cooled to 18 °C, induced with 1 mM isopropyl-β-Dgalactopyranoside (IPTG), and incubated for an additional 4h at 18 °C. The cells were centrifuged at 8,000 x *rpm* for 45 min minutes and the cell pellet was measured. The cells were resuspended in 10 mL per 1 g of cells in lysis buffer (100 mM Tris pH 7.5, 100 mM NaCl, 25 mM Imidazole). The resuspended cells were passed through a microfluidizer (Avestin Emulsiflex C3 Homogenizer; 8,000 psi x2 rounds) to ensure lysis. The cell debris was removed by centrifugation (20,000 x *g*, 45 min*)* and the supernatant was resuspended with 1 mL of Hispur Ni-NTA agarose resin (Thermo Scientific™, PI88222). The sample was washed with two rounds of lysis buffer (2 x 50 mL). His-tagged caspase-2 was eluted from the resin using an elution buffer with high imidazole concentration (100 mM Tris pH 7.5, 100 mM NaCl, 250 mM Imidazole). The eluted sample was concentrated (Amicon Ultra Centrifugal Filter Unit, 4 mL 10 kDa, Fisher Scientific, UFC801024) and buffer exchanged via PD10 desalting column (Cytiva, GE17-0851-01) into storage buffer (20 mM Tris pH 7.5, 50 mM NaCl, 5 mM DTT). Recombinant TEV protease was purchased from the Berkeley MacroLab QB3, where it was was expressed as a double mutant (L56V / S135G) pRK793 plasmid in Rosetta2(DE3)pLysS cells (TEV-DM-Prk793 L56V/ S135G) and stored in 25 mM HEPES pH 7.5, 400 mM NaCl, 10% glycerol, 1 mM DTT. Recombinant proCASP8 and proCASP10 were purified as reported previously[10].

*Reduction and Zeba desalting*

Before gel-based ABPP analysis, all recombinant protein samples were first treated with 5 mM DTT for 15 minutes followed by buffer exchange using Zeba desalting columns to ensure that all caspase protein was fully reduced prior to analysis. Samples were buffer exchanged using the Zeba Desalting Columns 7K MWCO (Thermo Scientific, 89882) following the manufacturer's desalting procedure. Briefly, Zeba columns were first harvested by centrifugation in a 1.5 mL microcentrifuge tube using a variable-speed bench-top microcentrifuge at 1,500 x *g* for 1 minute. The columns were prepared by first loading 300 uL of PBS followed by centrifugation at 1,500 x *g* for 1 minute. The washes were repeated 2 more times. 100 uL of treated or non-treated samples were then loaded onto the desalting column, harvested by centrifugation at 1,500 x *g* for 1 minute and collected in a fresh 1.5 mL microcentrifuge tube.

*Gel based ABPP.*

Recombinant proteins were first subjected to *Reduction and Zeba desalting*. Subsequently recombinant proteins were added to 1 mg/mL Jurkat cell lysates to a final concentration of 2 µM recombinant protein. 25 µL of the cell lysate-recombinant protein mixture was then treated with 100 µM or 10 µM of electrophiles (**P01-P06, 5-16**) for 1h at ambient conditions or 30℃. The samples were then incubated for 1h with 1 µL of 250 µM click probes (**IAA, KB18, KB19, KB61, 1-4**) at a final concentration of 10 µM. Samples were then subjected to either click conjugation to rhodamine azide in 3 uL of click mix containing TBTA (1.5 µL of 1.7 mM for a final concentration of 100 µM), $CuSO_4$ (0.5 µL of 50 mM for a final concentration of 1 mM), Rhodamine-azide (0.5 µL of 1.25 mM for a final concentration of 25 µM), and TCEP (0.5 of µL 50 mM for a final concentration of 1 mM) or activity-based probe **Rho-DEVD-AOMK (2** µM **)** Next, 10 µL of 4x laemmli loading dye (BioRad, 1610747) was added and the samples were denatured at 95°C for 5 minutes. Samples were resolved by SDS-PAGE and imaged using a BioRad ChemiDoc Imaging System. Coomassie InstantBlue (Fisher Scientific, ISB1L) was used for visualization of protein loading.

*TEV activation Gel based ABPP*

Purified recombinant caspase-2 TEV cleavable constructs (proCASP2TEV and proCASP2TEV_C37A) at 2 µM final concentration in 1 mg/mL Jurkat lysates were first treated with either compound **3** or **Rho-DEVD-AOMK** at 10 µM final concentration for 1h at ambient conditions. proCASPTEV constructs were then activated with TEV protease (2 mg/mL stock) at increasing final concentrations (0 µM, 0.1 µM, 0.5 µM, 1.0 µM, 2.5 µM, 5.0 µM) for 1h at ambient conditions. Samples treated with compound **3** were then subjected to 'click' conjugation to rhodamine azide as prepared in the "Gel based ABPP" section and visualized by SDS-PAGE in-gel fluorescence using a BioRad ChemiDoc Imaging System. Coomassie InstantBlue was used for visualization of protein loading.

*Determining in vitro apparent $IC_{50}$ values*
Purified recombinant caspase-2 protein was spiked into 1 mg/mL cell lysates to make a final concentration of 2 µM recombinant protein. This mixture was aliquoted into 25 µL samples and treated with 1 µL of electrophile (final concentration of 0.1 µM, 1.0 µM, 2.5 µM, 5.0 µM, 10 µM, 20 µM, 50 µM, 75 µM, and 100 µM) for one 1h at ambient temperature. These samples were then treated with 1 µL of 250 µM compound **3** to a final concentration of 10 µM. Following incubation with the alkyne probe, 3 µL click mix was added as prepared in the "Gel based ABPP" sectionand reaction was allowed to proceed at ambient temperature for 1h. Next, 10 µL of 4x laemmli loading dye was added and the samples were denatured at 95℃ for 5 minutes. Samples were resolved by SDS-PAGE and imaged using a BioRad ChemiDoc Imaging System. Images were analyzed using ImageJ and plotted in GraphPad Prism 9 to obtain a final $IC_{50}$ value. The percentage of labeling was determined by quantifying the integrated optical intensity of each individual band after subtracting the background signal for each fluorescent gel following the "Quantification of Gel Bands by ImageJ" protocol. Nonlinear regression analysis was used to determine the IC50 values from the dose-response curve generated in GraphPad Prism 9.

*Quantification of Gel Bands by an Image J*
Raw TIFF files were opened on ImageJ[16]. Bands were selected with a rectangle as region of interest (ROI). After all bands were selected and registered, the Band/Peak Quantification macro was started. Selected lanes were plotted followed by manually drawing baseline for each selected lane. Quantification of peak area was then determined using the wand tool.

*Enzyme activity assay*
Enzymatic activity of pro-caspase and active-caspase constructs (1 µM recombinant protein, 5 mM DTT and 333 mM citrate in PBS) were determined using the fluorescent caspase-2 substrate Ac-VDVAD-AFC(Cayman Chemicals, item number 37351) at 10 µM final concentration. Caspase and substrate buffers were mixed and immediately ran on the plate reader. 7-amino-4-trifluoromethylcoumarin (AFC) released by substrate cleavage was detected at $\lambda_{ex}$ = 400 nm and $\lambda_{em}$ = 505 nm using a multimodal Synergy H1 microplate reader (BioTek). Reads were collected every minute after substrate addition. Percent activities were calculated from the slope of the linear range determined from the reaction progress curves. Graphpad Prism (Version Prism 9.5.1) was used to obtain $K_m$ and $k_{cat}$ values by fitting reaction velocities into the Michaelis-Menten equation using varied fluorogenic substrate Ac-VDVAD-AFC concentrations of 100, 75, 50, 20, 10, 5, 2.5 and 1.0 µM.

*Compound inhibition assays*
Recombinant protein was first treated with 5 mM DTT for 15 minutes followed by buffer exchange using Zeba desalting columns (Thermo Scientific, 89882) as described in *Reduction and Zeba desalting.* Subsequently, 1 µM recombinant caspase-2, caspase-3, caspase-8 and caspase-10

were treated with electrophiles at indicated final concentrations (i.e. 10 µM, 100 µM, etc.) for 1h (2h for figure 2C, 2D, and 2F) at 30°C. Samples were then treated with 10 µM fluorogenic caspase-3/-7 substrate, Ac-VDVAD-AFC (Cayman , 37351) and 333 mM citrate in PBS. Reaction progress curves from cleaved substrate fluorescence were then monitored by a multimodal plate reader as described in *Enzyme activity assay.*

*Size Exclusion*

Ni-NTA agarose-purified recombinant caspase-2 proteins were subjected to fast protein liquid chromatography (FPLC) using a prepackaged gel filtration column consisting of a HiLoad Superdex 75 prep grade (pg) (GE Healthcare Life Sciences) column and a NGC medium-pressure chromatography system and BioFrac collector from Bio-Rad (NGC10 Chromatography System). Column was pre-equilibrated with 1 column volume (120 mL total volume) degassed Size exclusion buffer (20 mM Tris, 50 mM NaCl, 1 mM DTT) following which analytes loaded using a 1 mL sample loop were subjected chromatography (0.5 mL/min). 1mL fractions were collected for 1 column equivalent volume (120 mL total volume). For compound treatment, recombinant protein at 0.3 mg/mL was first treated with 100 µM of compound (**P01**, **3**, **11**, **12**) at 30 ℃ for 1 h. Percent of component was determined by area-under-the-curve (AUC) calculations (AUC integrated measurement tool from GraphPad Prism 9) of the UV absorbance spectra (mAU) at 280 nm for the dimeric gel filtration column volume fractions (0.30 - 0.45) and monomeric fractions (0.48 - 0.55) from each recombinant protein sample (0.3 mg/mL).

*CellTiter-Glo® Cell viability analysis*

Jurkat cells in complete RPMI 1640 media (100 µL of 1.0 x$10^6$ cells/mL) were added to a 96-well white/clear bottom tissue culture treated plate using a multichannel pipette. To each well, samples were treated with either vehicle (2 µL of DMSO), or compound (2 µL of 1.25 mM stock in DMSO for a final concentration of 25 µM for **P01** and **5**, or 2 µL of 5 mM stock for a final concentration of 100 µM for **8**, **9**, **11**, and **12**). Following 1h incubation, staurosporine (STS; Fisher Scientific; S-9300-10MG), or mega FasL (AdipoGen; AG-40B-0130-C010) were added to reach concentrations of 1 µM, or 50 ng/mL, respectively. Equal volume of DMSO or buffer was used for vehicle-treated samples. After incubation for either 4h in the case of the FasL and STS or 10h for etoposide, 100 µL of CellTiter-Glo® 2.0 Cell Viability Assay (Promega, G9242) was added to each well and the relative luminescence (RLU) measured. Percent cell viability was calculated relative to the DMSO treated Jurkat cells. Samples were analyzed in three biological replicates.

*Western blotting*

To 8 mL of Jurkat cells at a density of 1.3 x $10^6$ cells/mL in fresh complete RPMI 1640 media was added 2 mL of compound stocks prediluted in FBS-free RPMI media. For compounds assayed at 100 µM, 20 µL of 50 mM stock added to the 2 mL of FBS-free media, and for compounds assayed at 25 µM, 5 µL of 50 mM stock added to the 2 mL of FBS-free media. After compound addition, the cell suspensions were incubated for 1h at 37℃ following which 4 µL of 25 mM stock of etoposide (ETO) [Cayman Chemical, 33419-42-0] or DMSO for a final concentration of 10 µM. Following the 1h compound incubation + 10h ETO incubation, the cells were harvested (1,400 x *g*, 3 min), washed 2x with cold PBS (1,400 x *g*, 3 min), and cell pellets were collected in 1.5 mL microcentrifuge tubes. Cell pellets were lysed using 3-[(3-Cholamidopropyl) dimethylammonio]-1-propanesulfonate (CHAPS) buffer [0.3% CHAPS in PBS; 100 µL; 30 minutes on ice] and centrifuged (1,400*g*, 10 min). The clarified supernatant was then transferred to a new tube. The concentration of the cell lysates were determined using DC Protein Assay (BioRad) and the samples were diluted to 3.8 mg/mL. Samples were then treated with 4x SDS loading buffer and boiled at 95 ℃ for 5 min. The samples were resolved by SDS-PAGE gel followed by wet transfer to an activated PVDF membrane (35 Volts for 90 min at 4°C). Membranes were then blocked in 5% milk in 1x Tris-Buffer Saline, 0.1% Tween® 20 (TBST) for 1h and probed with the indicated antibodies diluted in 5% milk in TBST. The primary antibodies and dilutions used are as follows: anti-PARP (Cell Signaling, 9532, 1:1,000), anti-CASP3 (Cell Signaling, 9662, 1:1,000), anti-CASP8 (cleaved form, Cell Signaling, 9746, 1:1,000), anti-CASP8 (pro-form, Cell Signaling, 4970, 1:1,000), anti-actin (Cell Signaling, 3700, 1:5,000), anti-CASP2 (ProteinTech, 10436-1-AP, 1:1,000), and anti-CASP9 (Cell Signaling, 9502, 1:1,000). Blots were incubated with primary antibody overnight at 4 °C with rocking, then washed (3x5 min TBST), and incubated with secondary antibody (IRDye 800CW or IRDye 647CW, 1:10,000) for 1h at ambient temperature. Blots were washed (3x5 min TBST) and visualized on a BioRad ChemiDoc Imaging System.

*Statistics*
Statistical significance was calculated with unpaired two-tailed Student's t-test using GraphPad Prism 9. Data are shown as mean ± SDEV. *P* values of <0.05 were considered significant.

### (D) Chemistry Methods

**General Methods.** All solution-phase reactions were performed in oven-dried glassware under an atmosphere of dry $N_2$ except where water was used as a solvent. Reactions were monitored by thin-layer chromatography, and plates were visualized by fluorescence quenching under UV light or by staining with iodine.

For compounds that required purifications, purifications were done via column chromatography. silica gel flash column chromatography. Silica gel P60 (SiliCycle) as the stationary phase. Typical stationary phase column loadings was column height: 250 mm, diameter: 100 mm, 100-200 mesh silica gel Typical eluents used were hexanes/Ethylacetate, Methanol/$CH_2Cl_2$, Petroleum ether/Ethyl acetate or hexanes/$CH_2Cl_2$.

Other reagents were purchased from Sigma-Aldrich (St. Louis, MO), Alfa Aesar (Ward Hill, MA), EMD Millipore (Billerica, MA), Fisher Scientific (Hampton, NH), Oakwood Chemical (West Columbia, SC), Combi-blocks (San Diego, CA) and Cayman Chemical (Ann Arbor, MI) and used without further purification. $^1$H NMR and $^{13}$C NMR spectra for characterization of new compounds and monitoring reactions were collected in $CDCl_3$, $CD_3OD$, $CD_6CO$ or DMSO-$d_6$ (Cambridge Isotope Laboratories, Cambridge, MA) on a Bruker AV 400 MHz spectrometer or Bruker AV 500 MHz in the Department of Chemistry & Biochemistry at University of California, Los Angeles. All chemical shifts were reported in the standard notation of parts per million using the peak of residual proton signals of the deuterated solvent as an internal reference. Coupling constant units are in Hertz (Hz). Splitting patterns were indicated as follows: br, broad; s, singlet; d, doublet; t, triplet; q, quartet; m, multiplet; dd, doublet of doublets; dt, doublet of triplets. Low-resolution mass spectroscopy was performed on an Agilent Technologies InfinityLab LC/MSD single quadrupole LC/MS (ESI source).

**Purchased electrophilic fragments (P01-P06) were obtained from the following vendors:** **P01** (Fisher Scientific, AC334590050), **P02** (Fisher Scientific, P06105G), **P03** (Combi-blocks, QB-

5712), (**P04** (Combi-blocks, ST-8644), **P05** (Combi-blocks, QC-2990) and **P06** (Combi-blocks, QF-4549)

**General procedure 1:**

To an oven-dried flask with a stir bar was charged the aldehyde (1 equiv.), anhydrous $CH_2Cl_2$ , the amine (1.1 equiv.), and acetic acid (1.2 equiv.). Sodium triacetoxyborohydride was added (1.5 equiv.) in three separate portions over the course of 1.5 h. The reaction was allowed to stir for 16 h. under nitrogen. The reaction was then quenched with water and extracted three times with DCM. Organic layers were washed with brine, dried over anhydrous $Na_2SO_4$, and concentrated *in vacuo* to give the crude product, which was purified by silica gel flash column chromatography to give the titled compound.

**General procedure 2:**

To a solution of the amine (1.1 equiv.) and the aldehyde (1.0 equiv.) in anhydrous MeOH (1.0 mL) was added AcOH (1.0 equiv.). The reaction was stirred at 25 °C for 3 h. $NaBH_3CN$ (2.0 equiv.) was added at 0 °C. The reaction was stirred at 25 °C for another 1 h. LCMS showed the reaction was completed. The residue was poured into ice-water. The aqueous phase was extracted three times with ethyl acetate.The combined organic phase was washed with brine, dried with anhydrous $Na_2SO_4$, filtered and concentrated in vacuo and purified by silica gel flash column chromatography to give the titled compound.

**General procedure 3:**

The amine (1 equiv.) was added to a round bottom flask with a stir bar and dissolved in DCM with stirring. Triethylamine (2 equiv.) was added and the reaction was cooled to 0 ˚C. Chloroacetyl chloride (2 equiv.) was added dropwise to the flask, and the reaction was allowed to warm to room temperature as it stirred for 2 h. The reaction was washed with water, extracted with ethyl acetate, dried over $Na_2SO_4$ and purified by silica gel flash column chromatography to give the titled compound.

**General procedure 4:**

The amine (1 equiv.) was added to a round bottom flask with a stir bar and dissolved in DCM. Triethylamine (1 equiv.) was added, and the reaction was cooled to 0 ˚C. Acryloyl chloride (2 equiv.) was added dropwise to the flask, and the reaction was allowed to warm to room temperature as it stirred for 2 h. The reaction was washed with water, extracted with ethyl acetate, dried over $Na_2SO_4$ and purified silica gel flash column chromatography to give the titled compound.

**General procedure 5:**

To a round bottom flask equipped with a reflux condenser, the ester (1 equiv.) was dissolved in 3M HCl and allowed to reflux for 24 h. The reaction was cooled, extracted with three times with

DCM. The combined organic layers were washed with brine, dried over $Na_2SO_4$, , filtered and concentrated *in vacuo* to give the titled compound.

**General procedure 6:**

Aryl iodide (1 equiv.) , $Cs_2CO_3$ (2 equiv.) , $Pd(PPh_3)_4$ (0.05 equiv.), and CuI (0.05 equiv.) were added to an over-dried pressure tube and dissolved in THF, the reaction was heated to 80 °C and then propiolate (2 equiv.) was added dropwise and the mixture was allowed to stir for 24 h at 80 °C. The crude mixture was then diluted with DCM and filtered through a bed of celite. The mixture was then washed with water, extracted with DCM, dried with anhydrous sodium sulfate, filtered, and concentrated *in vacuo*. The product was then purified by silica gel flash column chromatography to give the titled compound.

**General procedure 7:**

To a round bottom flask was added the carboxylic acid (1 equiv.) and the alcohol (1 equiv.) which were then dissolved in $CH_2Cl_2$ and cooled in an ice bath. DCC (1.05 equiv.) dissolved in $CH_2Cl_2$ was added to the above solution in one portion under vigorous stirring and stirred for 20 min. DMAP (0.05 equiv.) was then added and the reaction mixture was allowed to warm to room temperature and stirred for 19 h. The precipitated N,N-dicyclohexylurea was filtered off and the solvent was concentrated. The crude material was then purified via silica gel flash column chromatography to give the titled compound.

**Synthesis of *N*-(4-ethynylbenzyl)aniline (S01)**

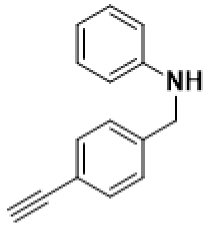

Prepared according to general procedure 1, using Aniline (737 ***µ***l, 8.06 mmol) as the amine; 4-ethynylbenzaldehyde (0.937 g, 7.2 mmol) as the aldehyde . Product: off white solid (748 mg, 50%)

**$^1$H NMR (400 MHz, $CDCl_3$) :** δ 7.39 (d, J = 8.3 Hz, 2H), 7.36 – 7.28 (m, 3H), 7.18 (d, J = 8.4 Hz, 2H), 6.99 (dd, J = 7.9, 1.5 Hz, 2H), 6.43 (d, J = 18.8 Hz, 1H), 6.09 – 5.99 (m, 1H), 5.56 (d, J = 12.3 Hz, 1H), 4.96 (s, 2H), 3.05 (s, 1H). **$^{13}$C NMR (101 MHz, $CDCl_3$) :** δ 165.64, 141.61, 138.19, 132.23, 129.56, 128.71, 128.43, 128.29, 127.99, 121.14, 83.47, 52.93.

**LC-MS** (ESI, m/z): calcd for $C_{15}H_{14}N^+$ $[M+H]^+$ 208.1; found 208.1

**Synthesis of 2-chloro-*N*-(4-ethynylbenzyl)-*N*-phenylacetamide (1)**

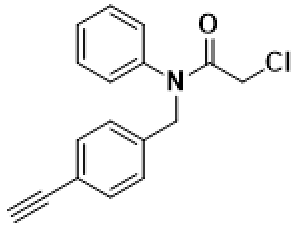

Prepared according to general procedure 3, using N-(4-ethynylbenzyl)aniline (500 mg, 1 eq, 2.4 mmol) as the amine source. Yield: 479.1 mg, 70%.

**$^1$H NMR (400 MHz, $CDCl_3$)** : δ 7.40 (d, J = 8.2 Hz, 2H), 7.35 (d, J = 1.5 Hz, 3H), 7.16 (d, J = 8.2 Hz, 2H), 7.01 (dd, J = 5.7, 3.8 Hz, 2H), 4.88 (s,

2H), 3.84 (s, 2H), 3.07 (s, 1H). **$^{13}$C NMR (101 MHz, CDCl$_3$)** : δ 166.32, 140.66, 137.37, 132.31, 129.99, 128.97, 128.21, 121.51, 83.32, 53.48, 41.95.

**HRMS (ESI-TOF, m/z)**: calcd for $C_{17}H_{15}ClNO^+$ [M+H]$^+$ 284.0842; found 284.0836

**Synthesis of 1-((4-ethynylphenyl)sulfonyl)piperazine (S02)**

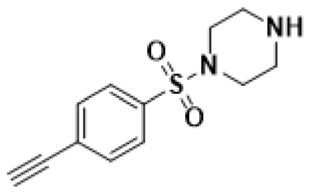

4-(4-Ethynylbenzenesulfonyl)-piperazine-1-carboxylic acid tert-butyl ester (100 mg, 0.285 mmol) was added to a round bottom flask with a stir bar, and dissolved in DCM (3 mL) with stirring and cooled to 0 ˚C. Trifluoroacetic acid (3 mL) was added to the reaction dropwise and the reaction was allowed to warm to room temperature as it stirred for 1h. The reaction was washed with water, extracted with ethyl acetate, dried over anhydrous $Na_2SO_4$ worked up to yield 59.4 mg of pure 1-((4-ethynylphenyl)sulfonyl)piperazine (59.4 mg, 99%).

**LC-MS** (ESI, m/z): calcd for $C_{12}H_{15}N_2O_2S^+$ [M+H]$^+$ 251.09; found 251.1

**Synthesis of 2-chloro-1-(4-((4-ethynylphenyl)sulfonyl)piperazin-1-yl)ethan-1-on (2)**

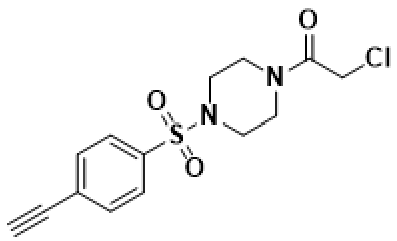

Prepared according to general procedure 3, using 4-(4-Ethynylbenzenesulfonyl)-piperazine-1-carboxylic acid tert-butyl ester (35 mg, 1 eq, 0.140 mmol) as the amine source. Product: off white solid (30.5 mg, 66%)

**$^1$H NMR (400 MHz, CDCl$_3$) :** δ 7.72 – 7.69 (m, 2H), 7.65 (d, J = 8.6 Hz, 2H), 4.00 (s, 2H), 3.67 (dt, J = 49.0, 4.9 Hz, 4H), 3.29 (s, 1H), 3.08 (dt, J = 24.8, 4.8 Hz, 4H). **$^{13}$C NMR (101 MHz, CDCl$_3$) :** δ 165.12, 135.33, 132.90, 127.59, 81.70, 81.32, 45.93, 45.75, 45.57, 41.43, 40.50.

**HRMS (ESI-TOF, m/z)**: calcd for $C_{14}H_{16}ClN_2O_3S^+$ [M+H]$^+$ 327.0492; found 327.0489

**Synthesis of *N*-(4-ethynylbenzyl)-*N*-phenylacrylamide (3)**

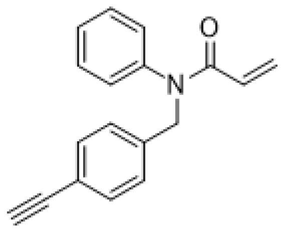

Prepared according to general procedure 4, using N-(4-ethynylbenzyl)aniline, **S01** (500 mg, 1 eq, 2.4 mmol) as the amine source. Yield: 116.2 mg , 50%.

**$^1$H NMR (400 MHz, CDCl$_3$)** : δ 7.39 (d, J = 8.3 Hz, 2H), 7.36 – 7.30 (m, 3H), 7.18 (d, J = 8.4 Hz, 2H), 7.03 – 6.95 (m, 2H), 6.44 (d, J = 2.0 Hz, 1H), 6.03 (dd, J = 16.8, 10.3 Hz, 1H), 5.56 (dd, J = 10.2, 2.0 Hz, 1H), 4.96 (s, 2H), 3.05 (s, 1H).**$^{13}$C NMR (101 MHz, CDCl$_3$)** : δ 165.64, 141.61, 138.19, 132.23, 129.56, 128.71, 128.43, 128.29, 128.24, 127.99, 121.14, 83.47, 52.93.

**HRMS (ESI-TOF, m/z)**: calcd for $C_{17}H_{15}ClNO^+$ [M+H]$^+$ 262.1231; found 262.1227

**Synthesis of 1-(4-((4-ethynylphenyl)sulfonyl)piperazin-1-yl)prop-2-en-1-one (4)**

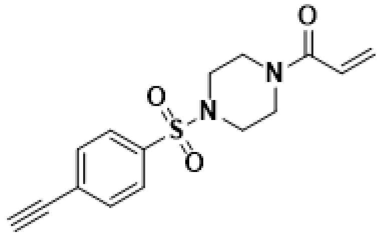

Prepared according to general procedure 4, using 1-((4-ethynylphenyl)sulfonyl)piperazine,**(S02)** (25.6 mg, 1 eq, 0.102 mmol) as the amine source. Product: off white solid (19 mg, 60%)

**$^{1}$H NMR (400 MHz, $CDCl_3$) :** δ 7.70 (d, J = 8.6 Hz, 2H), 7.64 (d, J = 8.6 Hz, 2H), 6.46 (dd, J = 16.8, 10.5 Hz, 1H), 6.25 (dd, J = 16.8, 1.7 Hz, 1H), 5.69 (s, 1H), 3.70 (d, J = 50.5 Hz, 4H), 3.28 (s, 1H), 3.05 (s, 4H). **$^{13}$C NMR (101 MHz, $CDCl_3$) :** δ 165.34, 135.29, 129.00, 127.62, 127.47, 126.72, 81.72, 81.24.

**HRMS (ESI-TOF, m/z)**: calcd for $C_{15}H_{17}N_2O_3S^+$ $[M+H]^+$ 305.0960; found 305.0953

**Synthesis of isopropyl 3-phenylpropiolate (5)**

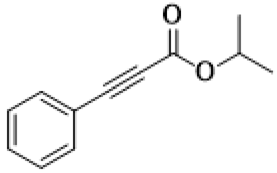

Prepared according to general procedure 7 using 3-phenylpropiolic acid (151 mg, 1.03 mmol) and propan-2-ol (61.9 mg, 1.03 mmol) as the carboxylic acid and alcohol respectively. Yield: 152 mg, 84%.

**$^{1}$H NMR (400 MHz, $CDCl_3$)** 1H NMR (400 MHz, Chloroform-d) δ 7.58 (dd, J = 8.3, 1.3 Hz, 2H), 7.47 – 7.41 (m, 1H), 7.40 – 7.33 (m, 2H), 5.16 (p, J = 6.3 Hz, 1H), 1.34 (d, J = 6.3 Hz, 6H).**$^{13}$C NMR (101 MHz, $CDCl_3$)** δ 153.67, 132.95, 130.52, 128.54, 119.75, 85.65, 81.07, 70.04, 21.74.

**HRMS (ESI-TOF, m/z)**: calcd for $C_{12}H_{13}O_2^+$ $[M+H]^+$ 189.0915; found 189.0927

**Synthesis of tert-butyl 3-(pyridin-4-yl)propiolate (6)**

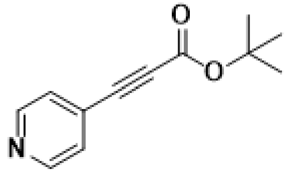

Prepared according to general procedure 6 using 4-iodopyridine (40 mg, 0.2mmol) and tert-butyl propiolate (50 mg, 0.4 mmol) as the aryliodide and alkyl propiolate respectively. Yield: 8 mg, 20%.

**$^{1}$H NMR (400 MHz, $CDCl_3$)** δ 8.09 (d, J = 8.4 Hz, 2H), 7.65 (d, J = 8.4 Hz, 2H), 1.55 (s, 9H). **$^{13}$C NMR (101 MHz, DMSO-$d_6$)** δ 151.11 , 150.04 , 132.67 , 76.37 , 75.71 , 27.25 .

**HRMS (ESI-TOF, m/z)**: calcd for $C_{12}H_{14}NO_2^+$ $[M+H]^+$ 204.1025; found 204.1020

**Synthesis of methyl 3-(quinolin-6-yl)propiolate (7)**

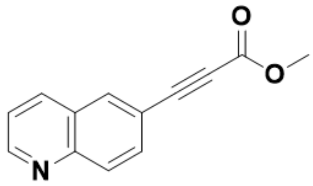

Prepared according to general procedure 6 using 6-iodoquinoline (40 mg, 0.2 mmol) and methyl propiolate (50 mg, 0.4 mmol) as the aryliodide and alkyl propiolate respectively. Yield: 4.2 mg (14%).

**$^{1}$H NMR (400 MHz, $CDCl_3$)** δ 8.98 (s, 1H), 8.17 (s, 1H), 8.13 (d, J = 5.9 Hz, 2H), 7.82 (d, J = 10.5 Hz, 1H), 7.53 – 7.45 (m, 1H), 3.87 (s, 3H).

$^{13}$**C NMR (101 MHz, $CDCl_3$)** δ 154.25, 151.74, 136.57, 133.94, 132.25, 129.82, 122.19, 118.02, 85.58, 81.27, 52.97.

**HRMS (ESI-TOF, m/z)**: calcd for $C_{13}H_{10}NO_2^+$ $[M+H]^+$ 212.0712; found 212.0718

**Synthesis of methyl 3-(9H-carbazol-3-yl)propiolate (8)**

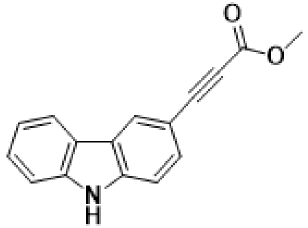

Prepared according to general procedure 6 using 3-iodo-9H-carbazole (60 mg, 0.2 mmol) and methyl propiolate (30 mg, 0.4 mmol) as the aryliodide and alkyl propiolate respectively. Yield: 9 mg, 18%

$^{1}$**H NMR (400 MHz, $CDCl_3$)** 8.40 (s, 1H), 8.03 (d, J = 7.8 Hz, 1H), 7.67 (dd, J = 8.6, 1.7 Hz, 1H), 7.45 (ddd, J = 8.3, 7.3, 1.2 Hz, 1H), 7.04 (d, J = 8.6 Hz, 1H), 6.91 (s, 1H), 6.13 (s, 1H), 3.75 (s, 3H).

$^{13}$**C NMR (101 MHz, $CDCl_3$)** δ 140.67, 139.89, 134.31, 134.16, 129.22, 127.44, 126.79, 125.98, 122.21, 120.73, 120.50, 111.83, 109.89, 82.96, 52.81.

**HRMS (ESI-TOF, m/z)**: calcd for $C_{16}H_{12}O_2^+$ $[M+H]^+$ 250.0868; found 250.0879

**Synthesis of methyl 4-((phenylamino)methyl)benzoate (S03)**

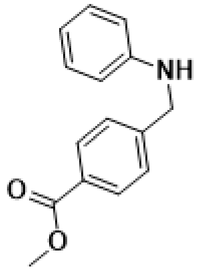

Prepared according to general procedure 1, using Aniline (1.25 mL, 13.6 mmol) as the amine; 4-methylformylbenzoate (2.00 g, 12.2 mmol) as the aldehyde . Product: brown oil. Yield: 2.79 g, 95%.

$^{1}$**H NMR (400 MHz, $CDCl_3$) :** δ 8.01 (d, J = 8.3 Hz, 2H), 7.44 (d, J = 8.4 Hz, 2H), 7.18 (dd, J = 8.5, 7.4 Hz, 2H), 6.74 (t, J = 7.3 Hz, 1H), 6.66 – 6.60 (m, 2H), 4.41 (s, 2H), 3.91 (s, 3H). $^{13}$**C NMR (400 MHz, $CDCl_3$) :** δ 166.97, 147.76, 144.98, 129.98, 129.34, 129.13, 127.18, 117.91, 112.96, 52.10, 48.04.

**LC-MS** (ESI, m/z): calcd for $C_{15}H_{16}NO_2^+$ $[M+H]^+$ 242.1; found 242.1

**Synthesis of methyl 4-((N-phenylacrylamido)methyl)benzoate (9)**

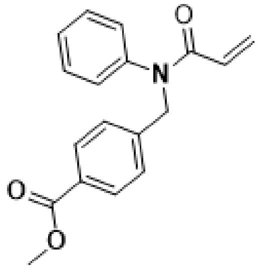

Prepared according to general procedure 4 – under nitrogen – using 4-((phenylamino)methyl)benzoate (1.47g, 6.10 mmol) as the amine source. Product: yellow oil. Yield: 1.69 g, 94%.

$^{1}$**H NMR (400 MHz, $CDCl_3$) :** d 7.94 (d, *J* = 8.3 Hz, 2H), 7.26-7.35 (m, 5H), 6.99 (dd, *J* = 7.9, 2.2 Hz , 2H), 6.44 ( *J* = 16.7, 2.0 Hz), 6.04 (dd, *J* = 16.8, 6.5 Hz, 1H), 5.5 (dd, *J* = 10.3, 2 Hz, 1H), 5.0 (s, 2H), 3.9 (s, 3H). $^{13}$**C NMR (400 MHz, $CDCl_3$) :** δ 169.79, 166.85, 141.57, 140.53, 130.03, 129.85, 129.57, 129.02, 128.88, 128.09, 52.18, 41.81.

**HRMS (ESI-TOF, m/z)**: calcd for $C_{18}H_{18}NO_3^+$ $[M+H]^+$ 296.1287; found 296.1279

**Synthesis of 5-chloro-N-(pyridin-2-ylmethyl)pyrimidin-2-amine (S04)**

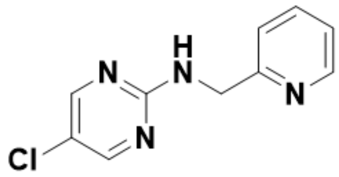

Prepared according to general procedure 2, using 5-chloropyrimidin-2-amine (266.1 mg, 1.1 eq, 2.05 mmol) as the amine; picolinaldehyde (200 mg,1 eq, 1.87 mmol) as the aldehyde . Product: off white (356.1 mg, 86.4%)

**$^1$H NMR (400 MHz, CDCl$_3$) :** δ 8.58 (s, 1H), 8.23 (s, 2H), 7.77 (s, 1H), 7.42 (d, J = 7.9 Hz, 1H), 7.33 – 7.27 (m, 1H), 6.45 (s, 1H), 4.78 (s, 2H). **$^{13}$C NMR (101 MHz, CDCl$_3$) :** δ 160.11, 156.69, 156.13, 147.46, 138.00, 122.35, 119.35, 77.16, 45.88.

**LC-MS** (ESI, m/z): calcd for $C_{10}H_{10}ClN_4^+$ [M+H]$^+$ 221.1; found 221.1

**Synthesis of N-(5-chloropyrimidin-2-yl)-N-(pyridin-2-ylmethyl)acrylamide (10)**

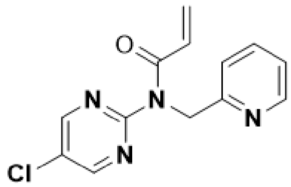

Prepared according to general procedure 4, using 5-chloro-N-(pyridin-2-ylmethyl)pyrimidin-2-amine, **S04** (200 mg, 1 eq, 0.91 mmol) as the amine source. Product: pale yellowish oil (152 mg, 61%)

**$^1$H NMR (400 MHz, CDCl$_3$) :** δ 8.57 (s, 2H), 8.52 (s, 1H), 7.63 (d, J = 17.2 Hz, 1H), 7.28 (s, 2H), 7.14 (s, 1H), 6.80 (dd, J = 16.8, 10.3 Hz, 1H), 6.52 – 6.44 (m, 1H), 5.75 (dd, J = 10.3, 1.7 Hz, 1H), 5.51 (s, 2H). **$^{13}$C NMR (101 MHz, CDCl$_3$) :** δ 167.55, 157.11, 156.38, 148.91, 136.99, 130.48, 128.05, 126.39, 122.12, 121.31, 51.42.

**HRMS (ESI-TOF, m/z)**: calcd for $C_{13}H_{12}ClN_4O^+$ [M+H]$^+$ 275.0670; found 275.0698

**Synthesis of tert-butyl 5-formyl-1H-benzo[d]imidazole-1-carboxylate (S05)**

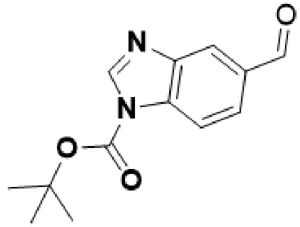

To a mixture of 3H-benzimidazole-5-carbaldehyde (500.00 mg, 3.42 mmol, 1.00 *eq*) and $Et_3N$ (692.37 mg, 6.84 mmol, 948.45 uL, 2.00 *eq*) in DCM (5.00 mL) was added $Boc_2O$ (1.49 g, 6.84 mmol, 1.57 mL, 2.00 *eq*) in one portion at 25 °C .The mixture was stirred at 25 °C for 16h. TLC (PE:EA=5:1,Rf =0.43) and LCMS showed the reaction was completed. The mixture was concentrated in a vacuum. The reaction mixture was filtered and the filterate was concentrated. The residue was purified by silica gel flash column chromatography to yield the titled compound (800 mg, 95%)

**$^1$H NMR (400 MHz, $CDCl_3$) :** δ 10.11 (s, 1H), 8.53 (s, 1H), 8.31 – 8.10 (m, 1H), 7.91 (dd, J = 2.2, 1.0 Hz, 1H), 1.73 (s, 9H). **$^{13}$C NMR (101 MHz, $CDCl_3$) :** δ 191.65, 147.50, 145.02, 143.79, 133.75, 133.30, 125.95, 125.45, 123.71, 121.24, 117.15, 115.06, 86.70, 28.06.

**LC-MS** (ESI, m/z): calcd for $C_{13}H_{15}N_2O_3^+$ [M+H]$^+$ 247.1; found 247.1

**Synthesis of tert-butyl 5-((phenylamino)methyl)-1H-benzo[d]imidazole-1-carboxylate (S06)**

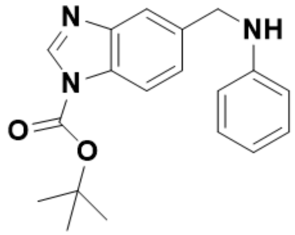

Prepared according to general procedure 2, using aniline (74.88 mg, 804.03 umol, 73.41 uL) as the amine; tert-butyl 6-formylbenzimidazole-1-carboxylate, **S05** (180.00 mg, 730.93 umol) as the aldehyde . Product: yellow oil (240 mg, 91.4%)

**$^1$H NMR (400 MHz, $CDCl_3$) :** δ 8.40 (s, 1H), 8.03 (s, 1H), 7.73 (d, J = 8.2 Hz, 1H), 7.36 (dd, J = 8.3, 1.6 Hz, 1H), 4.82 (s, 2H), 1.69 (s, 9H). **$^{13}$C NMR (101 MHz, $CDCl_3$) :** δ 147.99, 143.55, 142.28, 138.54, 131.59, 123.53, 123.53, 120.55, 112.99, 85.75, 53.45, 28.07.

**LC-MS** (ESI, m/z): calcd for $C_{19}H_{22}N_3O_2^+$ [M+H]$^+$ 324.1; found 324.1

**Synthesis of tert-butyl 5-((N-phenylacrylamido)methyl)-1H-benzo[d]imidazole-1-carboxylate (S07)**

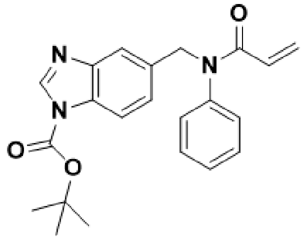

Prepared according to general procedure 4, using tert-butyl 5-((phenylamino)methyl)-1H-benzo[d]imidazole-1-carboxylate, **S06** (200 mg, 0.62 mmol) as the amine source. Product: yellow oil; and *used as is without further characterisation* (140 mg, 60%)

**LC-MS** (ESI, m/z): calcd for $C_{22}H_{24}N_3O_3^+$ [M+H]$^+$ 378.2; found 378.2

**Synthesis of N-((1H-benzo[d]imidazol-5-yl)methyl)-N-phenylacrylamide (11)**

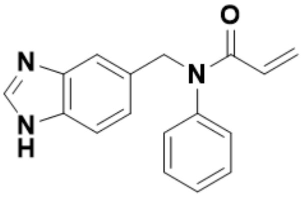

TFA (5mL.) was added to a solution of tert-butyl 5-((N-phenylacrylamido)methyl)-1H-benzo[d]imidazole-1-carboxylate, **S07** (100 mg) in DCM (20 mL) at 18° C. The resulting mixture was stirred at 18 °C for 3 h. Upon completion, the reaction mixture was concentrated in vacuo to yield the crude. To this crude was added 10 mL of saturated sodium bicarbonate solution stirred for 15 mins, next extracted into DCM and purified by purified by silica gel flash column chromatography to yield the titled compound (57mg, 73.5%).

**$^1$H NMR (400 MHz, $CDCl_3$) :** δ 7.99 (s, 1H), 7.52 (d, J = 8.7 Hz, 2H), 7.31 (s, 3H), 7.12 (s, 1H), 6.99 (s, 2H), 6.41 (d, J = 18.8 Hz, 1H), 6.13 – 5.99 (m, 1H), 5.55 (d, J = 10.3 Hz, 1H), 5.09 (s,

2H). **$^{13}$C NMR (101 MHz, CDCl$_3$) :** δ 166.27, 142.10, 141.63, 132.47, 129.96, 129.21, 128.77, 128.43, 124.17, 116.43, 115.58, 77.16, 54.07.

**HRMS (ESI-TOF, m/z)**: calcd for $C_{17}H_{16}N_3O^+$ $[M+H]^+$ 278.1248; found 278.1255

**Synthesis of tert-butyl 6-((naphthalen-2-ylamino)methyl)-1H-benzo[d]imidazole-1-carboxylate (S08)**

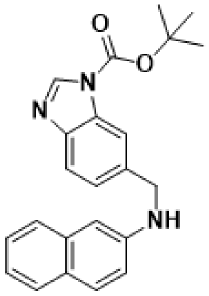

Prepared according to general procedure 2, using naphthalen-2-amine (200 mg, 1.88 mmol) as the amine; 5-formyl-1H-benzo[d]imidazole-1-carboxylate, **S05** (304.5 mg, 2.07 mmol) as the aldehyde . Product: off-white solid (268.5 mg, 88.5%)

**$^{1}$H NMR (400 MHz, CDCl$_3$) :** δ 8.42 (d, J = 8.2 Hz, 1H), 8.11 – 7.91 (m, 1H), 7.84 – 7.74 (m, 1H), 7.65 (dd, J = 11.7, 8.5 Hz, 2H), 7.57 (dd, J = 8.0, 2.7 Hz, 1H), 7.45 (dd, J = 16.1, 7.5 Hz, 1H), 7.39 – 7.30 (m, 1H), 7.19 (ddt, J = 8.1, 6.8, 1.2 Hz, 1H), 6.95 (dd, J = 8.8, 2.3 Hz, 1H), 6.85 (s, 1H), 4.58 (d, J = 6.8 Hz, 2H), 1.66 (d, J = 27.3 Hz, 9H). **$^{13}$C NMR (101 MHz, CDCl$_3$) :** δ 148.03, 144.50, 142.21, 135.12, 129.02, 127.64, 126.05, 126.02, 124.91, 123.96, 122.17, 120.70, 119.53, 117.90, 114.52, 113.36, 85.73, 48.69, 28.02.

**LC-MS** (ESI, m/z): calcd for $C_{23}H_{24}N_3O_2^+$ $[M+H]^+$ 374.1; found 374.1

**Synthesis of tert-butyl 6-((N-(naphthalen-2-yl)acrylamido)methyl)-1H-benzo[d]imidazole-1-carboxylate (S09)**

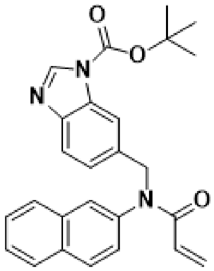

Prepared according to general procedure 4, using tert-butyl 6-((naphthalen-2-ylamino)methyl)-1H-benzo[d]imidazole-1-carboxylate, **S08** (200 mg, 0.54 mmol) as the amine source. Product: yellow oil; and *used as is without further characterisation* (180 mg, 78.6%)

**LC-MS** (ESI, m/z): calcd for $C_{26}H_{26}N_3O_3^+$ $[M+H]^+$ $C_{26}H_{26}N_3O_3^+$ ; found 428.1

**Synthesis of N-((1H-benzo[d]imidazol-6-yl)methyl)-N-(naphthalen-2-yl)acrylamide (12)**

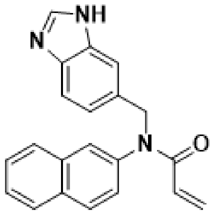

Prepared according to general procedure 2, using tert-butyl 6-((N-(naphthalen-2-yl)acrylamido)methyl)-1H-benzo[d]imidazole- 1-carboxylate, **S09** (200 mg, 1 eq, 0.84 mmol) as the amine source. Product: off-white oil (24 mg, 17.4%)

**$^{1}$H NMR (400 MHz, $CDCl_3$) :** δ 8.01 (s, 1H), 7.86 – 7.76 (m, 2H), 7.71 (d, J = 9.3 Hz, 1H), 7.64 – 7.45 (m, 5H), 7.19 – 7.09 (m, 2H), 6.44 (d, J = 18.7 Hz, 1H), 6.08 (dd, J = 16.7, 10.3 Hz, 1H), 5.52 (d, J = 12.1 Hz, 1H), 5.18 (s, 2H).

**$^{13}$C NMR (400 MHz, $CDCl_3$) :** δ 165.93, 139.07, 133.41, 132.43, 132.32, 129.62, 128.86, 128.13, 127.94, 126.87, 126.11, 123.98, 53.67.

**HRMS (ESI-TOF, m/z)**: calcd for $C_{21}H_{18}N_3O^+$ $[M+H]^+$ 328.1450; found 328.1441

**Synthesis of benzyl 3-phenylpropiolate (13)**

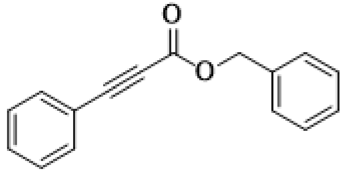

Prepared according to general procedure 7 using 3-phenylpropiolic acid (150 mg, 1.03 mmol) and benzyl alcohol (111 mg, 1.03 mmol) as the carboxylic acid and alcohol respectively. Yield: 208 mg, 86%.

**$^{1}$H NMR (400 MHz, $CDCl_3$)** δ 7.58 (dd, J = 8.3, 1.4 Hz, 2H), 7.48 – 7.33 (m, 8H), 5.27 (s, 2H).

**$^{13}$C NMR (101 MHz, $CDCl_3$)** δ 153.91, 134.93, 133.03, 130.70, 128.58, 119.56, 86.75, 80.51, 67.73.

**HRMS (ESI-TOF, m/z)**: calcd for $C_{16}H_{13}O_2^+$ $[M+H]^+$ 237.0916; found 237.0957

**Synthesis of phenyl 3-phenylpropiolate (14)**

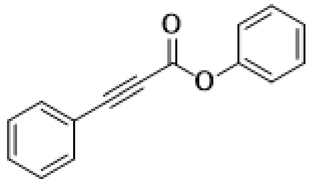

Prepared according to general procedure 7 using 3-phenylpropiolic acid (151 mg, 1.03 mmol) and phenol (96.9 mg, 1.03 mmol) as the carboxylic acid and alcohol respectively. Yield:134 mg, 83%

**$^{1}$H NMR (400 MHz, $CDCl_3$)** δ 7.64, 7.62, 7.51, 7.49, 7.47, 7.44, 7.42, 7.41, 7.40, 7.39, 7.30, 7.28, 7.26, 7.21, 7.21, 7.19, 7.18. **$^{13}$C NMR (101 MHz, $CDCl_3$)** δ 152.36, 150.17, 133.20, 131.05, 129.60, 128.68, 126.42, 121.48, 119.28..

**HRMS (ESI-TOF, m/z)**: calcd for $C_{15}H_{11}O_2^+$ $[M+H]^+$ 223.0759; found 223.0767

**Synthesis of tert-butyl 3-phenylpropiolate (15)**

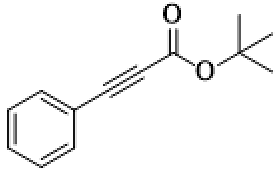

Prepared according to general procedure 6 using 3-phenylpropiolic acid (40 mg, 0.2 mmol) and tert-butyl alcohol (50 mg, 0.4 mmol) as the aryliodide and alkyl propiolate respectively. Yield: 8 mg, 20%

**$^1$H NMR (400 MHz, $CDCl_3$)** δ 7.60 – 7.54 (m, 2H), 7.45 – 7.39 (m, 1H), 7.39 – 7.32 (m, 2H), 1.54 (s, 9H). **$^{13}$C NMR (101 MHz, $CDCl_3$)** δ 153.15, 132.86, 130.30, 128.48, 119.99, 83.79, 83.52, 82.03, 28.08.

**HRMS (ESI-TOF, m/z)**: calcd for $C_{13}H_{15}O_2Na^+$ $[M+Na]^+$ 225.0892; found 225.0893

**Synthesis of methyl 3-(pyridin-4-yl)propiolate (16)**

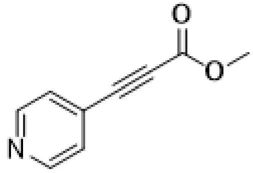

Prepared according to general procedure 6 using 4-iodopyridine (40 mg, 0.2 mmol) and methyl propiolate (50 mg, 0.4 mmol) as the aryliodide and alkyl propiolate respectively. Yield: 4.2 mg, 14%.

**$^1$H NMR (400 MHz, $CDCl_3$)** δ 8.71 – 8.63 (m, 2H), 7.46 – 7.39 (m, 2H), 3.86 (s, 3H).**$^{13}$C NMR (101 MHz, $CDCl_3$)** δ 153.65, 149.98, 127.93, 126.13, 83.53, 82.48, 53.15.

**HRMS (ESI-TOF, m/z)**: calcd for $C_9H_8NO_2^+$ $[M+H]^+$ 161.0470; found 161.0471

**(E) NMR Spectra**

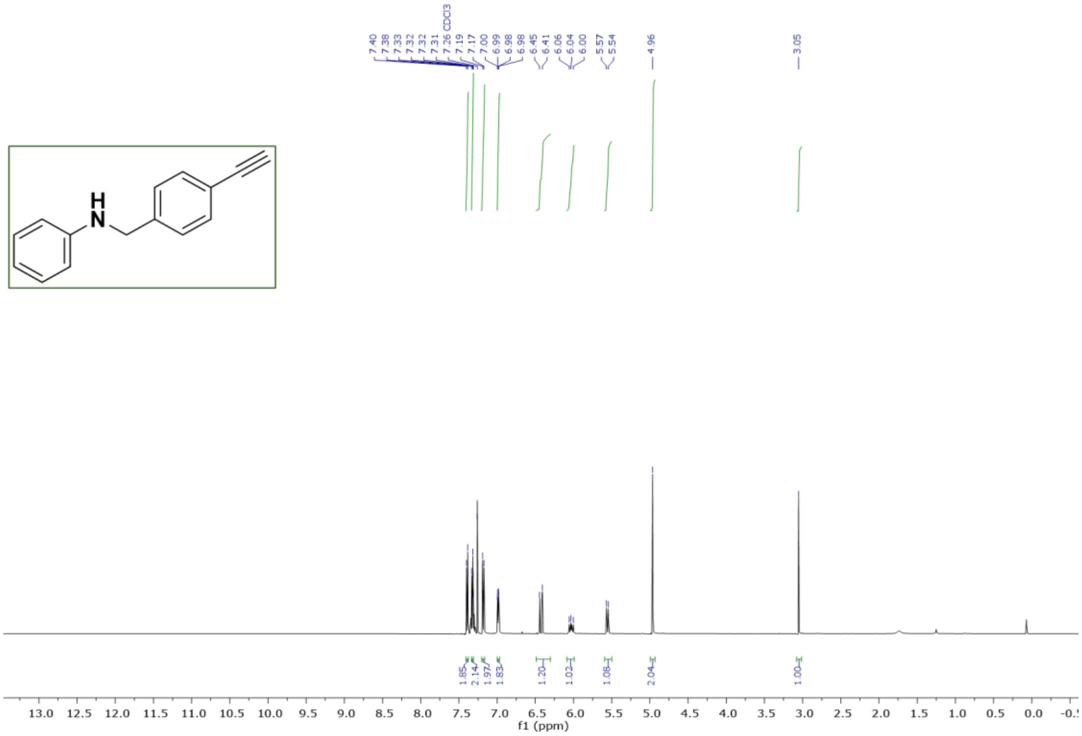

**Figure S1.** [1]H NMR of ***N*-(4-ethynylbenzyl)aniline (S01)** in $CDCl_3$

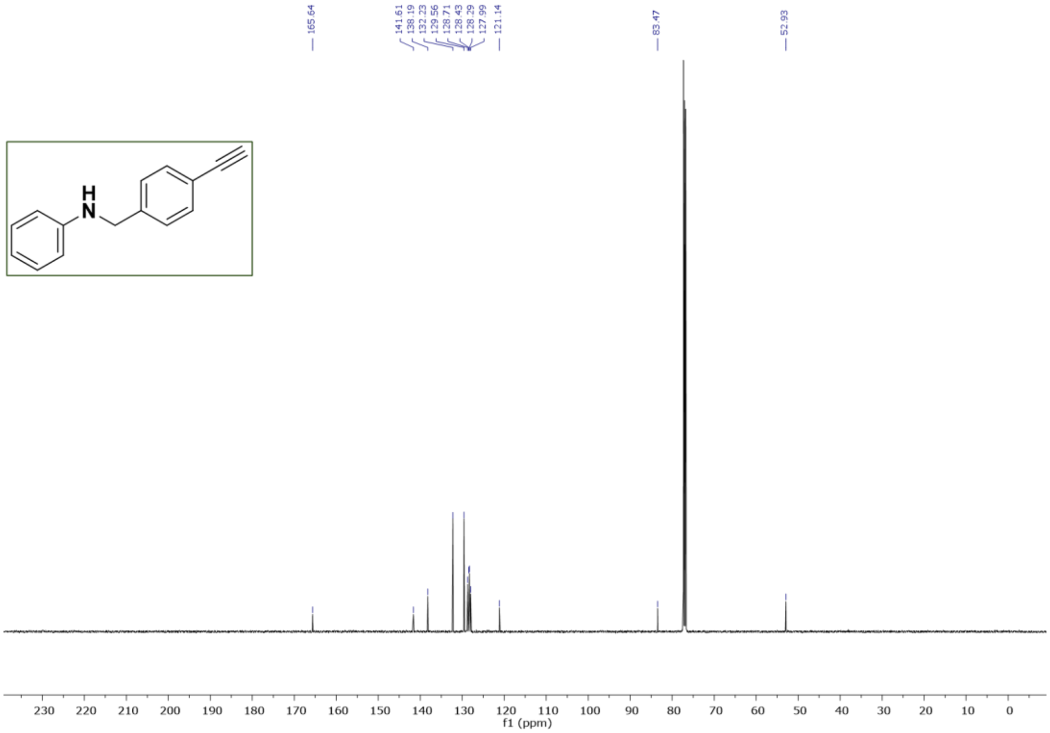

**Figure S2.** [13]C NMR of ***N*-(4-ethynylbenzyl)aniline (S01)** in $CDCl_3$

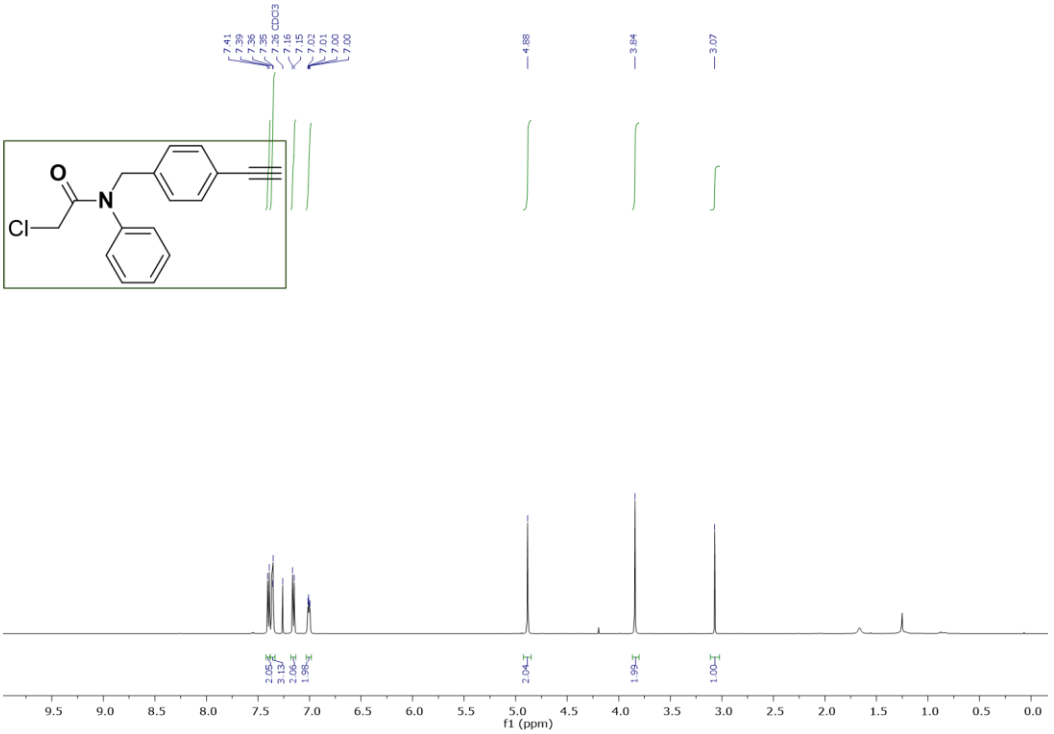

**Figure S3.**[1]H NMR **of 2-chloro-*N*-(4-ethynylbenzyl)-*N*-phenylacetamide (1)** in $CDCl_3$

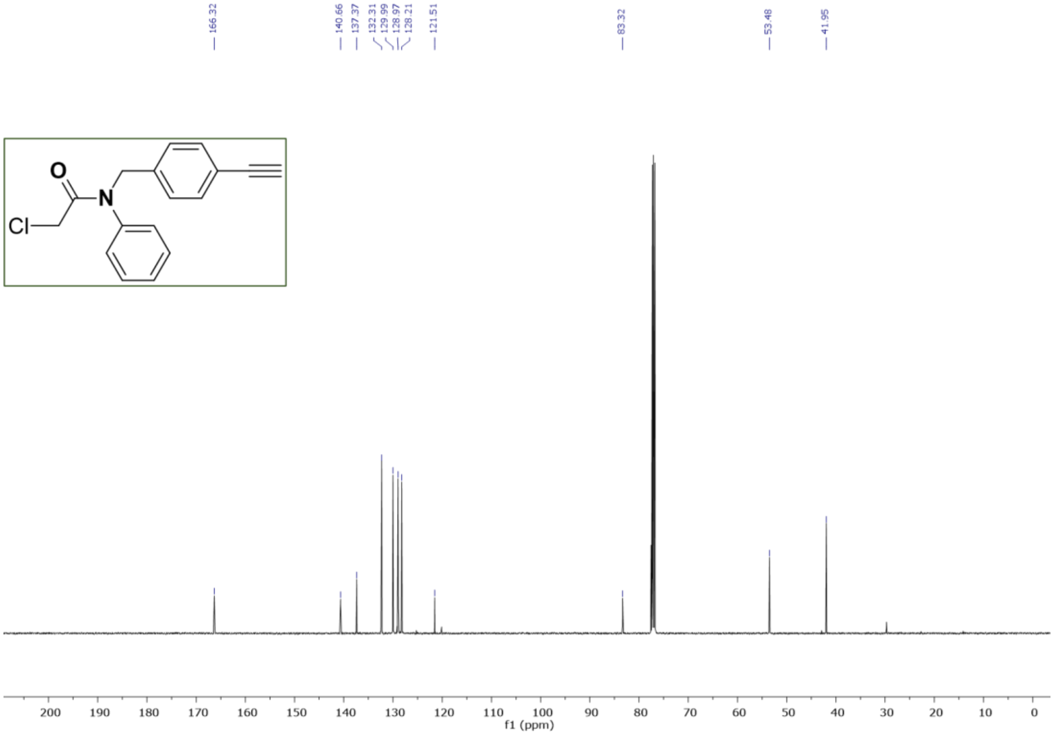

**Figure S4.** $^{13}$C NMR **of 2-chloro-N-(4-ethynylbenzyl)-N-phenylacrylamide (1)** in $CDCl_3$

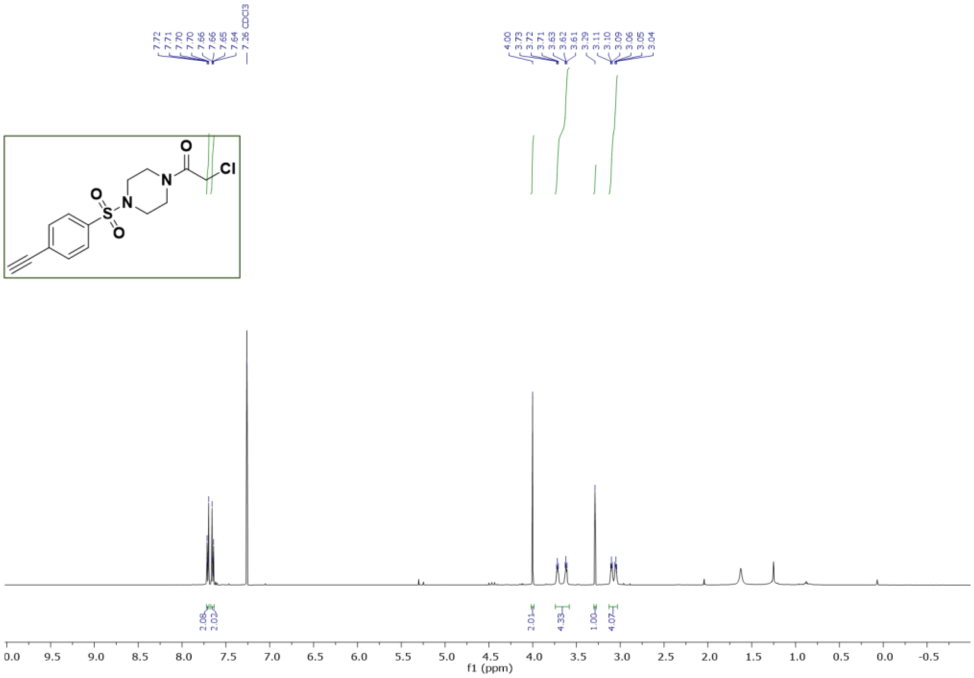

**Figure S5.**[1]H NMR **of 2-chloro-1-(4-((4-ethynylphenyl)sulfonyl)piperazin-1-yl)ethan-1-on (2)** in $CDCl_3$

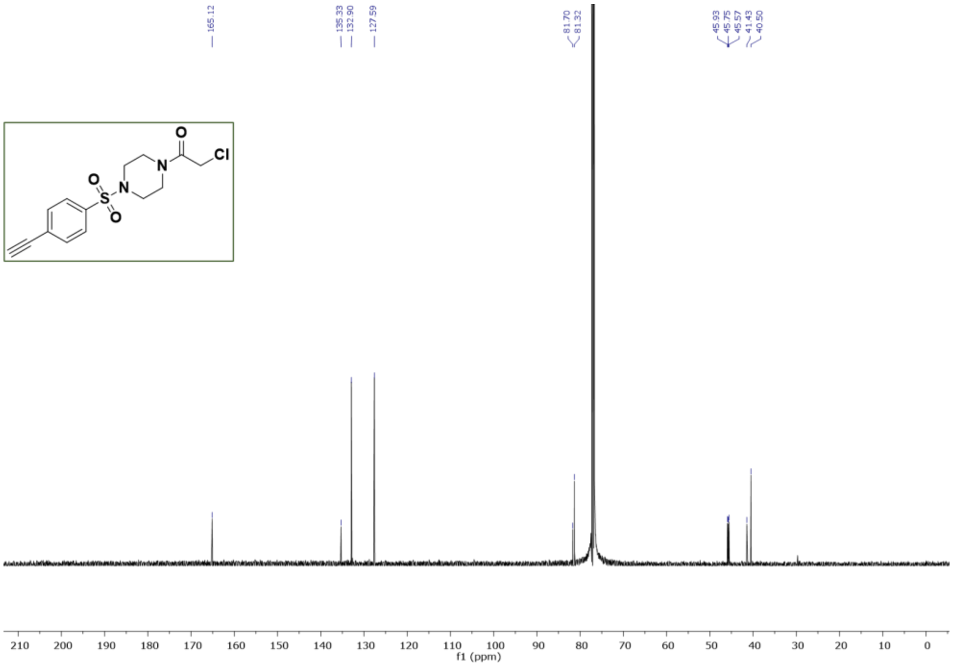

**Figure S6.**[13]C NMR **of 2-chloro-1-(4-((4-ethynylphenyl)sulfonyl)piperazin-1-yl)ethan-1-on (2)** in $CDCl_3$

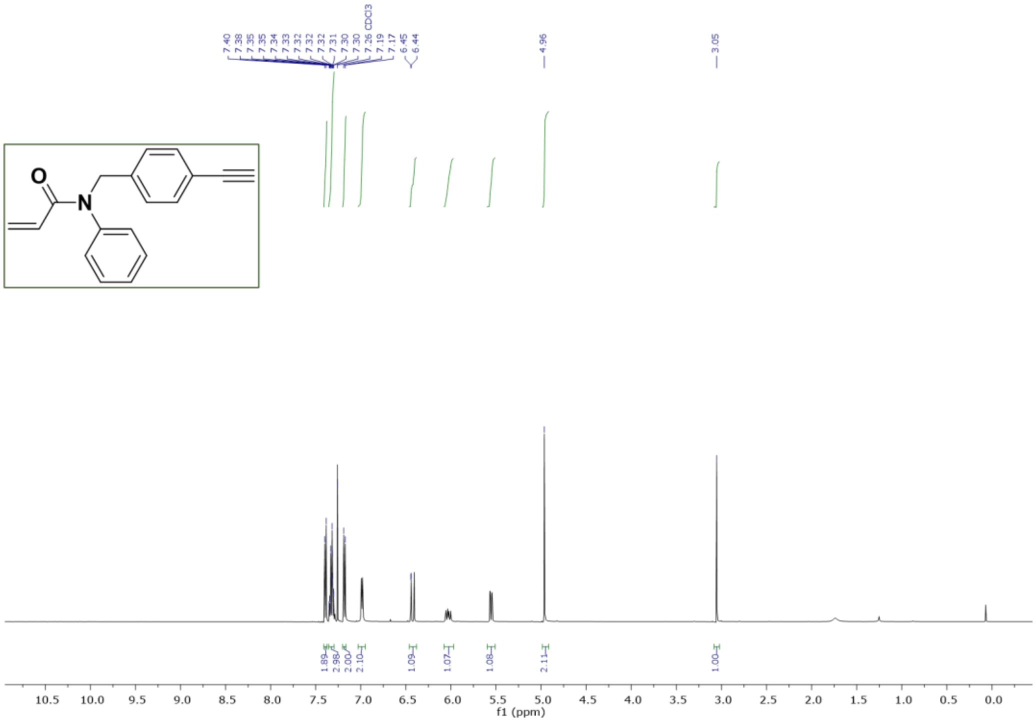

**Figure S7.**$^{1}$H NMR **of N-(4-ethynylbenzyl)-N-phenylacrylamide (3)** in $CDCl_3$

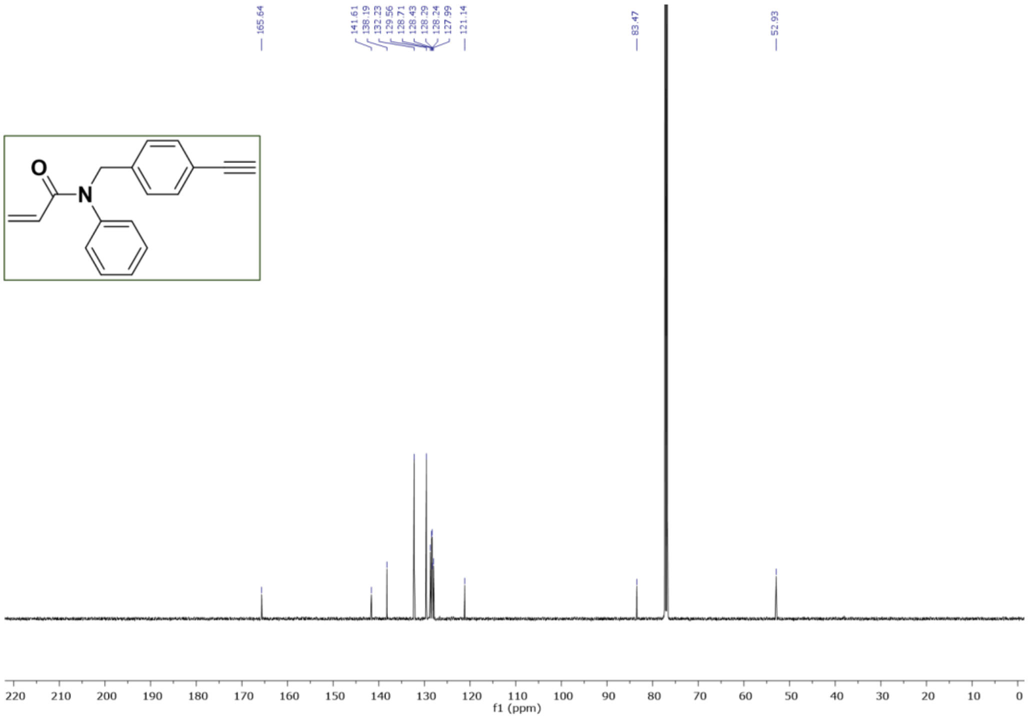

**Figure S8.**$^{13}$C NMR **of N2-chloro-N-(4-ethynylbenzyl)-N-phenylacetamide (3)** in $CDCl_3$

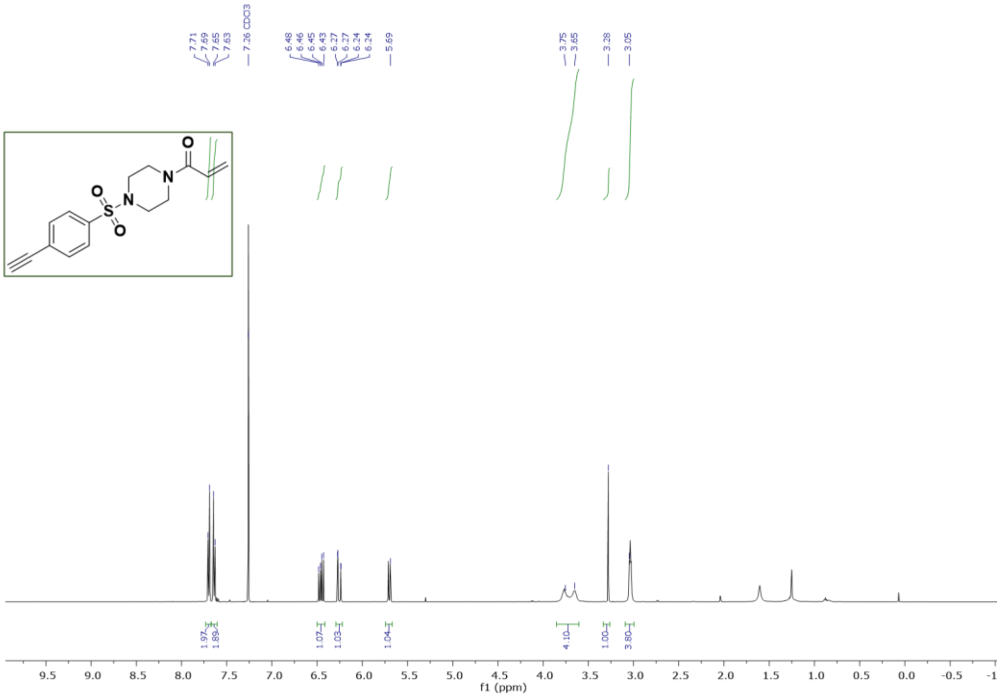

**Figure S9.**[1]H NMR **of 1-(4-((4-ethynylphenyl)sulfonyl)piperazin-1-yl)prop-2-en-1-one (4)** in $CDCl_3$

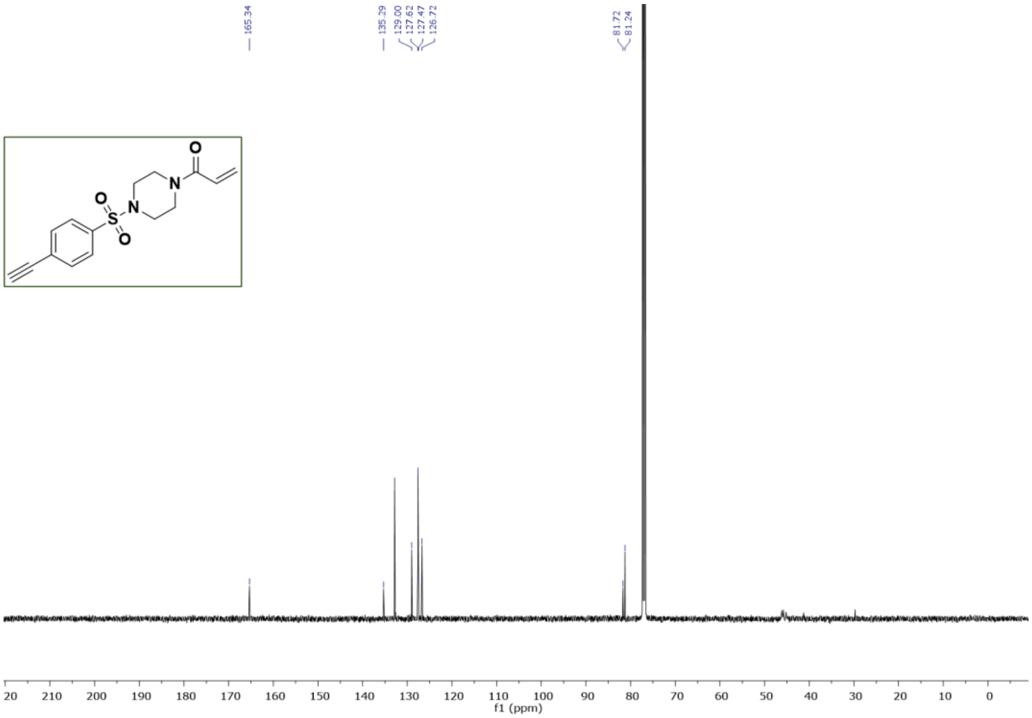

**Figure S10.**[13]C NMR **of 1-(4-((4-ethynylphenyl)sulfonyl)piperazin-1-yl)prop-2-en-1-one (4)** in $CDCl_3$

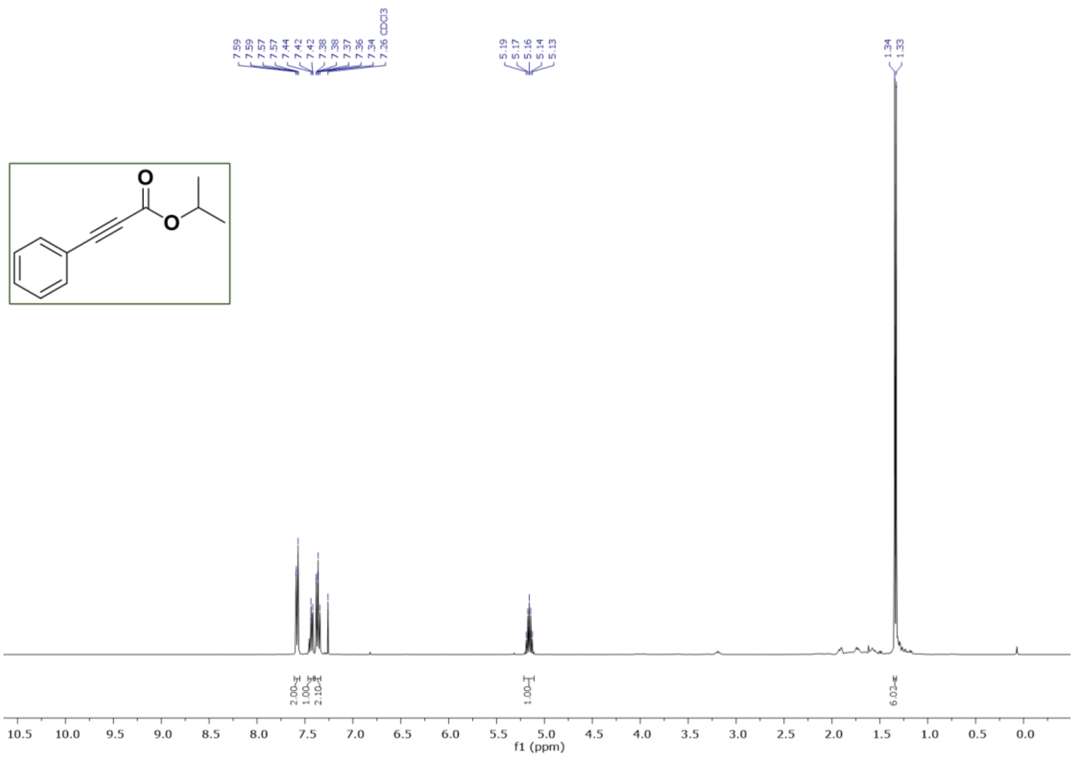

**Figure S11.** $^{1}$H NMR **of isopropyl 3-phenylpropiolate (5)** in $CDCl_3$

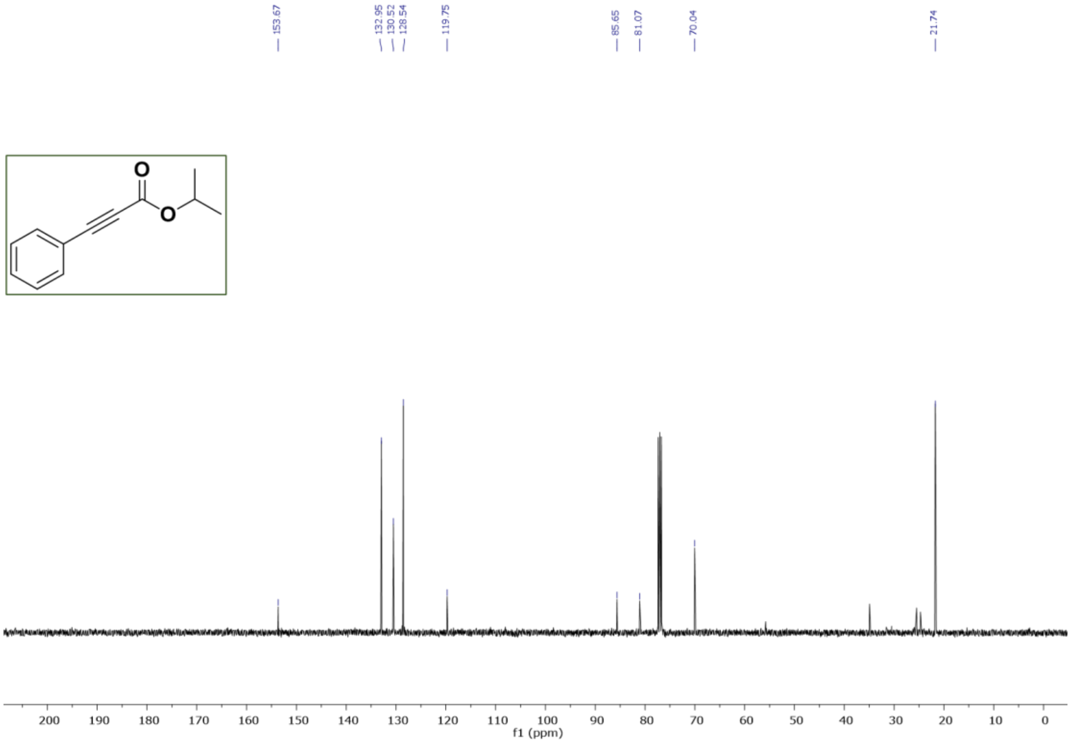

**Figure S12.** $^{13}$C NMR **of isopropyl 3-phenylpropiolate (5)** in $CDCl_3$

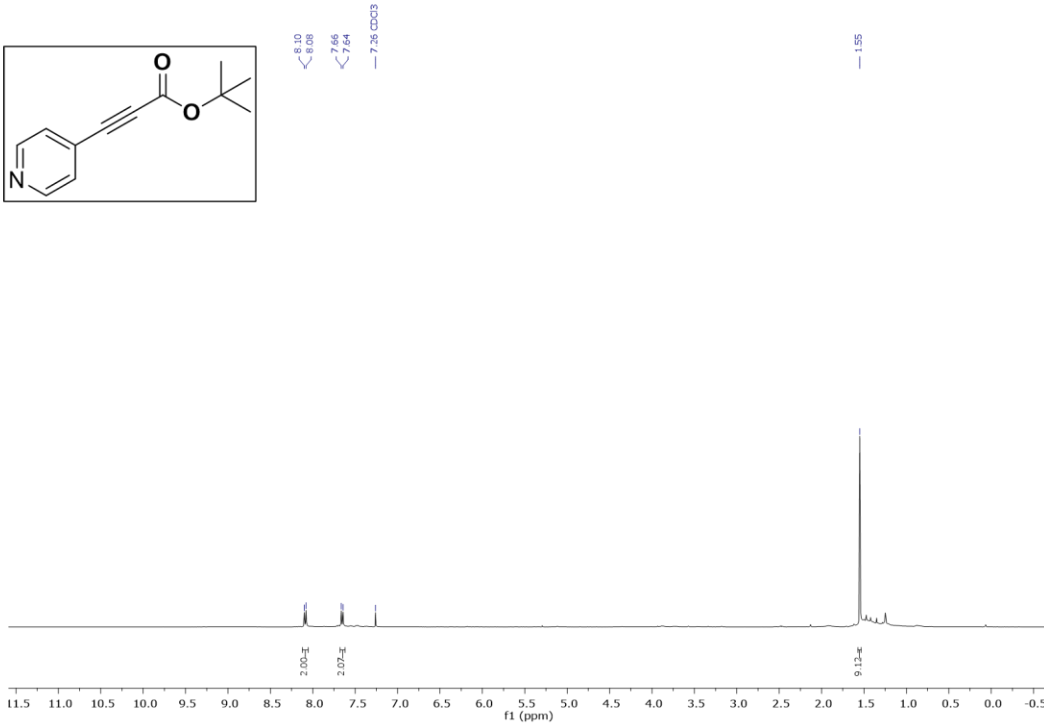

**Figure S13.** $^{1}$H NMR of **tert-butyl 3-(pyridin-4-yl)propiolate (6)** in $CDCl_3$

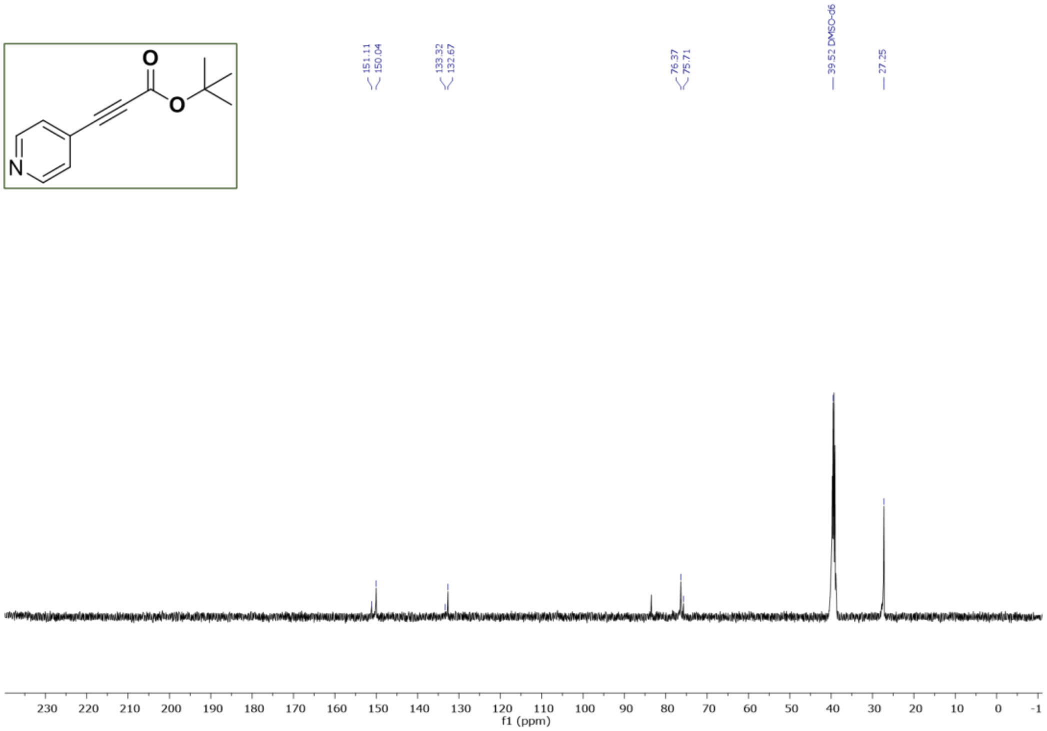

**Figure S14.** $^{13}$C NMR of **tert-butyl 3-(pyridin-4-yl)propiolate (6)** in DMSO-$d_6$

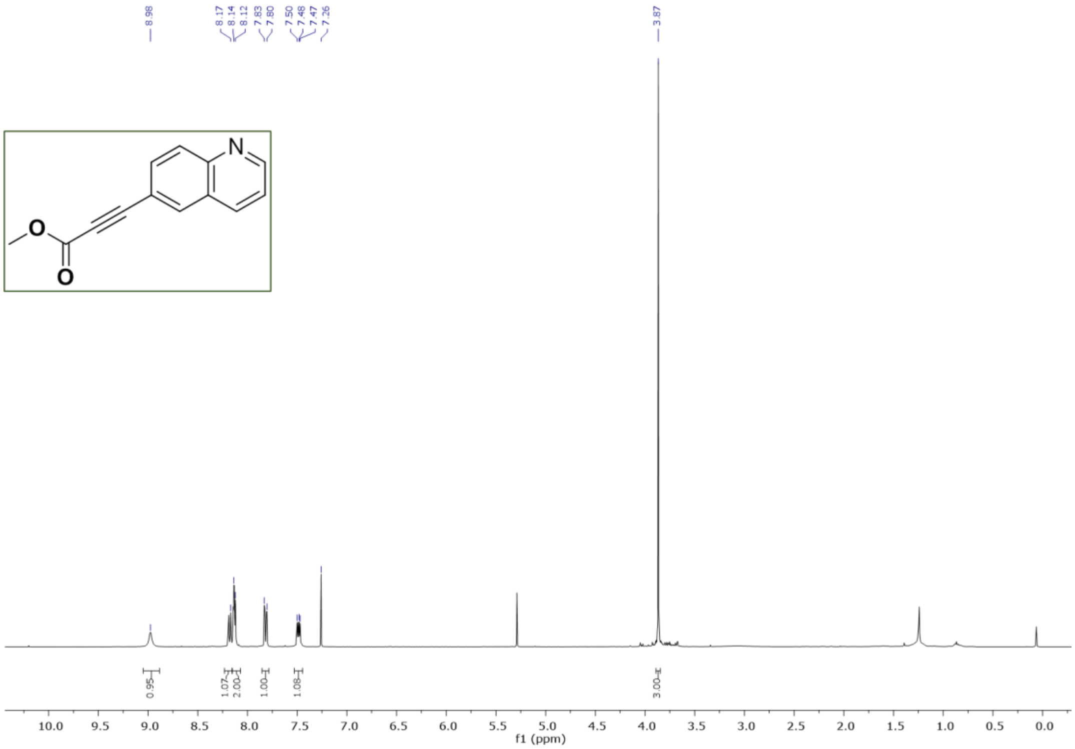

**Figure S15.** $^{1}$H NMR of **Methyl 3-(quinolin-6-yl)propiolate (7)** in $CDCl_3$

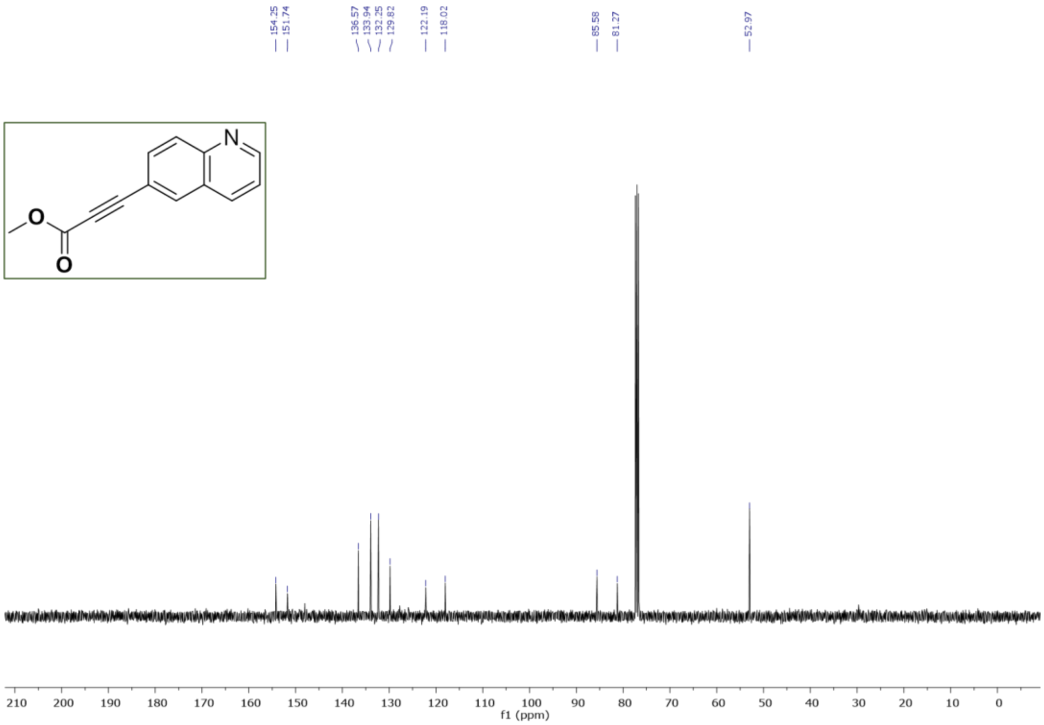

**Figure S16.** $^{13}$C NMR of **Methyl 3-(quinolin-6-yl)propiolate (7)** in $CDCl_3$

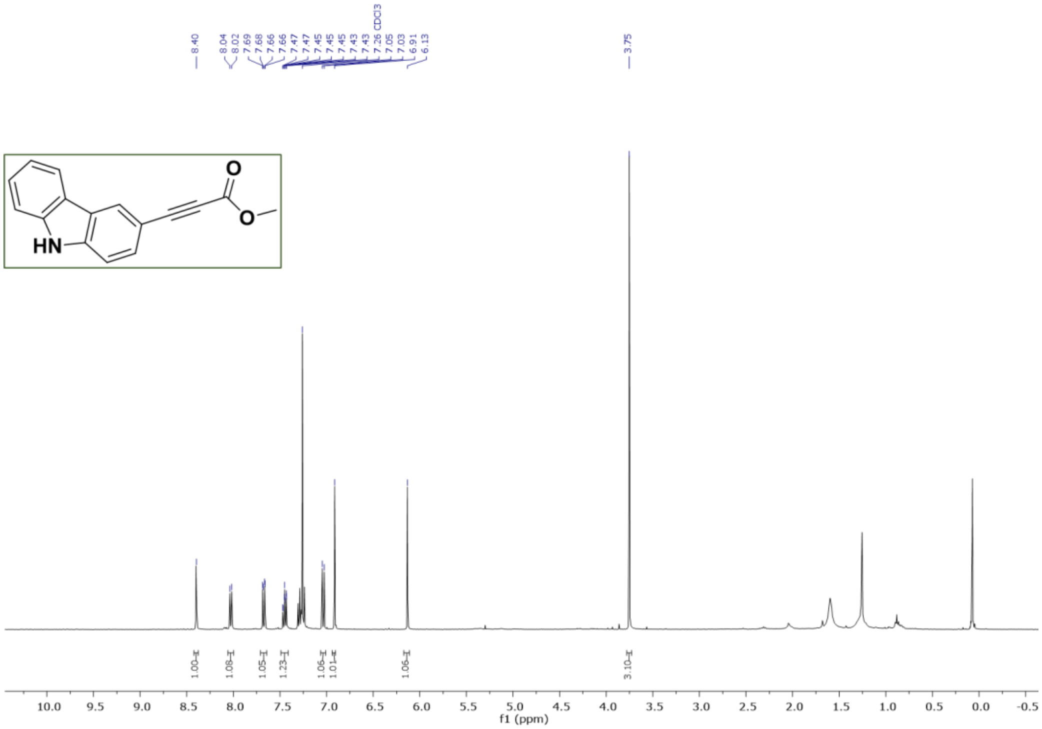

**Figure S17.** $^{1}$H NMR of **methyl 3-(9H-carbazol-3-yl)propiolate (8)** in $CDCl_3$

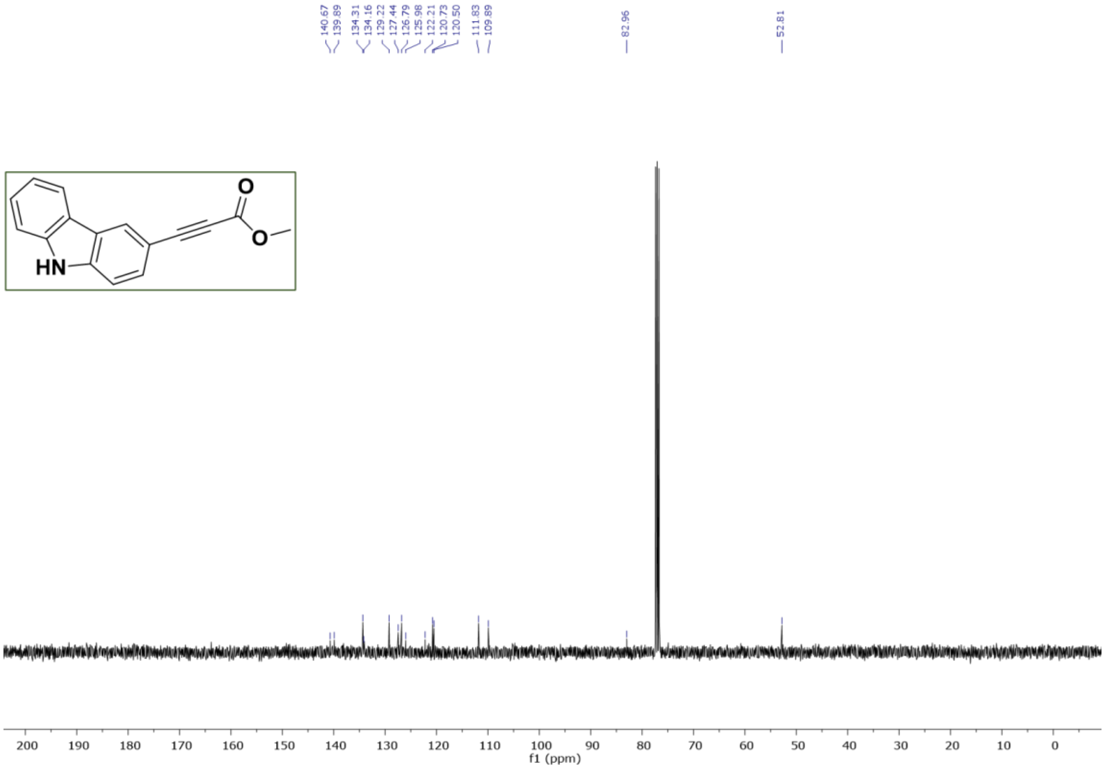

**Figure S18.** $^{13}$C NMR of **methyl 3-(9H-carbazol-3-yl)propiolate** (8) in $CDCl_3$

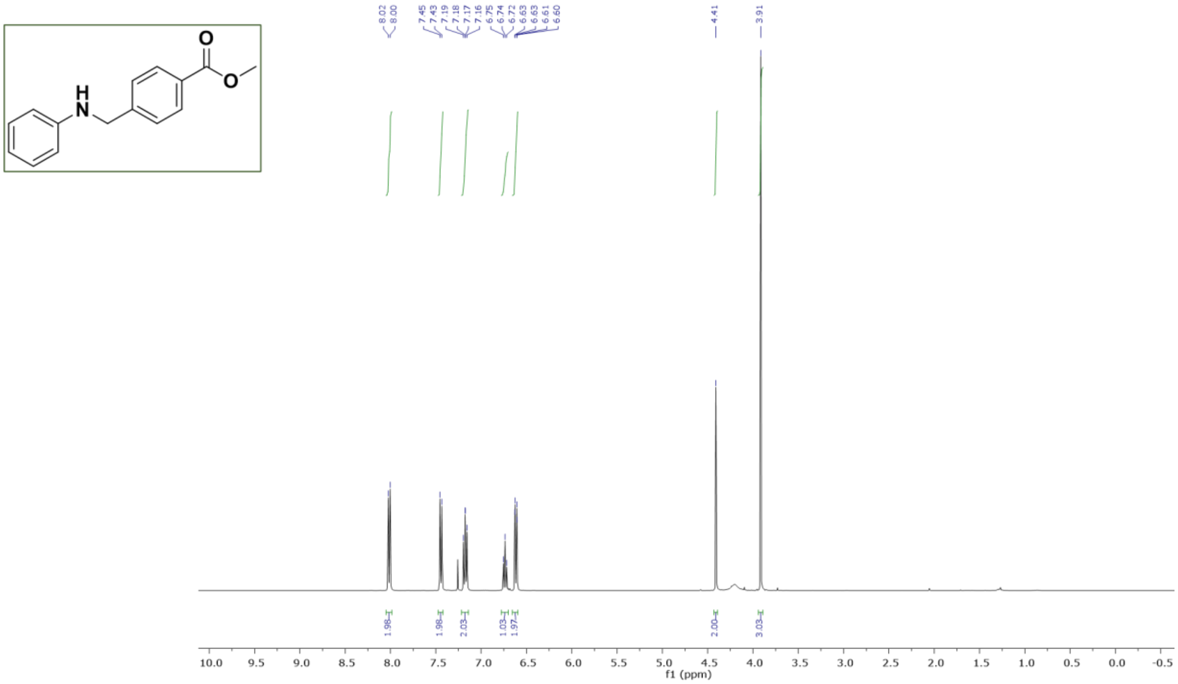

**Figure S19.**$^{1}$H NMR **of methyl 4-((phenylamino)methyl)benzoate (S03)** in $CDCl_3$

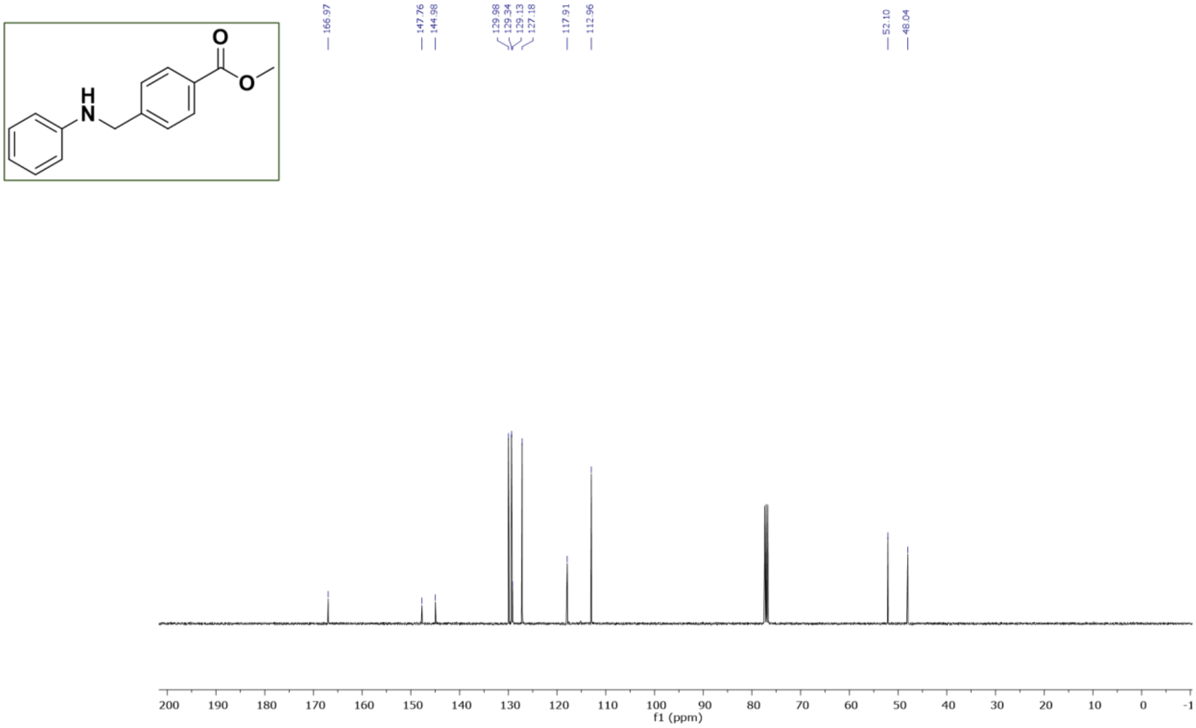

**Figure 20.**$^{1}$H NMR **of methyl 4-((phenylamino)methyl)benzoate (S03)** in $CDCl_3$

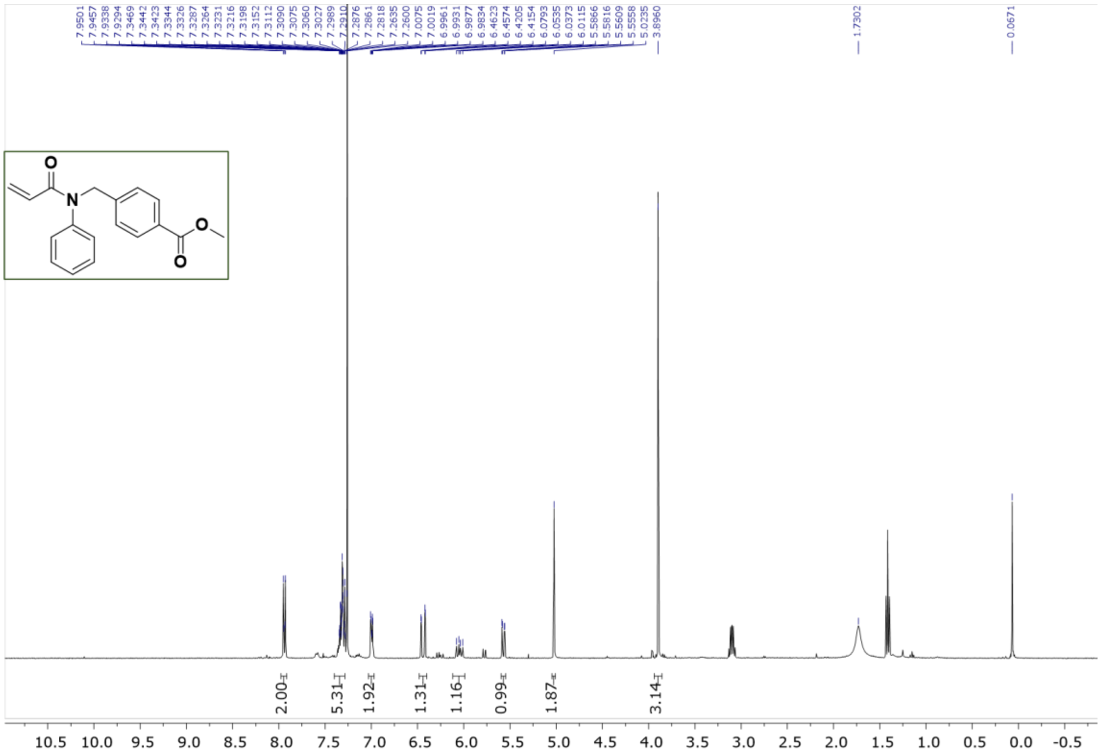

**Figure S21.** $^{1}$H NMR **of methyl 4-((N-phenylacrylamido)methyl)benzoate (9)** in $CDCl_3$

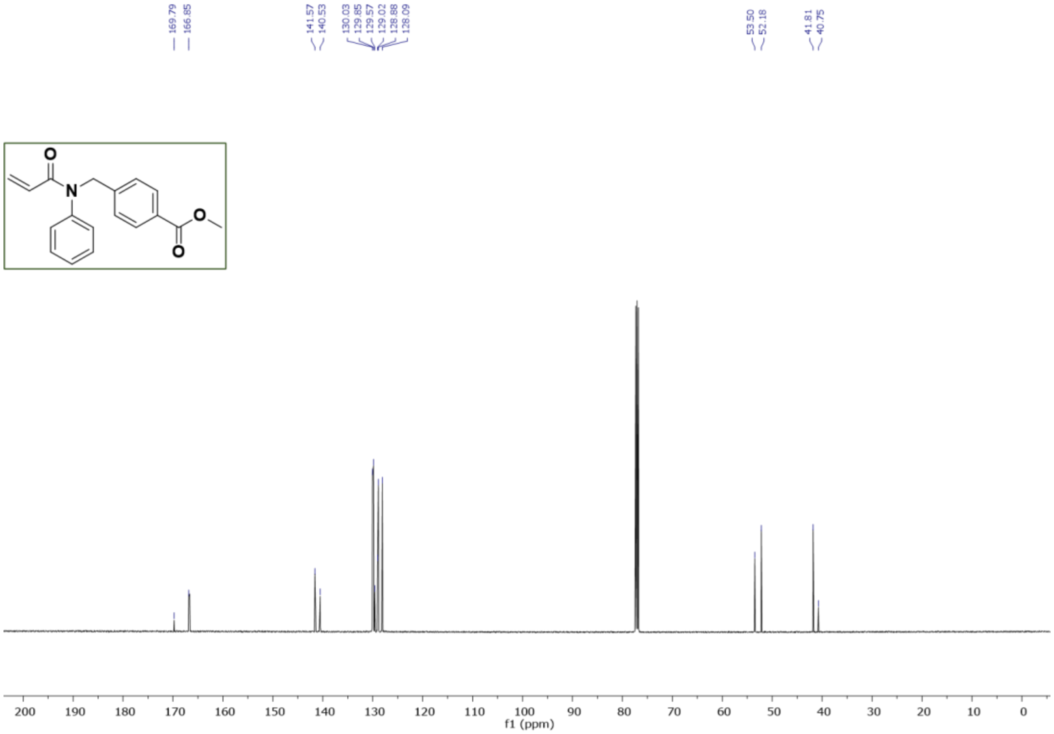

**Figure S22.** $^{13}$C NMR **of methyl 4-((N-phenylacrylamido)methyl)benzoate (9)** in $CDCl_3$

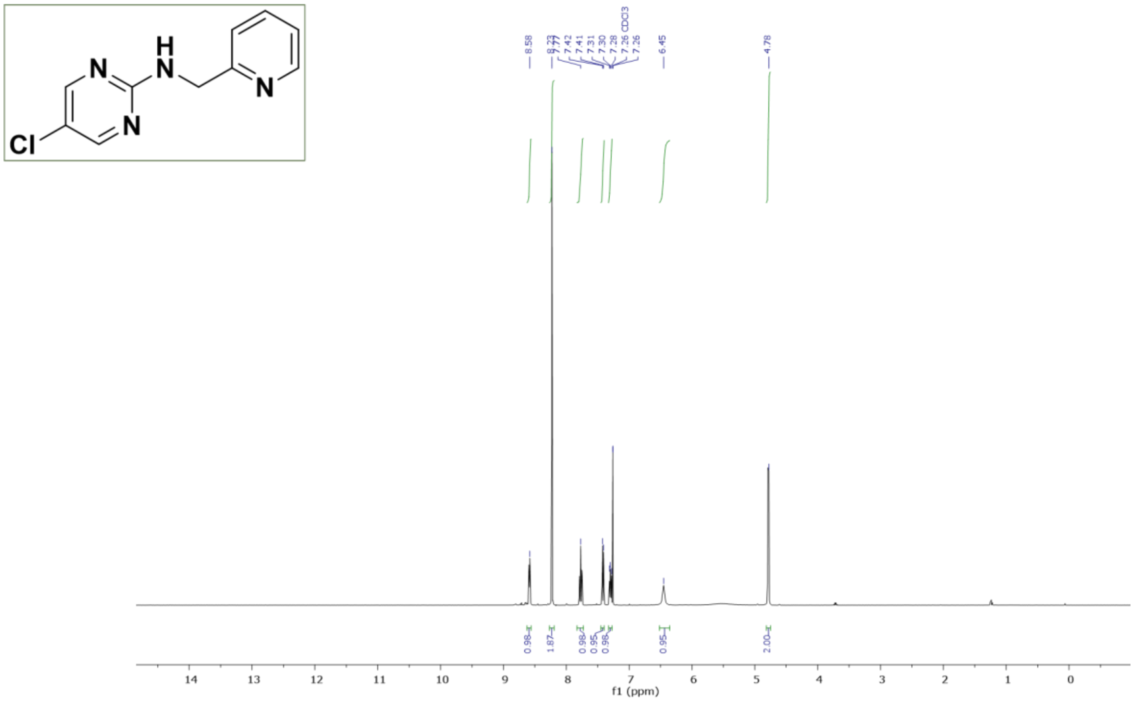

**Figure S23.** $^{1}$H NMR of **5-chloro-N-(pyridin-2-ylmethyl)pyrimidin-2-amine (S04)** in $CDCl_3$

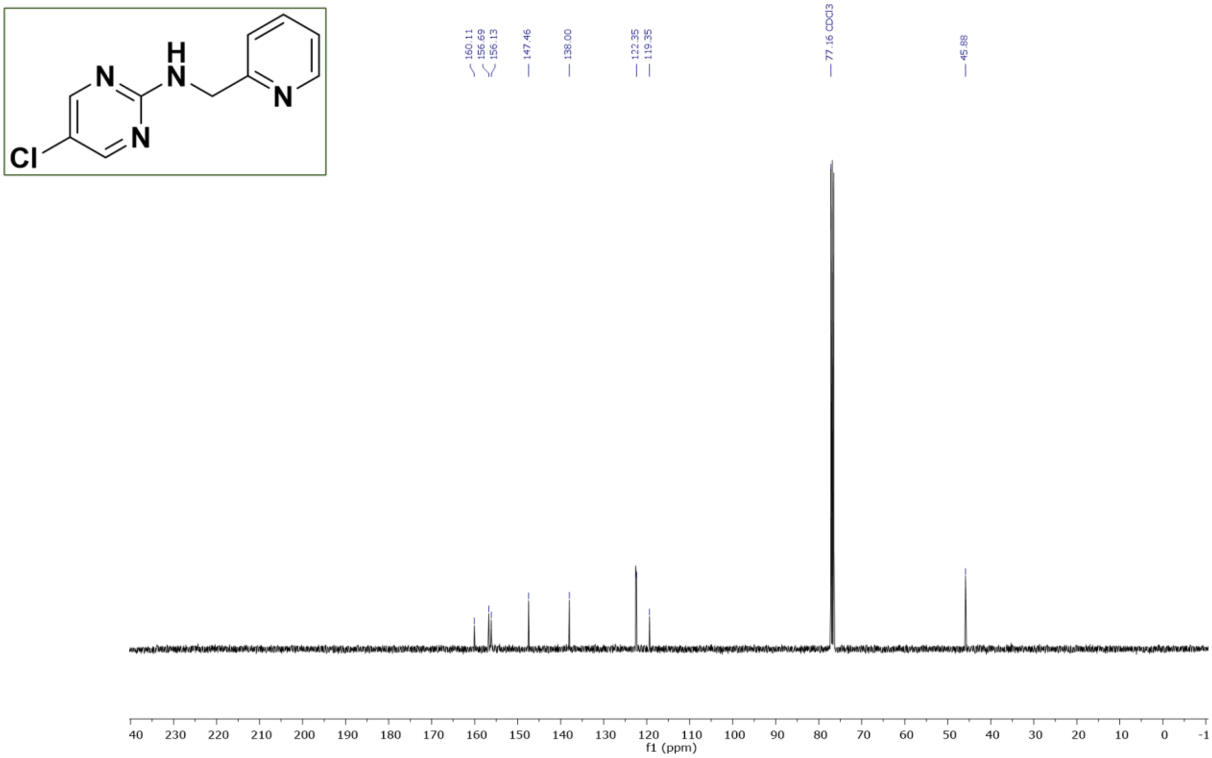

**Figure S24.** $^{13}$C NMR of **5-chloro-N-(pyridin-2-ylmethyl)pyrimidin-2-amine (S04)** in $CDCl_3$

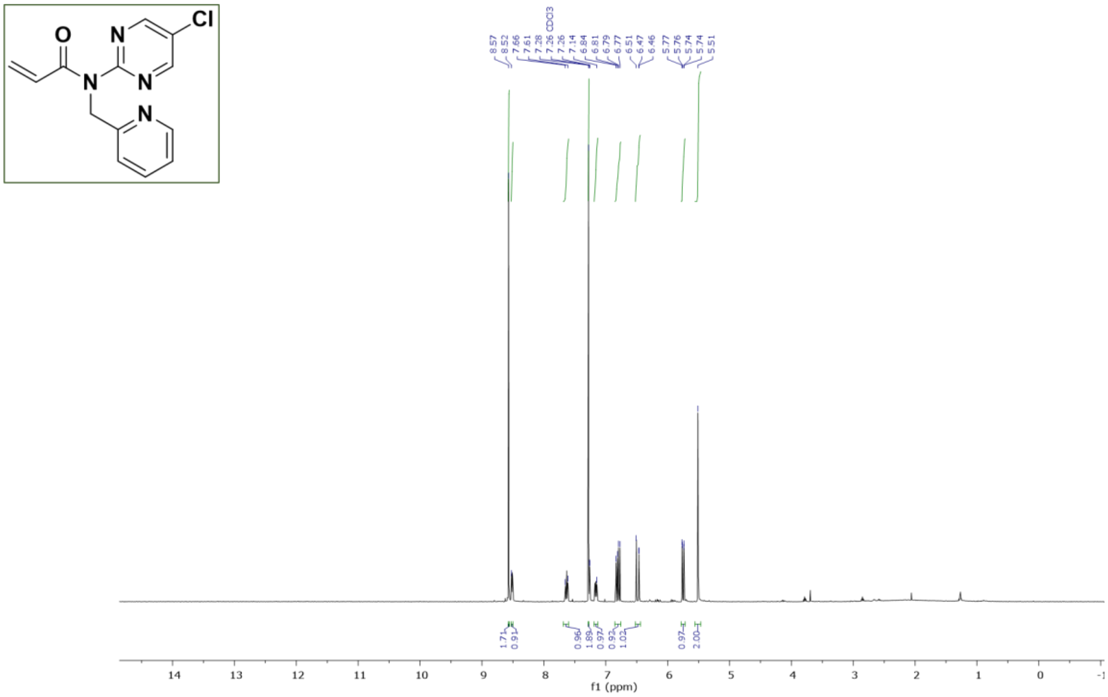

**Figure S25.** $^{1}H$ NMR of **N-(5-chloropyrimidin-2-yl)-N-(pyridin-2-ylmethyl)acrylamide (10)** in $CDCl_3$

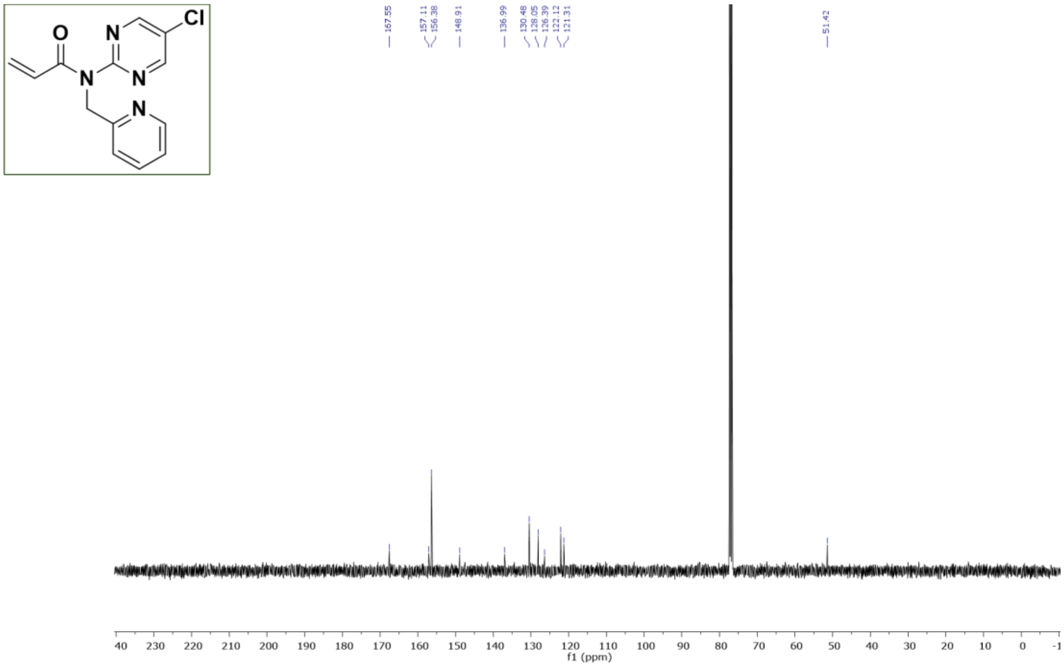

**Figure S26.** $^{13}C$ NMR of **N-(5-chloropyrimidin-2-yl)-N-(pyridin-2-ylmethyl)acrylamide (10)** in $CDCl_3$

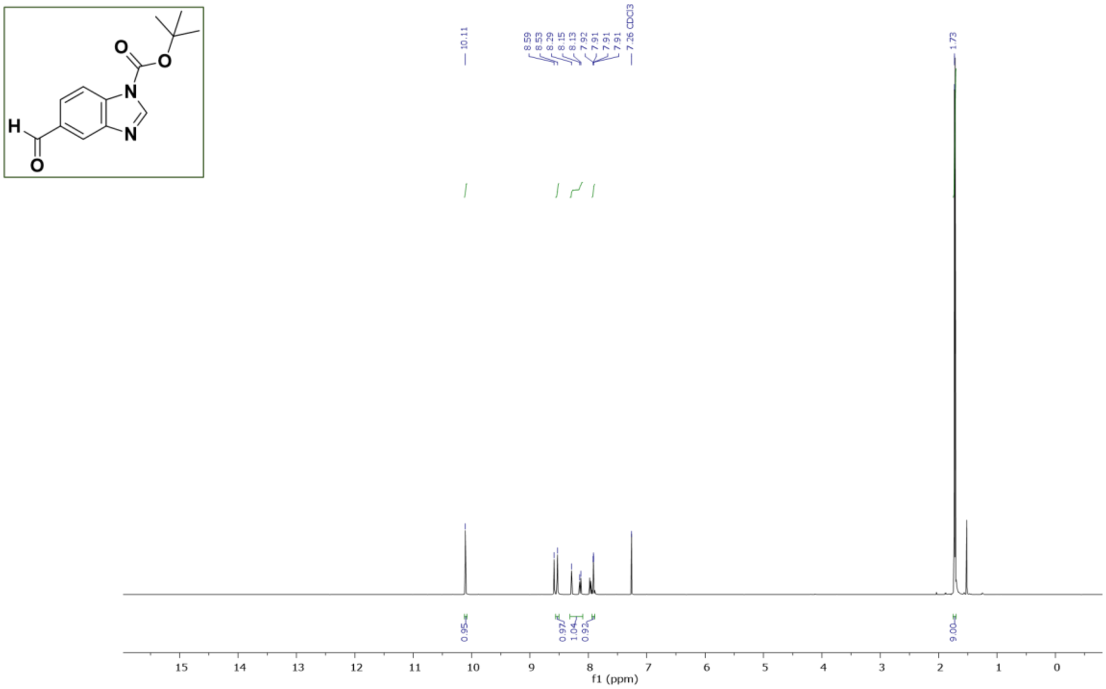

**Figure S27.**[1]H NMR of **tert-butyl 5-formyl-1H-benzo[d]imidazole-1-carboxylate (S05)** in $CDCl_3$

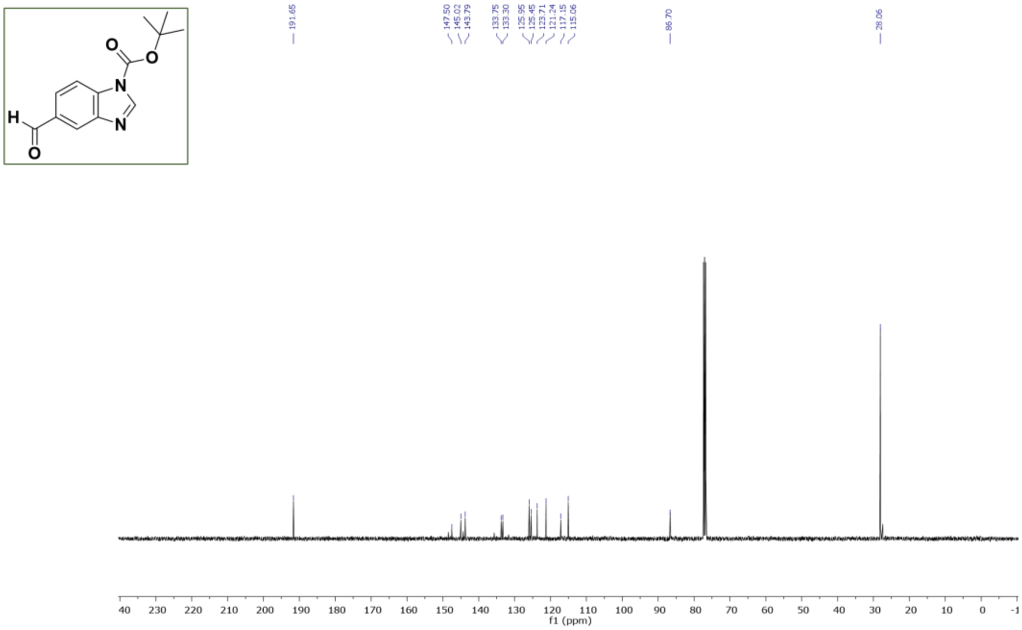

**Figure S28.** $^{13}C$ NMR of **tert-butyl 5-formyl-1H-benzo[d]imidazole-1-carboxylate (S05)** in $CDCl_3$

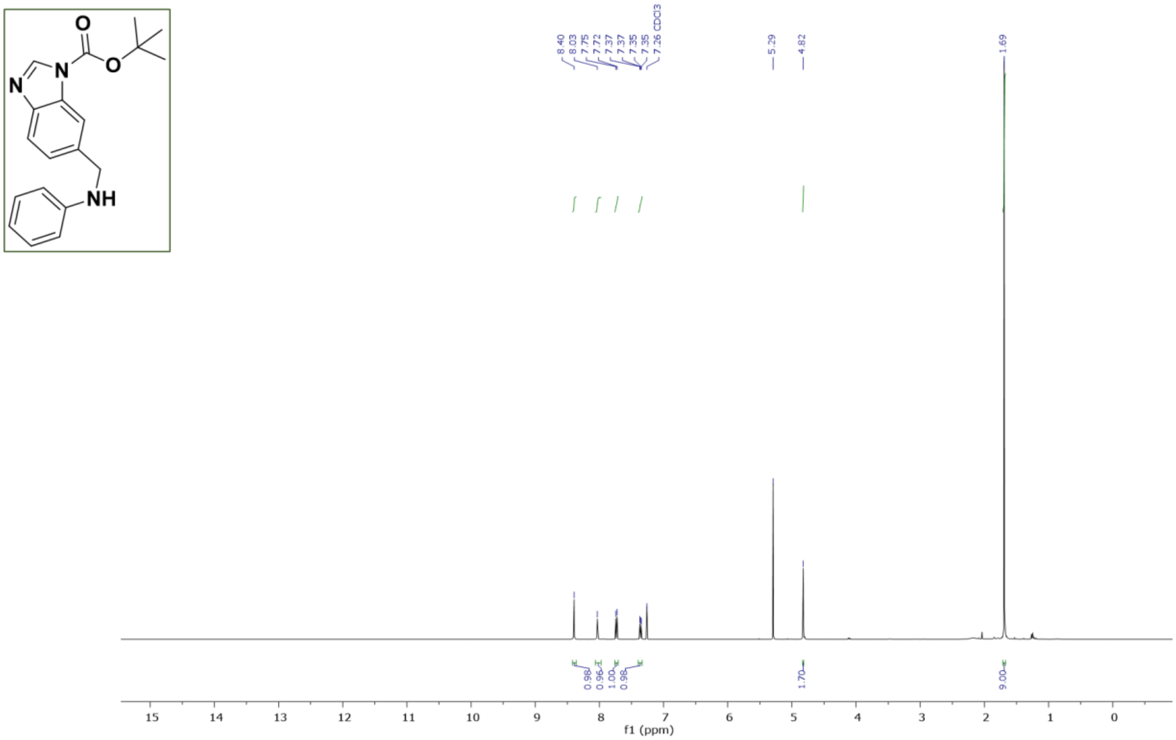

**Figure S29.** $^{1}$H NMR of **tert-butyl 5-((N-phenylacrylamido)methyl)-1H-benzo[d]imidazole-1-carboxylate (S06)** in $CDCl_3$

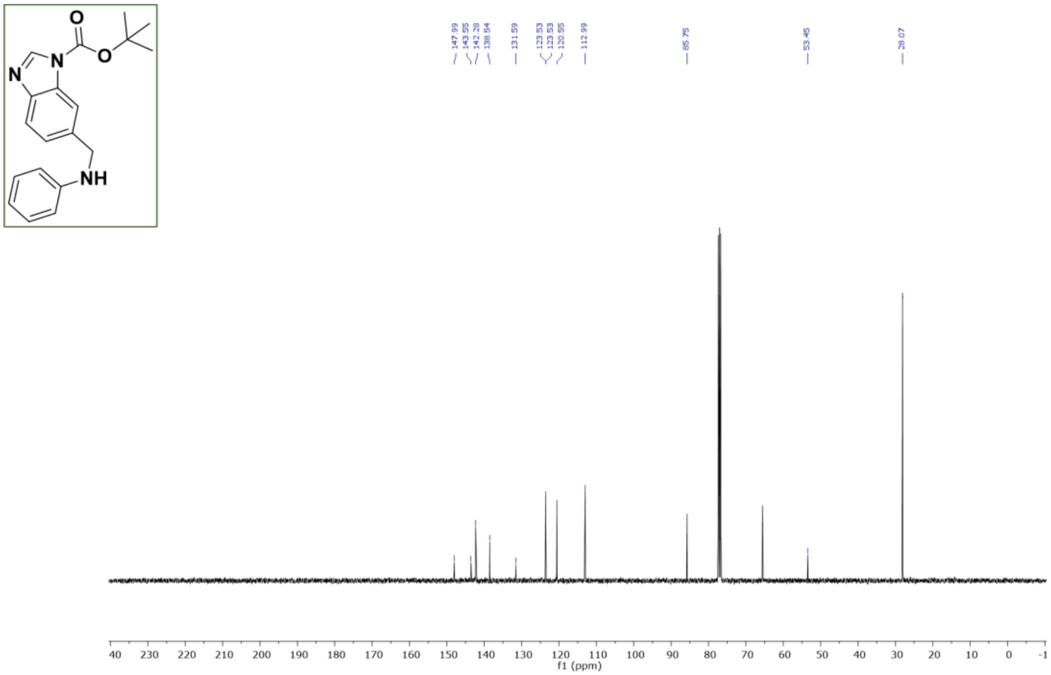

**Figure S30.** $^{13}$C NMR of **tert-butyl 5-((N-phenylacrylamido)methyl)-1H-benzo[d]imidazole-1-carboxylate (S06)** in $CDCl_3$

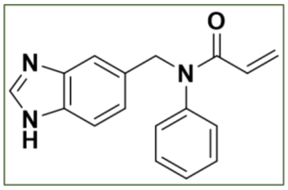

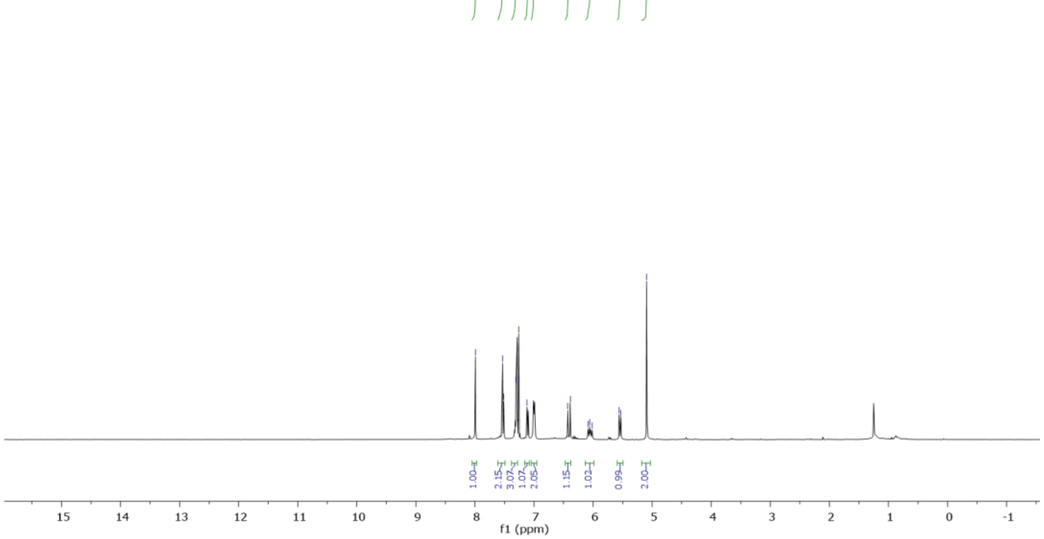

**Figure S31.** $^{1}$H NMR of **N-((1H-benzo[d]imidazol-5-yl)methyl)-N-phenylacrylamide (11)** in $CDCl_3$

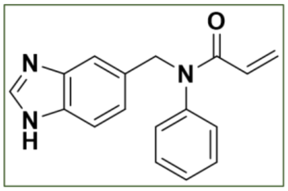

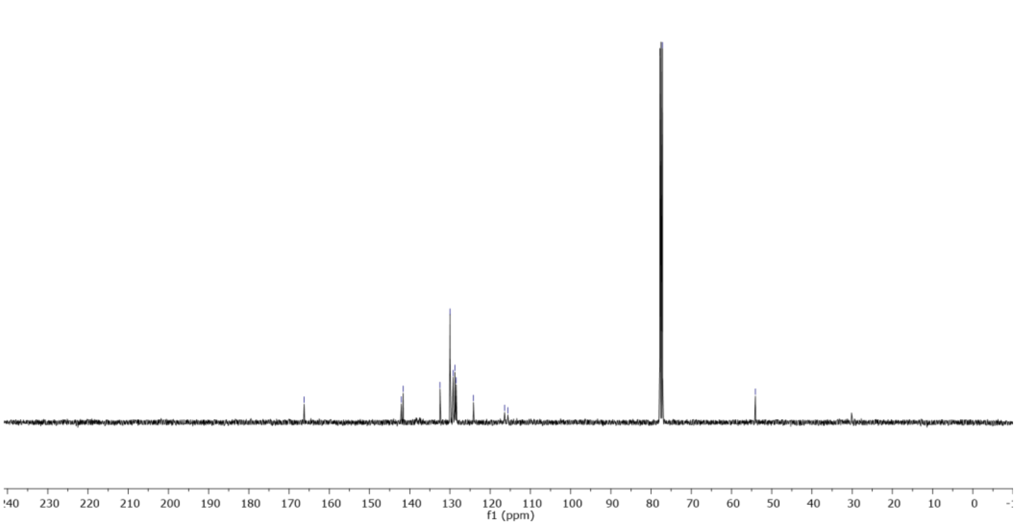

**Figure S32.** $^{13}$C NMR of **N-((1H-benzo[d]imidazol-5-yl)methyl)-N-phenylacrylamide (11)** in $CDCl_3$

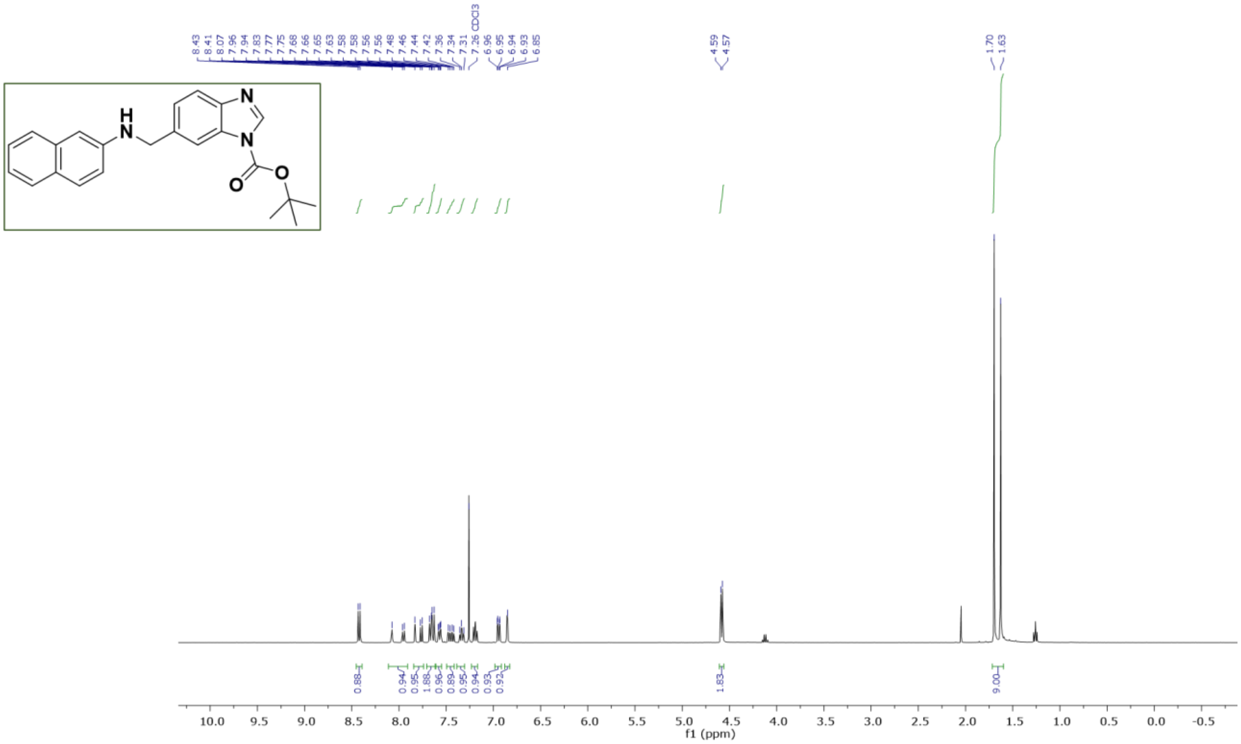

**Figure S33.** $^{1}$H NMR of **tert-butyl 6-((naphthalen-2-ylamino)methyl)-1H-benzo[d]imidazole-1-carboxylate (S08)** in $CDCl_3$

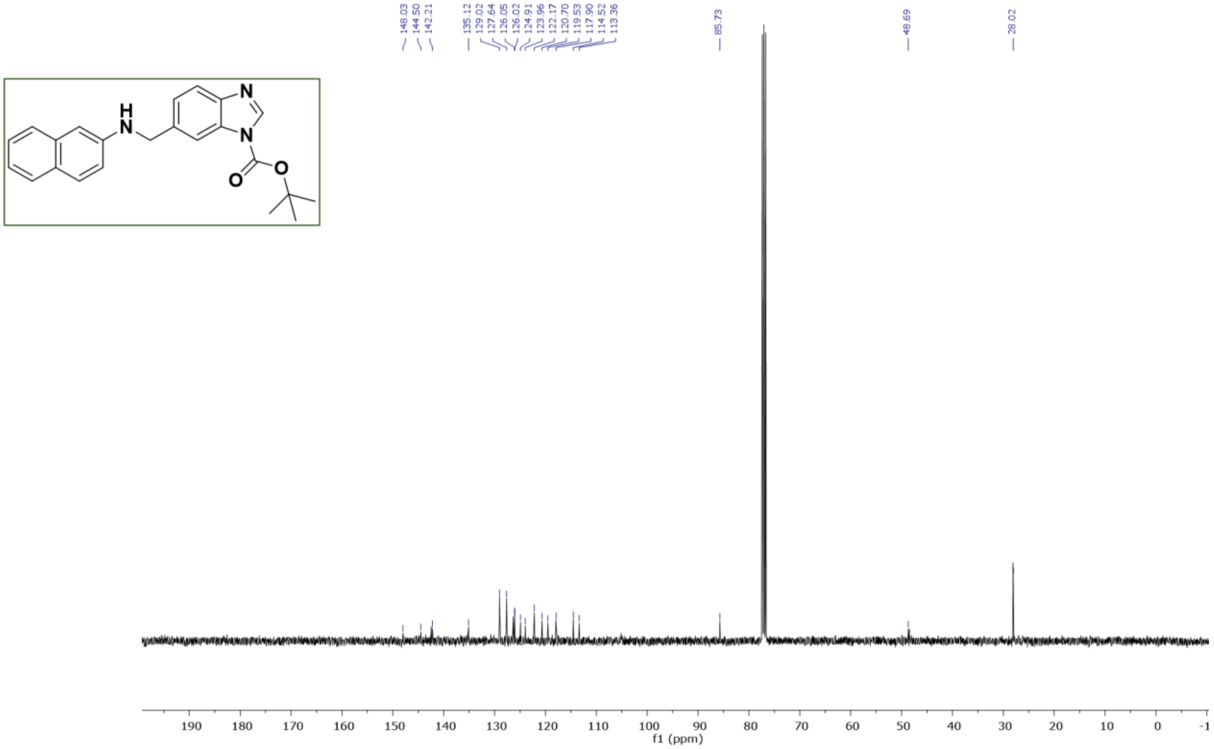

**Figure S34.** $^{1}$H NMR of **tert-butyl 6-((naphthalen-2-ylamino)methyl)-1H-benzo[d]imidazole-1-carboxylate (S08)** in $CDCl_3$

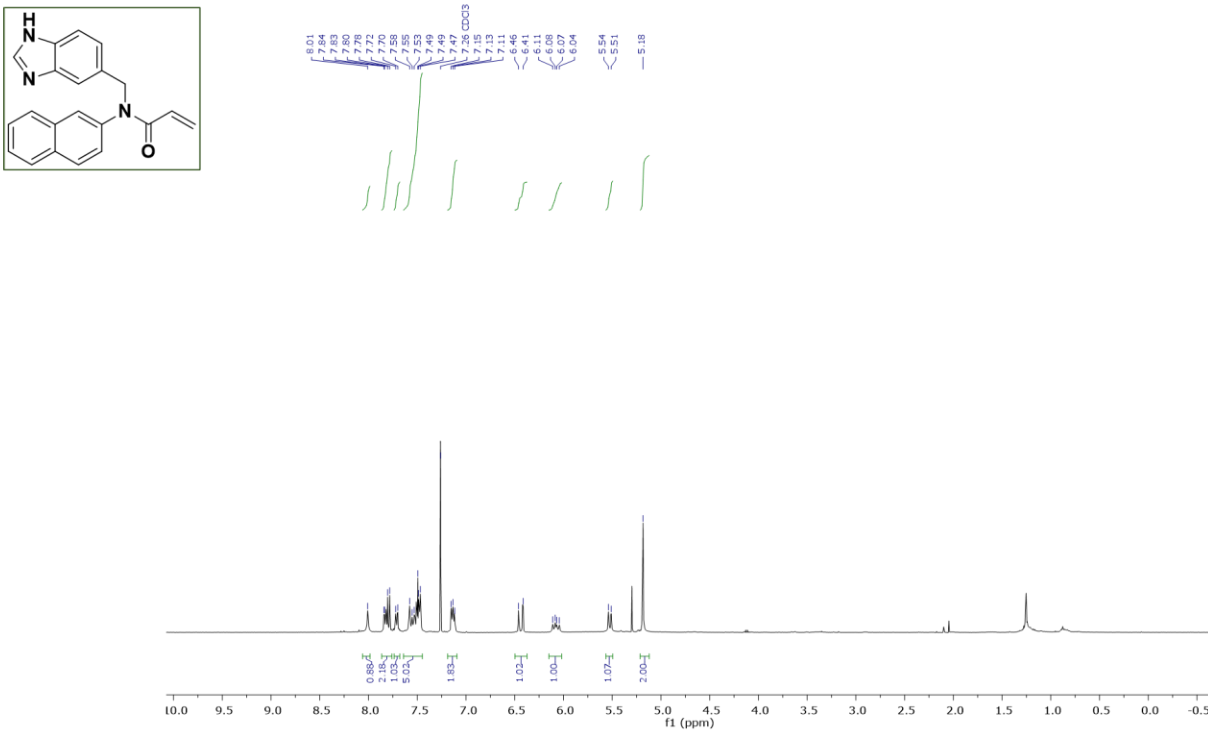

**Figure S35.** $^{1}H$ NMR of N-((1H-benzo[d]imidazol-6-yl)methyl)-N-(naphthalen-2-yl)acrylamide (12) in $CDCl_3$

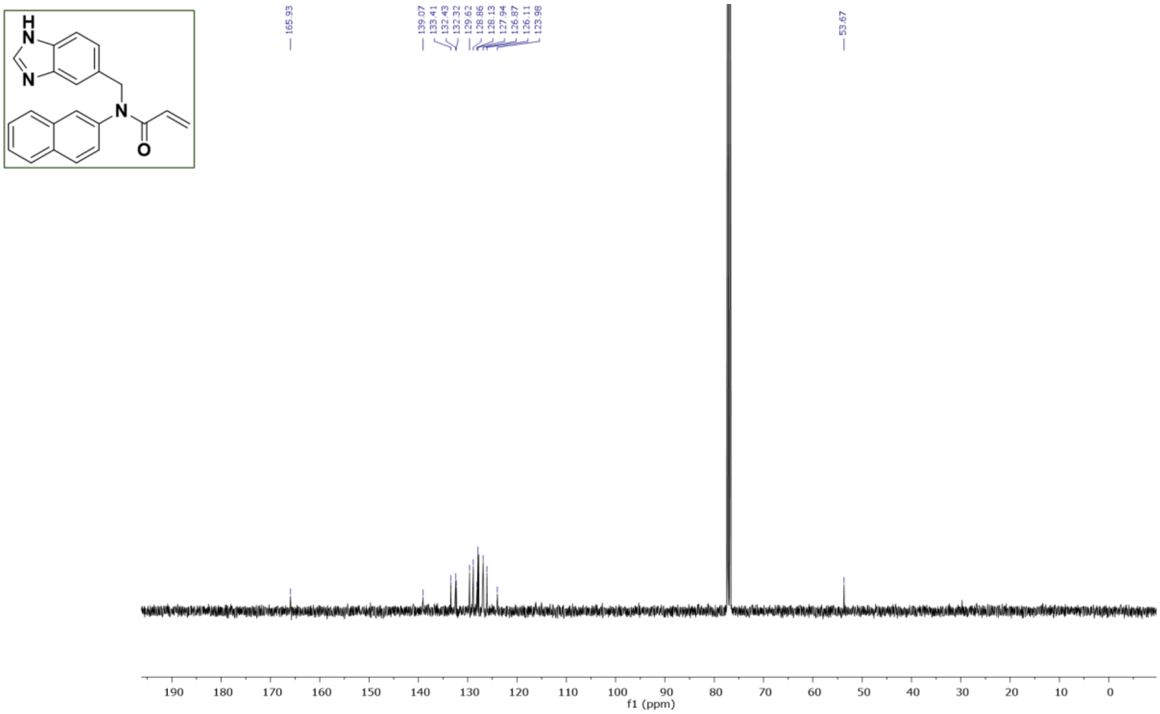

**Figure S36.** $^{13}C$ NMR of N-((1H-benzo[d]imidazol-6-yl)methyl)-N-(naphthalen-2-yl)acrylamide (12) in $CDCl_3$

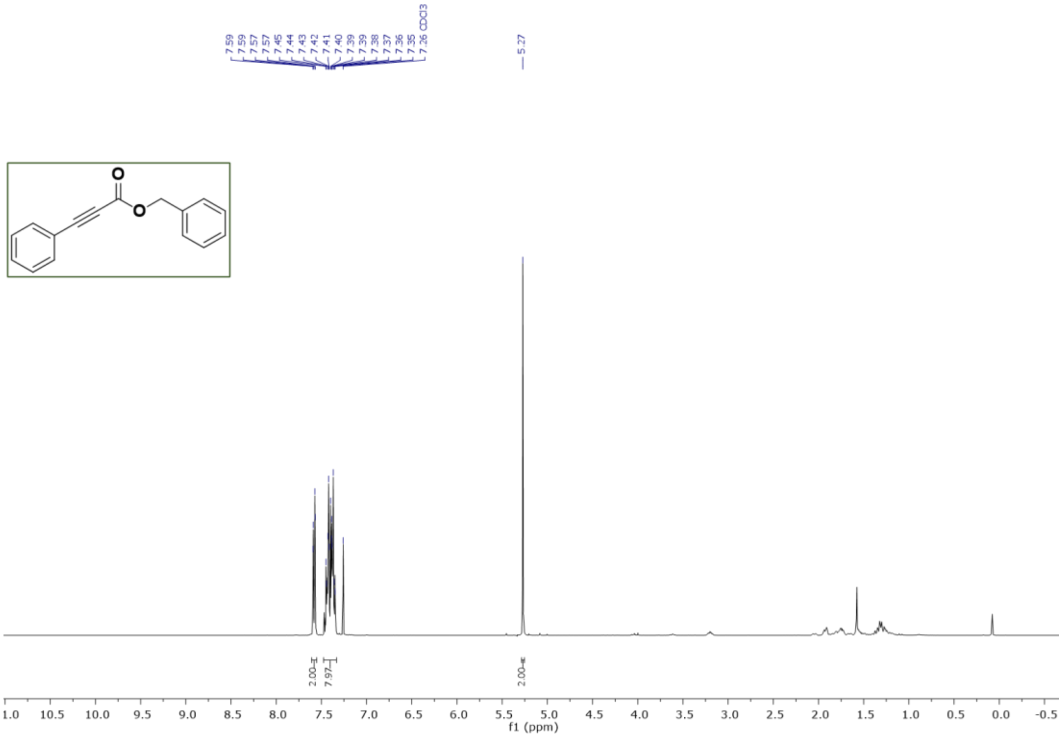

**Figure S37.** $^{1}$H NMR **of benzyl 3-phenylpropiolate (13)** in $CDCl_3$

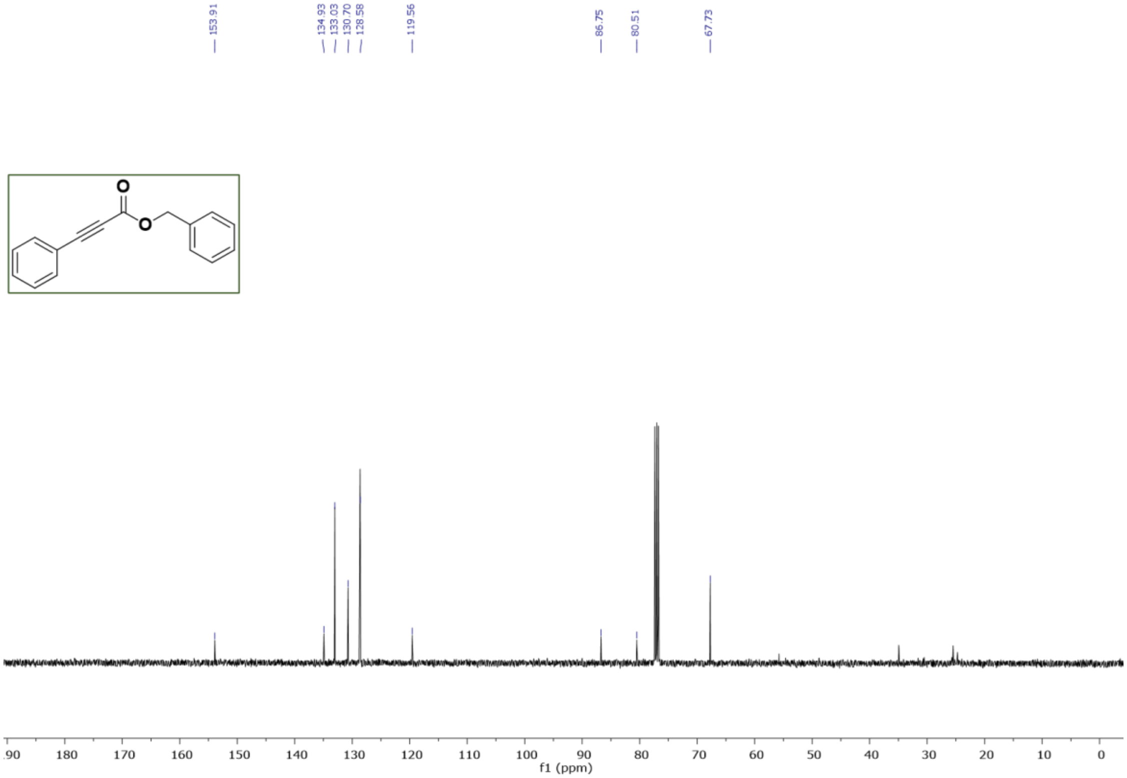

**Figure S38.** $^{13}$C NMR **of benzyl 3-phenylpropiolate (13)** in $CDCl_3$

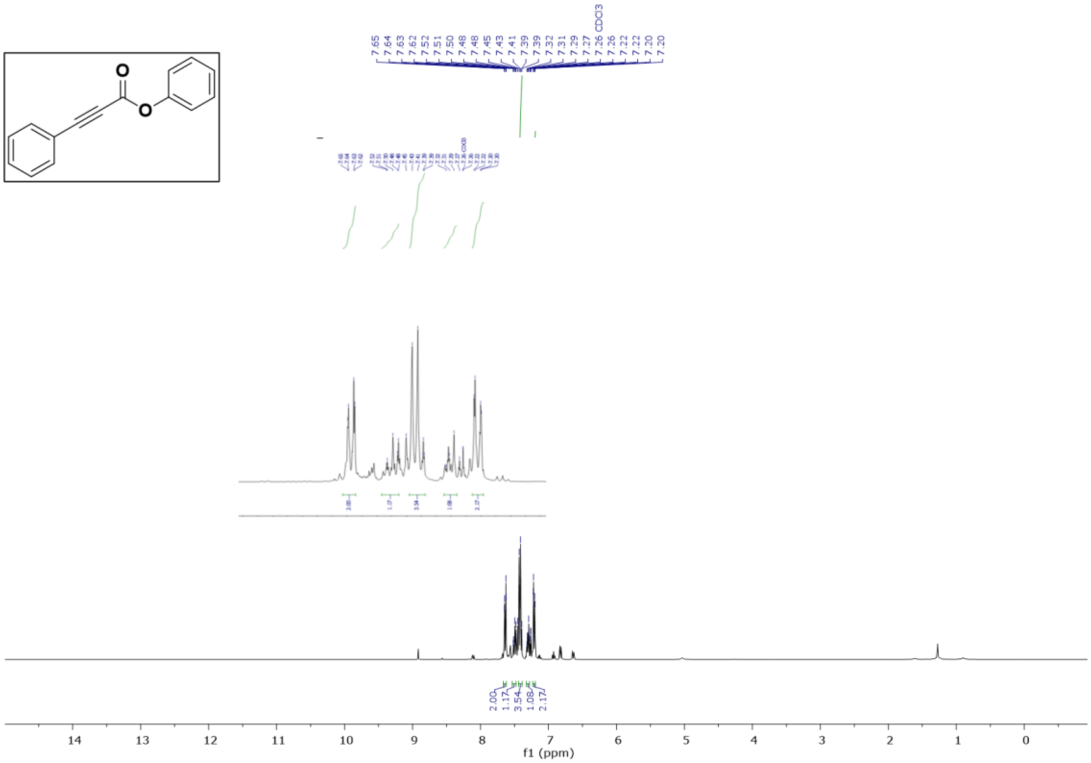

**Figure S39.**[1]H NMR **of phenyl 3-phenylpropiolate (14)** in $CDCl_3$

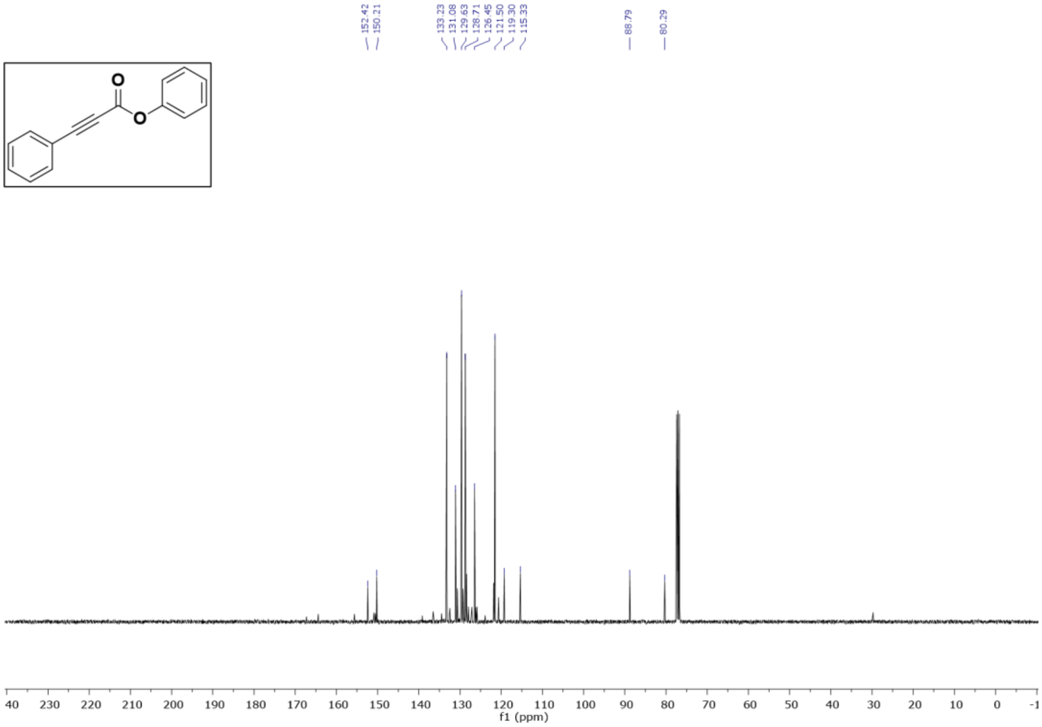

**Figure S40.**[13]C NMR **of phenyl 3-phenylpropiolate (14)** in $CDCl_3$

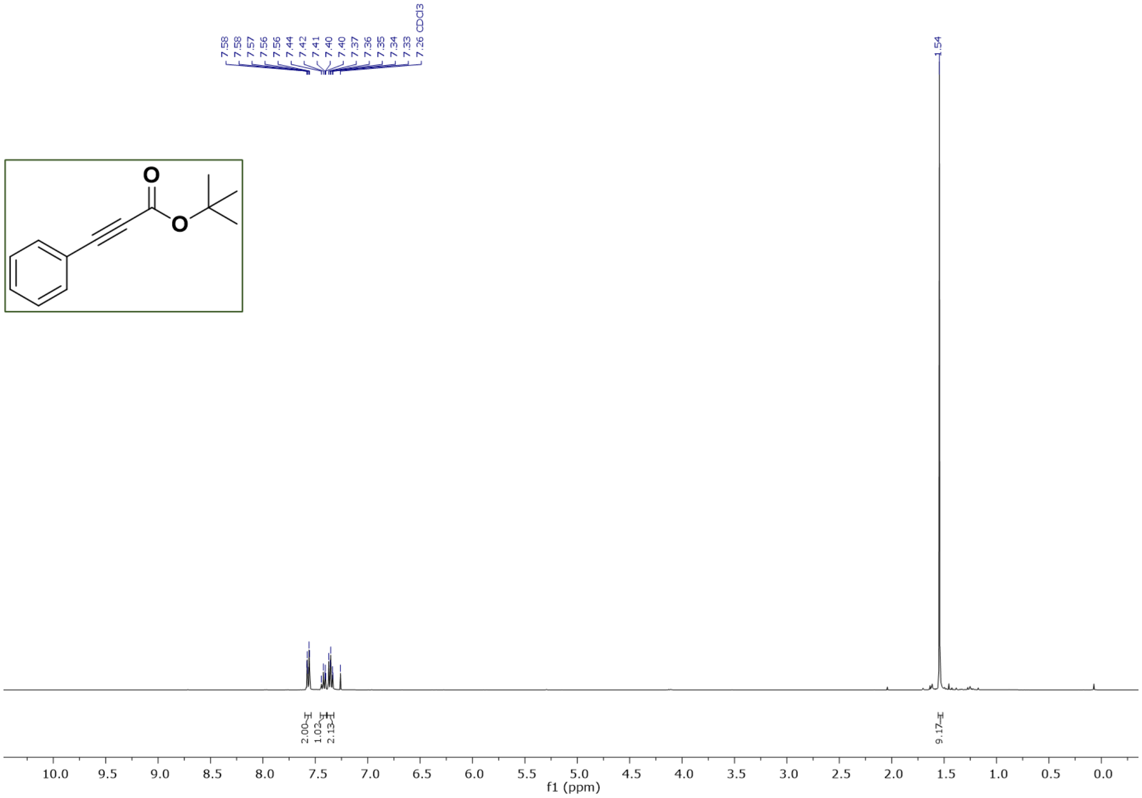

**Figure S41.** $^{1}$H NMR of **tert-butyl 3-phenylpropiolate (15)** in $CDCl_3$

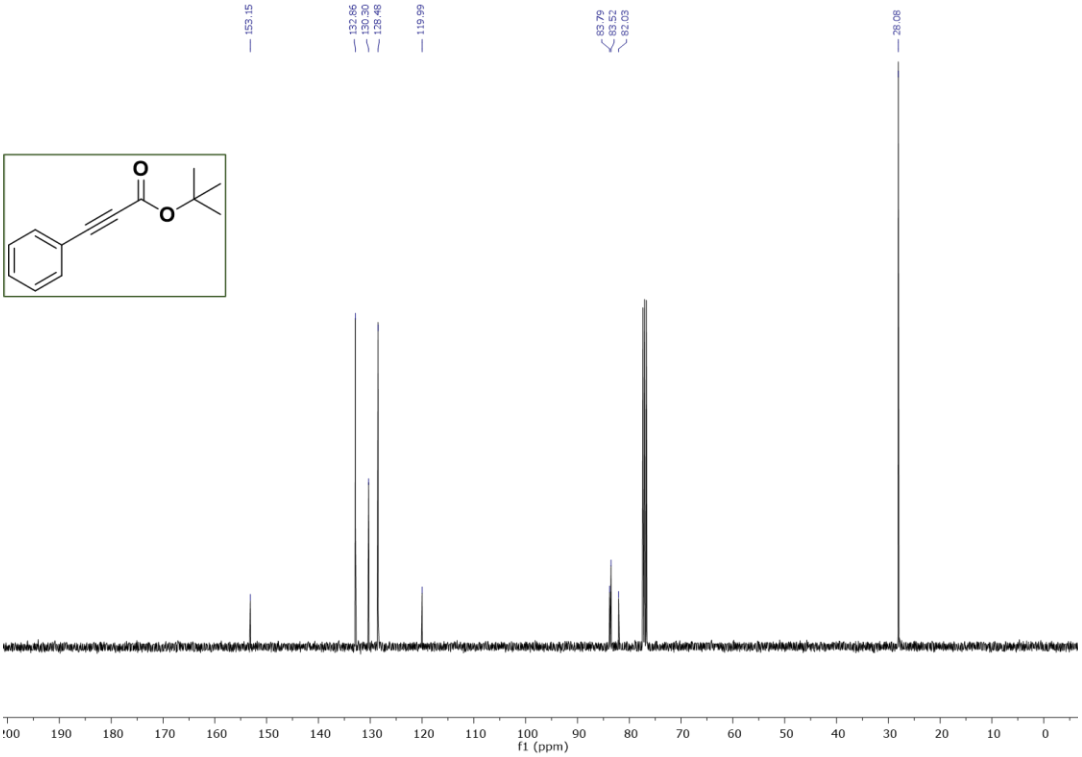

**Figure S42.** $^{13}$C NMR of **tert-butyl 3-phenylpropiolate (15)** in CDCl$_{3}$\

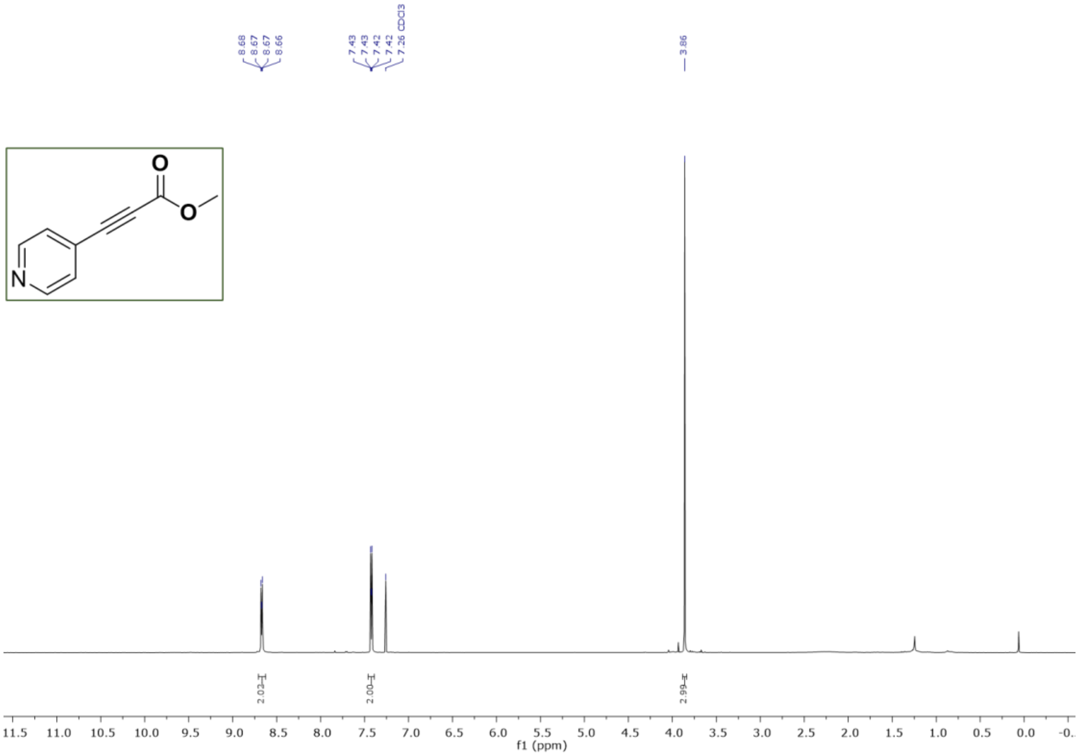

**Figure S43.** $^{1}$H NMR  of **methyl 3-(pyridin-4-yl)propiolate (16)** in $CDCl_3$

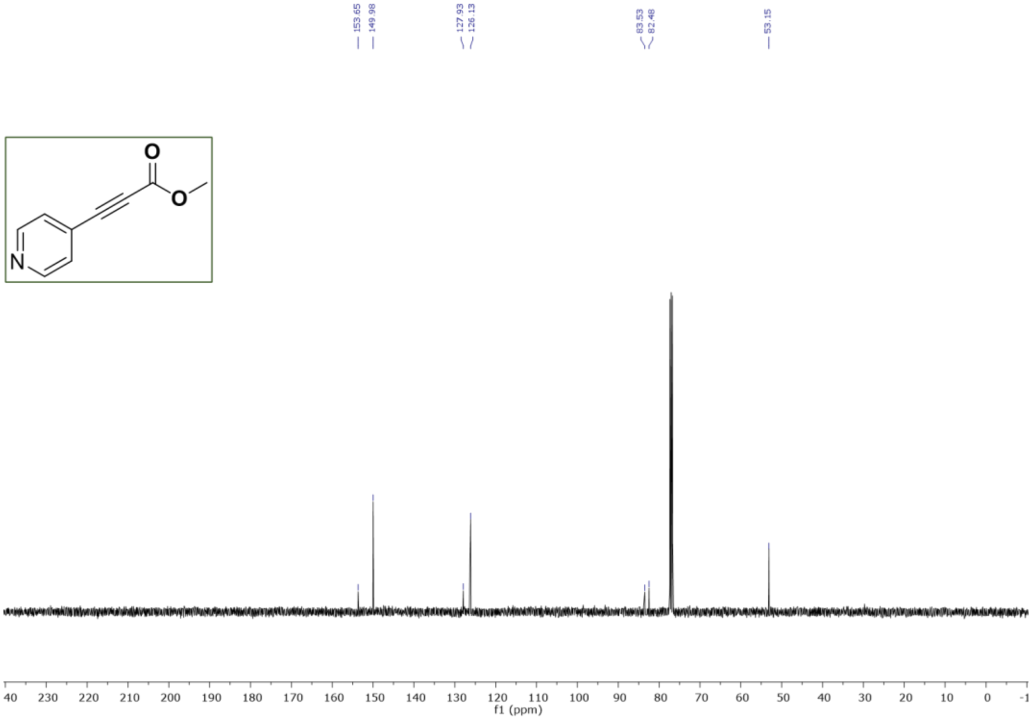

**Figure S44.** $^{13}$C NMR of **methyl 3-(pyridin-4-yl)propiolate (16)** in $CDCl_3$

**(F) References**

(1) Vickers, C. J.; González-Páez, G. E.; Wolan, D. W. Selective Detection and Inhibition of Active Caspase-3 in Cells with Optimized Peptides. *J. Am. Chem. Soc.* **2013**, *135* (34), 12869–12876. https://doi.org/10.1021/ja406399r.

(2) Weerapana, E.; Wang, C.; Simon, G. M.; Richter, F.; Khare, S.; Dillon, M. B. D.; Bachovchin, D. A.; Mowen, K.; Baker, D.; Cravatt, B. F. Quantitative Reactivity Profiling Predicts Functional Cysteines in Proteomes. *Nature* **2010**, *468* (7325), 790–795. https://doi.org/10.1038/nature09472.

(3) Weerapana, E.; Speers, A. E.; Cravatt, B. F. Tandem Orthogonal Proteolysis-Activity-Based Protein Profiling (TOP-ABPP)--a General Method for Mapping Sites of Probe Modification in Proteomes. *Nat. Protoc.* **2007**, *2* (6), 1414–1425. https://doi.org/10.1038/nprot.2007.194.

(4) Sievers, F.; Wilm, A.; Dineen, D.; Gibson, T. J.; Karplus, K.; Li, W.; Lopez, R.; McWilliam, H.; Remmert, M.; Söding, J.; Thompson, J. D.; Higgins, D. G. Fast, Scalable Generation of High-Quality Protein Multiple Sequence Alignments Using Clustal Omega. *Mol. Syst. Biol.* **2011**, *7*, 539. https://doi.org/10.1038/msb.2011.75.

(5) Goujon, M.; McWilliam, H.; Li, W.; Valentin, F.; Squizzato, S.; Paern, J.; Lopez, R. A New Bioinformatics Analysis Tools Framework at EMBL–EBI. *Nucleic Acids Res.* **2010**, *38* (Web Server issue), W695–W699. https://doi.org/10.1093/nar/gkq313.

(6) Yang, J.; Yan, R.; Roy, A.; Xu, D.; Poisson, J.; Zhang, Y. The I-TASSER Suite: Protein Structure and Function Prediction. *Nat. Methods* **2015**, *12* (1), 7–8. https://doi.org/10.1038/nmeth.3213.

(7) Roy, A.; Kucukural, A.; Zhang, Y. I-TASSER: A Unified Platform for Automated Protein Structure and Function Prediction. *Nat. Protoc.* **2010**, *5* (4), 725–738. https://doi.org/10.1038/nprot.2010.5.

(8) Zhang, Y. I-TASSER Server for Protein 3D Structure Prediction. *BMC Bioinformatics* **2008**, *9* (1), 40. https://doi.org/10.1186/1471-2105-9-40.

(9) Yan, T.; Desai, H. S.; Boatner, L. M.; Yen, S. L.; Cao, J.; Palafox, M. F.; Jami-Alahmadi, Y.; Backus, K. M. SP3-FAIMS Chemoproteomics for High-Coverage Profiling of the Human Cysteinome**. *ChemBioChem* **2021**, *22* (10), 1841–1851. https://doi.org/10.1002/cbic.202000870.

(10) Backus, K. M.; Correia, B. E.; Lum, K. M.; Forli, S.; Horning, B. D.; González-Páez, G. E.; Chatterjee, S.; Lanning, B. R.; Teijaro, J. R.; Olson, A. J.; Wolan, D. W.; Cravatt, B. F. Proteome-Wide Covalent Ligand Discovery in Native Biological Systems. *Nature* **2016**, *534* (7608), 570–574. https://doi.org/10.1038/nature18002.

(11) Wang, Q.; Chan, T. R.; Hilgraf, R.; Fokin, V. V.; Sharpless, K. B.; Finn, M. G. Bioconjugation by Copper(I)-Catalyzed Azide-Alkyne [3 + 2] Cycloaddition. *J. Am. Chem. Soc.* **2003**, *125* (11), 3192–3193. https://doi.org/10.1021/ja021381e.

(12) Kong, A. T.; Leprevost, F. V.; Avtonomov, D. M.; Mellacheruvu, D.; Nesvizhskii, A. I. MSFragger: Ultrafast and Comprehensive Peptide Identification in Shotgun Proteomics. *Nat. Methods* **2017**, *14* (5), 513–520. https://doi.org/10.1038/nmeth.4256.

(13) Teo, G. C.; Polasky, D. A.; Yu, F.; Nesvizhskii, A. I. Fast Deisotoping Algorithm and Its Implementation in the MSFragger Search Engine. *J. Proteome Res.* **2021**, *20* (1), 498–505. https://doi.org/10.1021/acs.jproteome.0c00544.

(14) Yu, F.; Haynes, S. E.; Nesvizhskii, A. I. IonQuant Enables Accurate and Sensitive Label-Free Quantification With FDR-Controlled Match-Between-Runs. *Mol. Cell. Proteomics MCP* **2021**, *20*, 100077. https://doi.org/10.1016/j.mcpro.2021.100077.

(15) Shteynberg, D. D.; Deutsch, E. W.; Campbell, D. S.; Hoopmann, M. R.; Kusebauch, U.; Lee, D.; Mendoza, L.; Midha, M. K.; Sun, Z.; Whetton, A. D.; Moritz, R. L. PTMProphet: Fast and Accurate Mass Modification Localization for the Trans-Proteomic Pipeline. *J. Proteome Res.* **2019**, *18* (12), 4262–4272. https://doi.org/10.1021/acs.jproteome.9b00205.
